# Molecular basis for multidrug efflux by an anaerobic RND transporter

**DOI:** 10.1101/2025.04.04.646765

**Authors:** Ryan Lawrence, Mohd Athar, Muhammad R. Uddin, Christopher Adams, Joana S. Sousa, Oliver Durrant, Sophie Lellman, Lucy Sutton, C. William Keevil, Nisha Patel, Christine Prosser, David McMillan, Helen I. Zgurskaya, Attilio V. Vargiu, Zainab Ahdash, Eamonn Reading

**Affiliations:** School of Biological Sciences, University of Southampton, Life Sciences Building, Highfield Campus, SO17 1BJ, Southampton, UK; Department of Chemistry, Britannia House, 7 Trinity Street, King’s College London, London, SE1 1DB, UK; Department of Physics, University of Cagliari, Cittadella Universitaria, S.P. Monserrato-Sestu, 09042, Monserrato, (CA), Italy; Department of Chemistry and Biochemistry, University of Oklahoma, 101 Stephenson Parkway, Norman, OK, 73019, USA; Department of Protein Structure and Biophysics, UCB Biopharma, Slough, SL1 3WE

## Abstract

Bacteria can resist antibiotics and toxic substances within demanding ecological settings, such as low oxygen, extreme acid, and during nutrient starvation. MdtEF, a proton motive force-driven efflux pump from the resistance-nodulation-cell division (RND) superfamily, is upregulated in these conditions but its molecular mechanism is unknown. Here, we report cryo-electron microscopy structures of Escherichia coli multidrug transporter MdtF within native-lipid nanodiscs, including a single-point mutant with an altered multidrug phenotype and associated substrate-bound form. We reveal that drug binding domain and channel conformational plasticity likely governs promiscuous substrate specificity, analogous to its closely related, constitutively expressed counterpart, AcrB. Whereas we discover distinct transmembrane state transitions within MdtF, which create a more engaged proton relay network, altered drug transport allostery and an acid-responsive increase in efflux efficiency. Physiologically, this provides means of xenobiotic and metabolite disposal within remodelled cell membranes that presage encounters with acid stresses, as endured in the gastrointestinal tract.

## Introduction

Presenting as an increasing global health challenge that demands immediate intervention, bacterial antimicrobial resistance (AMR) was attributed to approximately 1.14 million deaths in 2021^1^. Bacteria can acquire resistance through multiple mechanisms, however, one of the most pertinent is the overexpression of efflux pumps. The Resistance-Nodulation-Division (RND) family of efflux transporters comprises the most clinically relevant efflux pumps in Gram-negative bacteria^2,3^. These large protein conduits span their double membrane as a tripartite assembly comprising an inner-membrane RND transporter, an outer-membrane channel, and a periplasmic membrane fusion protein which connects the other two, forming a tunnel through which substrates are translocated to the external environment^4,5^ (**Fig. 1A**). Energised by the proton motive force (PMF), the polytopic RND proteins export an extensive range of chemically and structurally dissimilar antibiotics, thereby conferring multidrug resistance (MDR) in clinical isolates^4^.

**Fig. 1.**
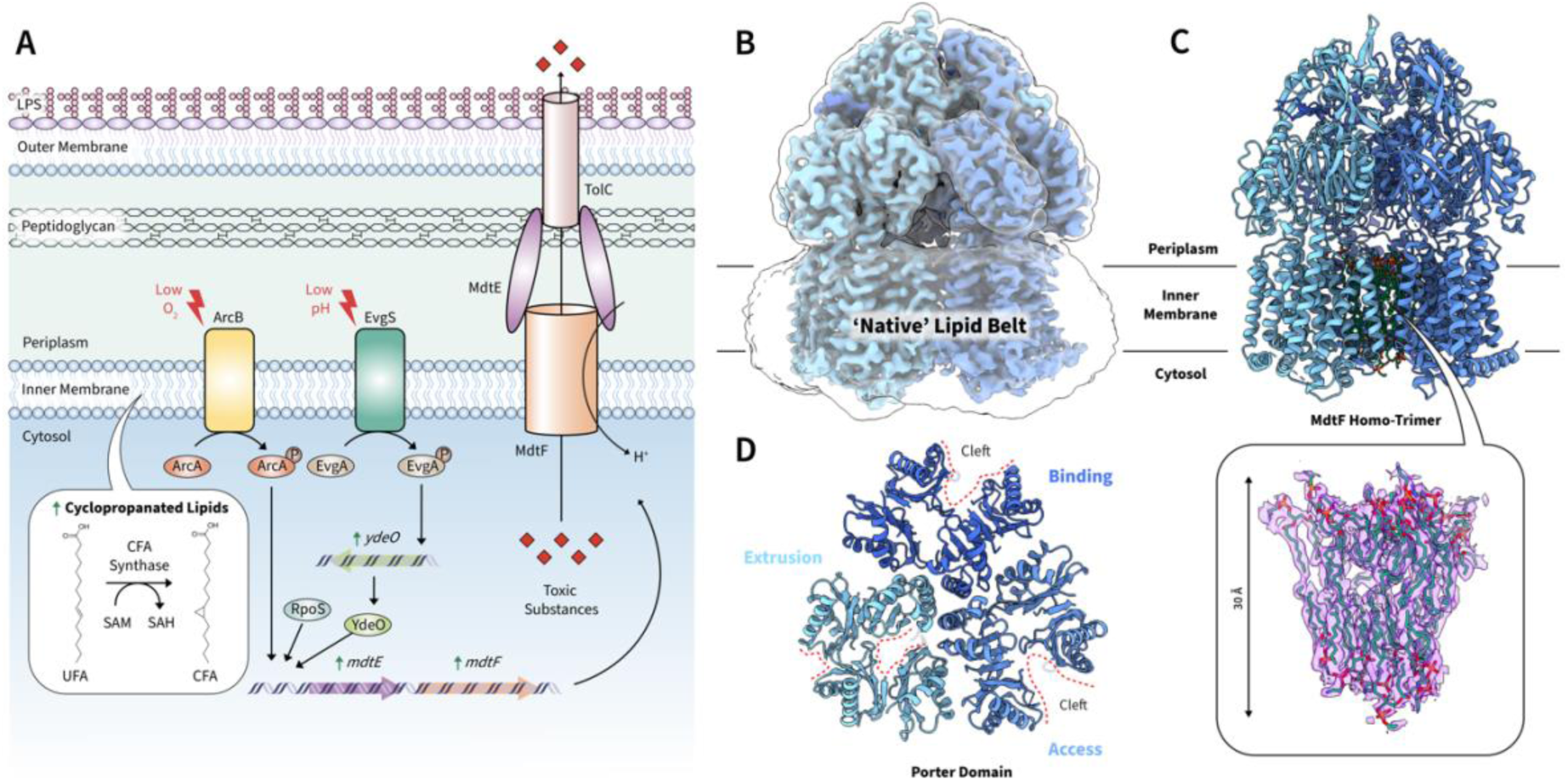
MdtF structure within a native-lipid environment. **a,** MdtEF is regulated by the two-component system EvgAS, a signal transduction system that confers acid resistance to E. coli. mdtF expression was also purported to be controlled by the conserved alternative sigma factor RpoS, required for E. coli survival in stress conditions associated with starvation, such as osmotic challenges, heat shock, and oxidative stress. In addition, the upregulation of MdtF, accompanied by increased efflux activity and drug tolerance, is observed in anaerobically grown E. coli. During this condition, the upregulation of MdtF is induced by the global anaerobic regulatory factor ArcA which controls the expression of genes related to the shift in the mode of energy metabolism from aerobic to anaerobic. MdtEF is also upregulated during stationary phase growth, which is accompanied by an increase in cyclopropanated lipid, whereby unsaturated fatty acids of the inner membrane are converted cyclopropane fatty acids. Here, a methylene group, deriving from S-adenosyl-L-methionine (SAM) is transferred to an unsaturated fatty acid chain in a reaction catalysed by cyclopropane fatty acid synthase. **b,** Structure of apo-MdtF^WT^ purified in SMALPs solved by cryo-EM to a resolution of 3.56 Å, demonstrating the maintenance of the native lipid belt encompassing the transmembrane domain of the protein in the electron density map. **c,** The ordered lipids buttressing the transmembrane domain of the three protomers of MdtF can be observed as elongated density in the cryo-EM structure. These were modelled in as PE lipids due to their predominance within E. coli membranes and as determined with our lipidomics experiments (**Supplementary Fig. 2**). **d,** Top view of the periplasmic porter domain, demonstrating its asymmetric structure as the open clefts are observed within the access and binding states. SAM: S-adenosyl-L-methionine, SAH: S-adenosylmethionine, CFA: Cyclopropanated fatty acid.

*Escherichia coli*, and closely related *Shigella* species, are facultative anaerobes that can cause severe infections in humans and exhibit varying susceptibilities to antibiotic treatments dependent on the oxygen levels within the environment they occupy^6^. Here, the constitutive AcrAB-TolC efflux pump from *E. coli* provides intrinsic resistance and remains the archetypal example of RND pumps. Extensive research on this system has facilitated a more precise structural comprehension of the functional mechanism of efflux adopted by this family^7–9^. However, during anaerobic conditions an alternative RND pump, MdtEF-TolC, is significantly upregulated and becomes the dominant efflux system (**Fig. 1A**) – being linked to the secretion of chemically diverse antibiotics, as well as expulsion of host-derived toxic nitrosyl indole derivatives as bacteria respire nitrate instead of oxygen^10^. Coincidently, the membrane is remodelled during hypoxic stress, harbouring an enrichment of cyclopropanated lipids, which exerts protective effects against extreme temperatures, low pH, and other environmental perturbations by decreasing permeability and increasing membrane rigidity^11–13^. This remodelled envelope seemingly being the MdtEF-TolC natural milieu. Indeed, *mdtEF* is also upregulated in other stress conditions where membrane remodelling also forms a mutual role in bacterial protection, such as nutrient starvation (i.e., transfer into stationary phase and biofilms), extreme acid, low pH, iron starvation, and during ‘catabolite induction’^10,14–19^.

Although these pertinent mechanisms highlight the clinical and physiological significance of this pump, molecular details which describe substrate efflux by MdtEF-TolC (formerly known as YhiUV-TolC^20^) is lacking. Multidrug recognition and export are controlled and powered by the integral membrane protein MdtF^20^ but the structural and mechanistic elements that define its condition-dependent activity and substrate selectivity and export are yet to be captured. In this article, we report three cryogenic-electron microscopy (cryo-EM) structures of MdtF, including substrate-free (apo-MdtF^WT^ and apo-MdtF^V610F^) and substrate-bound (R6G-MdtF^V610F^) states. In combination with molecular dynamics and functional assays, we uncover that MdtF adopts transmembrane conformational changes which provide an acid-responsive transport mechanism not found in other RND efflux pumps. This results in moderate efflux under neutral conditions which may be important in maintenance of PMF within stress conditions. But, when further acid-related stress is presented then this efflux pump can respond by increasing its efflux efficiency to battle xenobiotic attack and host-toxin clearance. This is important when we consider that anaerobic and acidic microhabitats are environmental signatures of the mammalian gut, where these bacteria colonise and infect, and that low pH amplifies the uptake of membrane-permeant weak acids that can impair cell growth in various ways and can reverse antibiotic resistance^21^.

Furthermore, we also reveal that the single-point mutation V610F, induced by antibiotic pressure in aerobic culturing, causes the swelling of the distal binding pocket (DBP), which reverberates allosteric changes to the entire protomer to make it more ‘AcrB-like’, and which likely determines its altered multidrug efflux activity. This adds to growing evidence that large-scale allosteric movements are the determinants of RND efflux pump specificity, and that select mutations can drastically affect its functional rotation mechanism to enact differing resistance profiles to antibiotics^22^.

Our study provides the first mechanistic view of how MdtF can recognise a wide range of harmful cellular and xenobiotic substances, including antibiotics, out of bacteria. These structures reveal molecular adaptations to efflux and survival within acidic and anaerobic-based stress. Together, these results provide a basis for future efflux pump inhibitors (EPIs) design, with the hope of restoring the activity of ineffective antibiotics^23,24^.

## Results

### The anaerobic RND-pump is conformationally distinct to its aerobic counterpart

The structure of *E. coli* MdtF^WT^ was determined by single-particle cryo-EM to a global resolution of 3.56 Å (**Fig. 1B**). To ensure that only the MdtF efflux pump was overexpressed, an *E. coli* strain deficient in the constitutively expressed native AcrB (C43Δ*acrAB*(DE3)) was used. MdtF is primarily upregulated in bacterial cells that have entered stationary phase and/or sessile biofilms; the composition of the lipidome within these states is drastically different to that of the exponential phase, with cyclopropanated acyl chains dominating the membranous environment^25–27^ (**Fig. 1A**). To obtain structures in a ‘native’ lipid environment and retain these lipids which may be essential in protein functionality and stabilisation, MdtF was extracted in SMA lipid particles (SMALPs) from stationary phase bacteria^28–32^. These polymer nanodiscs encapsulate membrane proteins within the intrinsic lipid mix of cells and are assembled via spontaneous portioning of the polymer into the bilayer. Native nanoparticles of *E. coli* MdtF^WT^ were purified to homogeneity before cryo-EM structural determination (**Supplementary Fig. 1**). Lipidomic analyses revealed the presence of phosphatidylethanolamine (PE), phosphatidylglycerol (PG), and cardiolipin (CL) phospholipids within the nanodisc sample. The fatty acid content was determined to comprise ∼ 50, 26 and 24 % saturated, unsaturated and cyclopropanated (SFA, UFA, and CFA), respectively (**Supplementary Fig. 2**). Cryo-EM data processing and analysis and an assessment of cryo-EM map quality is delineated in ***Supplementary Figs. 3-8*** *and* ***Table 1***.

To avoid the loss of lipid-bilayer structural information, the map reconstruction was performed with C1 symmetry. Here, we observe density for several lipids encompassing and buttressing the transmembrane region of MdtF in our structures, with those occupying the central cavity of the transmembrane region being better resolved. As PE was the predominant lipid type identified we, therefore, chose to model these ligands in the elongated density corresponding to the lipid molecules (**Fig. 1C**). In the apo-MdtF^WT^ structure we were able to resolve 21 lipid molecules and an additional 7 alkyl chains which could be from other lipid molecules within the central cavity. The bilayer is approximately 30 Å thick and exists as two triangular bilayer leaflet arrangements rotated relative to one another, with the inner leaflet being somewhat more organised than the outer leaflet chains, a phenomenon also found in AcrB^28^. Due to proximity and lack of regular arrangement between each monomer, this could indicate the importance of the lipid region in MdtF functionality and further highlights the amenability of this native purification system in membrane protein-based study.

Although the anaerobically upregulated MdtF shares high sequence similarity (∼71 %, **Supplementary Fig. 9**) and substrate profile to the aerobic, constitutively expressed AcrB pump, the residue differences are not localised to certain areas in terms of both sequence and 3D space (**Supplementary Fig. 10**). This indicates that MdtF must exhibit a unique functional purpose conferred by its three-dimensional structure. Therefore, we posit that a focused comparison between them is necessary to understand MdtF functional underpinnings. An important feature to recognise within RND-type efflux pumps is whether there is an existence of three functionally interdependent monomers, which could represent protomer states sequentially cycling through three conformations, i.e., access (or loose), binding (or tight), and extrusion (or open) (hereafter A, B, and E, respectively), during substrate translocation^9^. We observe asymmetric states between individual protomers of MdtF supportive of a functional cooperative mechanism analogous to AcrB, suggesting a conservation of this mechanism (**Fig. 1D** and **Supplementary Fig. 11**). This asymmetry is not captured by AlphaFold^33^ predicted structures (**Supplementary Fig. 12**), indicating the necessity of our experimentally derived atomic models.

MdtF can also be separated into analogous domains, loops, and channels to AcrB (**Fig. 2A** and **Supplementary Fig. 13A**). Interestingly, assignment of protomers in accordance with AcrB-related asymmetry revealed molecular dissimilarities in the MdtF access protomer especially (**Fig. 2B**), which is significant due to the implication of this state in ligand recognition and translocation. Here, adopting a different access state may suggest a different mechanism of initial substrate binding and lead to an altered broad-spectrum of substrate preferences between AcrB and MdtF. The funnel domain remains structurally conserved across all three protomers, exhibiting no significant asymmetry (**Supplementary Fig. 13B**). However, it is within the transmembrane domains where the structures begin to deviate more prominently, which we describe below.

**Fig. 2.**
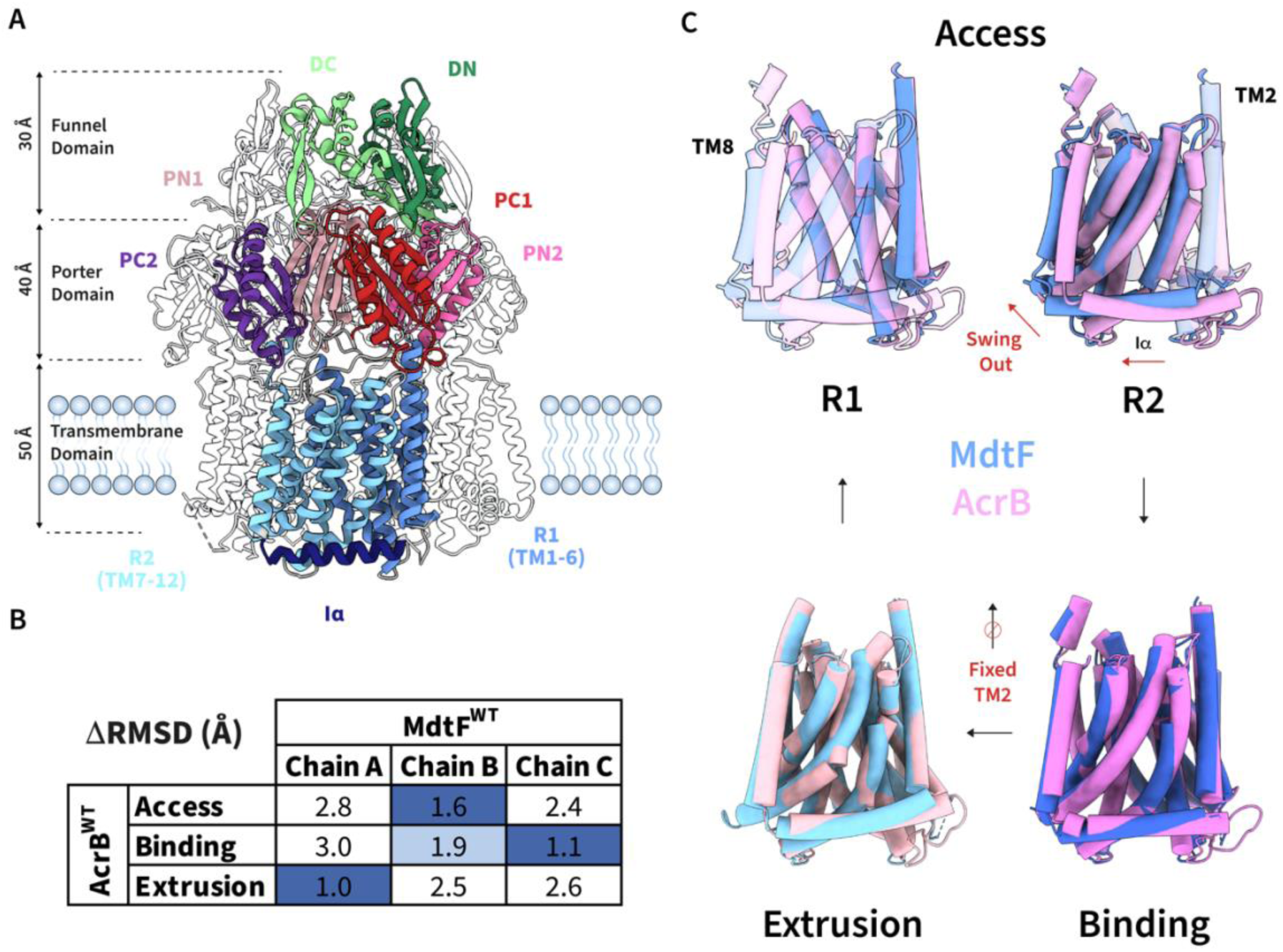
Altered conformations of MdtF^WT^ access state. **a,** Domains and subdomains of MdtF which are essential in its function, analogous to those observed within AcrB. **b,** Each chain was assigned to the access, binding, or access protomer state according to ΔRMSD measurements compared to the access, binding, and extrusion states observed within AcrB (PDB: 4DX5). **c,** Alignment of AcrB (pink) and MdtF^WT^ (blue) transmembrane domains which reveal structural differences between its helical arrangement as it cycles through the protomeric states. RMSD: Root mean square deviation.

In the transition between access and binding states of AcrB, the PC1 and PN2 subdomains undergo a structural change which causes the entire R2 domain (TM7-12) to swing away due to a pushing motion placed on the Iα helix arising from the TM2 downward displacement. Ligand binding to this state likely triggers the conformational change *in vivo*^34^. The swinging motion is mediated by a hinge point formed between engaged proton relay residues, D407, D408, and K940 in AcrB (K938 in MdtF). This ‘swing out’ mechanism is important in enabling proton translocation from cytoplasmic and periplasmic sides of the membrane, which powers substrate transport.

Strikingly, within our MdtF structure the identified access state already possesses a binding-like R2 state whereby it appears to be structurally ‘swung out’ (**Fig. 2C**). The TMD is flanked by two transmembrane helices, TM2 and TM8, which provide passive transmission of conformational energy to the porter domain, and vice versa, in AcrB^34,35^. Where the C-terminal end of TM8 harbours the essential ‘hoisting loop’ which confers its structural movements to the porter domain, coupling its dynamics to allosteric transmembrane domain transitions and functioning as a flexible hinge^35^. However, in MdtF^WT^, the TM2 helix remains ‘stationary’ through its binding to extrusion transition, moving only ∼0.3 Å in the z-direction of the membrane plane compared to ∼2.5 Å in AcrB (**Fig. 2C** and **Supplementary Fig. 14**). This is also corroborated by the lack of conformational translocation of the Iα helix during the transition between access and binding states (**Supplementary Fig. 15**). Whereas alterations in the hoisting loop from flexible to helical in structure remain within both asymmetric functional rotations of MdtF (**Supplementary Fig. 14**) and AcrB. Collectively, this suggests that MdtF adopts similar porter domain transitions to AcrB to elicit its broad substrate efflux but without requiring as large structural movements in its transmembrane domain between each protomeric state to do so.

### Drug-binding domain plasticity directs MdtF drug-specificity

Available, and allowable, drug channel and binding pocket conformations within RND efflux pumps act to define their promiscuous substrate efflux profiles^36^. Both residue patterning of substrate channels and entrances and conformational coupling have key roles in substrate selectivity across other homologs (**Supplementary Discussion** and **Figs. 28-31**)^9,34^. To better understand the conformational determinants in MdtF we solved the cryo-EM structure of a single-point mutation (Val610Phe) with an altered multidrug substrate profile^37^ at a resolution of 3.28 Å, following the same procedure used for MdtF^WT^. The single-point mutation conferred increased resistance to linezolid and tetracycline but reduced tolerance to azithromycin, telithromycin, and the efflux pump inhibitor, phenylalanine-arginine β-naphthylamide (PAβN)^37^.

The V610F conversion engenders a deepened hydrophobic pocket within the DBP (**Fig. 3A**). To explore this affected pocket volume and better understand the mechanism of substrate binding and translocation, we aimed to solve a substrate-bound structure of MdtF^WT^ and MdtF^V610F^ with the known planar aromatic cation (PAC) substrate Rhodamine 6G (R6G). Using a fluorescence polarisation assay^38^ we demonstrated that the purified protein is ligand recognition competent with similar binding affinity of the fluorescent R6G to MdtF^WT^ and MdtF^V610F^ (k_D_ ∼0.4-0.7 μM, **Fig. 3B**). Supporting thermal melting biophysical techniques, including circular dichroism (CD), and differential scanning calorimetry and fluorimetry (DSC and DSF) showed that R6G binding had a similar stabilising effect on the periplasmic drug-binding domain within both constructs (**Supplementary Fig. 16**). Although equilibrated with the same R6G concentration there was no observable, unambiguous density corresponding to R6G in the binding pocket of MdtF^WT^, we were able to solve the structure of R6G-MdtF^V610F^ at a resolution of 3.20 Å (**Fig. 3C**). R6G within our bound structure exclusively binds through hydrophobic interactions and is stabilised by cation-π (R6G-Phe626) and π-π (R6G-Phe178) interactions (**Fig. 3C**). A subtle rotation of the Phe626 ring accomodates R6G binding within the pocket (**Fig. 3D**). The higher polarisation signal observed for R6G bound to MdtF^V610F^ represents a binding mode that is more structurally restricted or homogenous which could rationalise the absence of unambiguous ligand density in our dataset for R6G-MdtF^WT^. This notion is reinforced by the observation of the contracted pocket volume in MdtF^WT^ which is penalised by steric clashes arising from the reorientation of phenylalanine rings F613, F626, F610, and Y327 which likely impede R6G ligand ordering within the pocket (**Fig. 3D**).

**Fig. 3.**
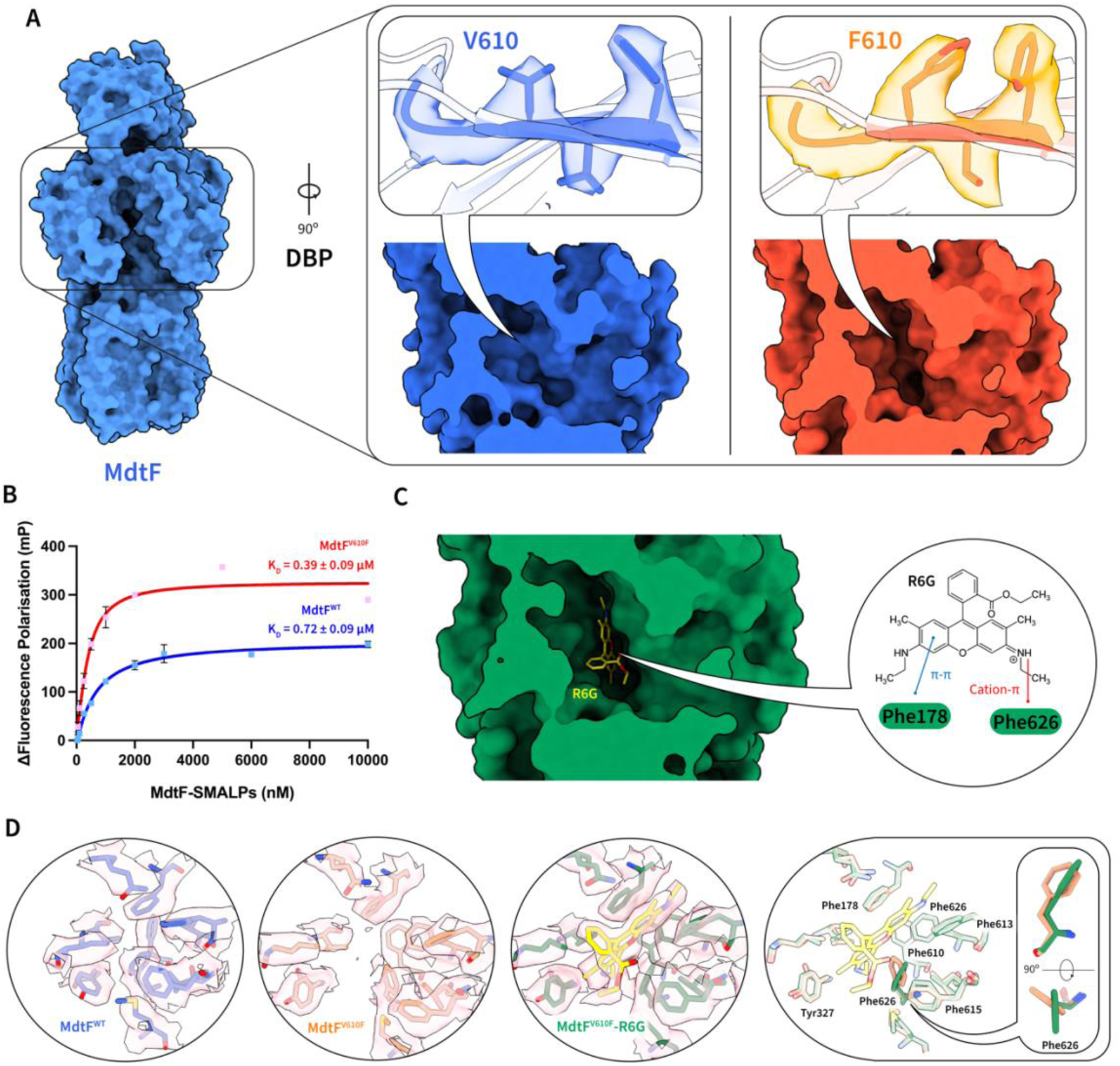
MdtF^V610F^ structure creates a hydrophobic nook within the DBP. **a,** Structure of apo-MdtF^V610F^ solved by cryo-EM at a resolution of 3.28 Å with an observed formation of a ‘hydrophobic nook’ within the DBP, arising due to the single-point mutation. **b,** Binding of R6G by MdtF^WT^ and MdtF^V610F^ as determined by a fluorescence polarisation assay. R6G was maintained at 1 mM throughout and its emission wavelength was 550 nm. The binding isotherms demonstrate a K_D_ of 0.39 and 0.72 nm for MdtF^WT^ and MdtF^V610F^, respectively, in a buffer containing 50 mM sodium phosphate, 150 mM NaCl, 10 % glycerol. Reported data are the average and standard deviation from independent measurements (n = 3) and were fitted to a hyperbola function (FP = (Bmax * [protein]) / (K_D_ + [protein])). **c,** Structure of R6G-MdtF^R6G^ solved by cryo-EM at a resolution of 3.20 Å, demonstrating cation-p (R6G-Phe626) and p-p (R6G-Phe178) interactions with R6G in the binding pocket of MdtF. **d,** MdtF^V610F^ demonstrates a more open cleft within the binding pocket compared to MdtF^WT^. R6G binding is facilitated by a reorientation of the Phe626 residue. DBP: Distal binding pocket, FP: Fluorescence polarisation, R6G: Rhodamine 6G.

Collectively, these results suggest that the V610F mutation creates altered ligand recognition profiles. This could result from the combination of two factors: i) altered global conformational preferences of the MdtF monomers. Intriguingly, a similar mutation in AcrB^V612F^, analogous to MdtF^V610F^, was recently discovered to change the global conformation of AcrB towards a ‘binding-like’ state, eliciting a similar alteration to its MDR phenotype^22^. This supports the notion of a long-range structural change resulting from this single-point mutation; ii) local conformational changes around the mutation site, altering the interaction with MdtF substrates. Indeed, the mutation causes the emergence of a ‘hydrophobic nook’, possibly enabling MdtF to engage differently with its substrates.

Computational modelling was performed to provide further insights into these hypotheses. First, to better understand the conformational dynamics of MdtF in a membrane-like environment, we performed molecular dynamics (MD) simulations for MdtF^WT^ and MdtF^V610F^ (as well as AcrB to compare dynamical features of the two transporters) in a model bilayer. Our investigations reveal major conformational differences in both the porter and the transmembrane domains. In the porter domain, the PC1 and PC2 subdomains of the entrance cleft, which in AcrB define the channel 2 (CH2) and the access pocket mediating the uptake of several substrates^36,39^, displayed different preferred arrangements, as highlighted by the distribution of their pseudo-contacts (**Fig. 4**; see *Materials and Methods* for details). Compared to AcrB, MdtF^WT^ has an overall more closed cleft at equilibrium (**Fig. 4A**). In particular, the distributions of pseudo-contacts almost overlap between the access and binding states of MdtF^WT^ in contrast to AcrB, where the access state features a more open cleft (**Supplementary Fig. 17**), further supporting our classification of a ‘binding-like’ access state. In addition, there were more distinct intermediate and extreme conformations allowed by MdtF^WT^ in its access and binding states compared to AcrB (3 vs. 2 and 2 vs. 1 in the access and binding states, respectively, **Fig. 4B**). We next considered whether the creation of the hydrophobic nook induced by the V610F mutation also leads to long-range conformational changes within MdtF. The cryo-EM structure of MdtF^V610F^ reveals that the mutation partly restores the AcrB-like difference in the arrangement of PC1 and PC2 domains between the access and binding states, mainly arising from a shift in the distribution of the binding protomer. The extrusion state in MdtF^V610F^ is characterised by a wider cleft compared to MdtF^WT^ (**Fig. 4A**). Such a difference may partly be ascribed by the protrusion of residues A675 and S676 in the access state of MdtF^WT^, seen in the cryo-EM structure of the MdtF^WT^ transporter but not in MdtF^V610F^ (**Supplementary Fig. 18**). Interestingly, MD simulations reveal that the V610F mutation induces an overall opening of the cleft in all monomers, resulting in a somewhat more symmetric MdtF conformation compared to the wildtype transporter.

**Fig. 4.**
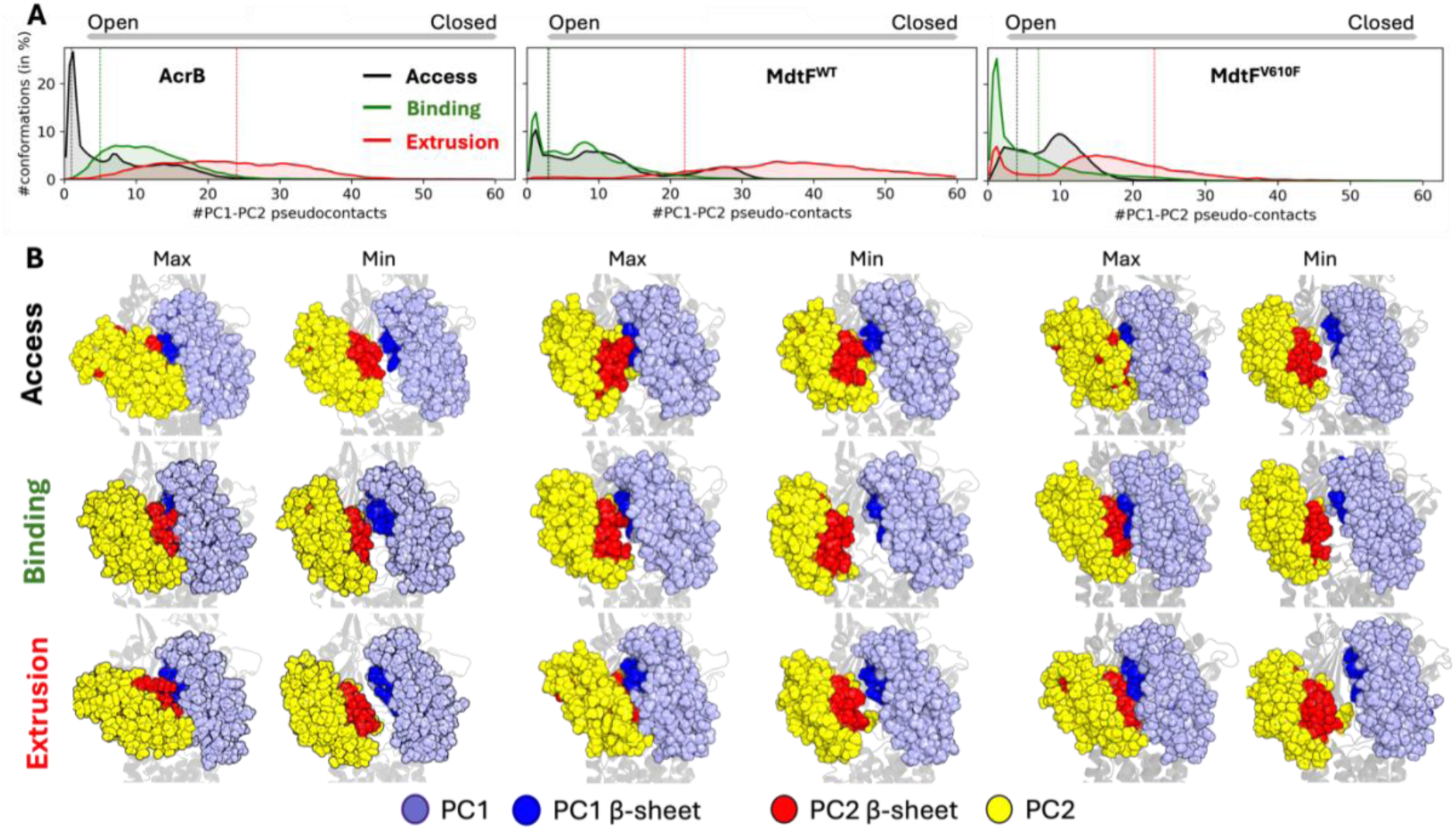
Molecular dynamics reveals PC1-PC2 domain transformations are similar between AcrB and MdtF^V610F^, but different in MdtF^WT^. **a,** Cumulative average of the native contacts at the PC1-PC2 cleft, calculated using a cut-off of 10 Å between C_a_ atoms and averaged from the simulation replicas of AcrB, MdtF^WT^ and MdtF^V610F^ (left to right). This plot represents the distribution of residue-residue native contacts throughout the simulation trajectory. Contacts in experimental reference structure are highlighted. Lesser contacts correspond to the cleft opening whereas higher number of contacts correspond to closing. **b,** Extreme conformations of the PC1-PC2 cleft observed in the MD simulations. PC2 is represented as yellow spheres, PC1 is shown in light blue, and the beta sheets of PC1 and PC2 are depicted in blue and red, respectively.

Next, we performed molecular docking of linezolid, ciprofloxacin, and PAβN on both MdtF^WT^ and MdtF^V610F^ to assess the impact of the mutation on possible direct interactions with substrates. Interestingly, there were very minor changes in the estimated affinities within the DBPs of MdtF^WT^ and MdtF^V610F^ (**Supplementary Table 3**). The largest change was the increase in affinity for ciprofloxacin within MdtF^V610F^, which is compatible with enhanced efflux compared to MdtF^WT^. Notably, the affinity of MdtF^WT^ to R6G and the natural nitrosyl indole substrate remained unchanged in comparison to MdtF^V610F^, suggesting their export would be unaffected in accordance with the fluorescence polarisation assay for R6G (**Fig. 3B**).

Further work exploring the structure-function relationships between MdtF and its known substrate classes will be required to fully explain how its conformations define its specificity. However, our data support the occurrence of a global conformational change resulting from a single-point mutation and the importance of this residue in determining substrate specificity in RND-based efflux pumps, in agreement with recent data for AcrB^22^. Taken together, these differences in substrate recognition and conformational preferences of MdtF (WT and V610F) may explain individual mechanisms of substrate binding, translocation and, therefore, efflux propensity.

### MdtF pump-identity is defined by its proton relay-related movements

The putative transmembrane proton relay network in MdtF (AcrB) comprises D407, D408, K938 (K940), and R969 (R971) (**Fig. 5A**). These residues are conserved both in terms of sequence and structural position and are likely essential for eliciting PMF-driven structural changes during efflux (**Supplementary Figs. 19 and 20**). To confirm the importance of the proton relay network in MdtF efflux, we employed a Nile Red efflux assay on a functional mutation of a proton relay residue, MdtF^D408A^. Here, *E. coli* Δ9-Pore cells lacking all nine TolC-dependent transporters were used to express plasmid-borne MdtEF. A decrease in fluorescence intensity is indicative of active efflux of the lipophilic dye Nile Red. In comparison to MdtEF^WT^, MdtEF^V610F^, and AcrAB^WT^, MdtEF^D408A^ eliminated efflux capability with no characteristic one-phase exponential decay observed (**Fig. 5B**), substantiating the importance of the proton relay network in MdtF function (**Supplementary Fig. 21**). Intriguingly, MdtEF^WT^ and MdtEF^V610F^ (t_efflux50%_ = ∼21 and ∼23 s, respectively) rate of Nile Red efflux was observed to be slower in comparison to AcrAB^WT^ (t_efflux50%_ = ∼9 s).

**Fig. 5.**
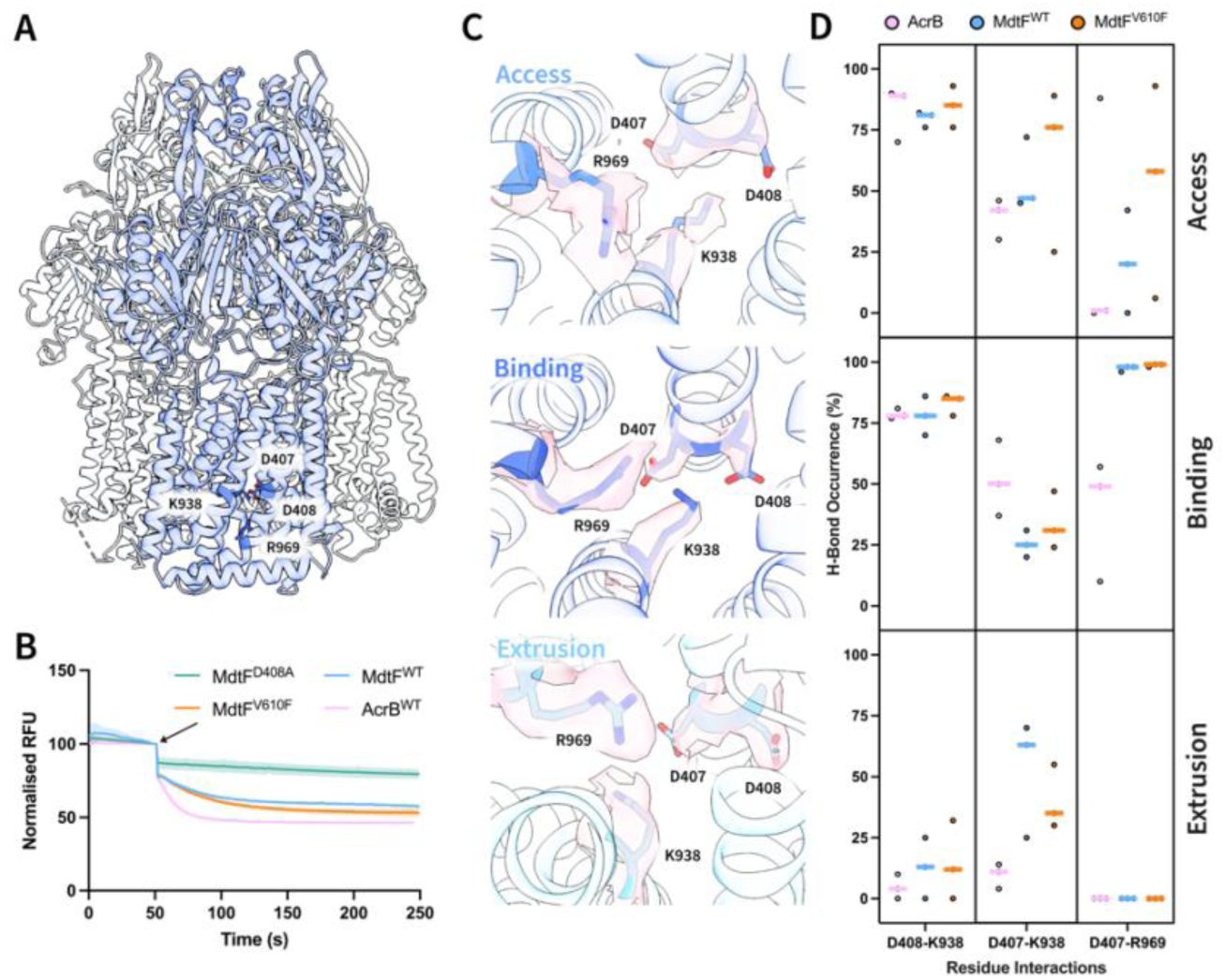
Engaged proton relay-network within binding and extrusion states within MdtF. **a,** Location of four essential proton relay residues (D407, D408, K938, and R969) within the transmembrane domain of RND transporters. **b,** Nile Red efflux assay demonstrates the reduced rate of MdtEF-mediated efflux compared to AcrAB and the MdtEF^D408A^ mutant renders the structure non-functional. E. coli Δ9-Pore cells harbouring p151A-AcrAB, pUC19-MdtEF^WT^, pUC19-MdtEF^V610F^, or pUC19-MdtEF^D408A^ plasmids were monitored for fluorescence decay at 650 nm for 200 s. Here, proton gradient-powered efflux pumps were restarted following the addition of 1 mM glucose after 50 s (indicated by the arrow). Reported data are the average and standard deviation from independent measurements (n = 3) and were fitted to a one-phase decay exponential function (Y = (Y_0_ - Plateau) * exp(-K * X) + Plateau). **c,** Proton relay residues of MdtF^WT^ between each of the access, binding, and extrusion states depicted in cartoon representation. The corresponding density for each residue is depicted in surface form (light pink). **d,** H-bond occurrence (in % over the simulation time) at the proton relay site between access, binding, and extrusion states from top to bottom. H-bonds were detected using 3.0 Å between acceptor and donor, and an angle smaller than 35° between H, donor, and acceptor (calculated using the cptraj module of Amber24). RFU: Relative Fluorescence Units.

We described earlier that the overall transitions and movements of transmembrane helices between the MdtF protomers are reduced compared to AcrB (**Fig. 2**). We propose that this could, therefore, direct the mechanism by which MdtF can utilise, and transfer, the energy stored in the transmembrane electrochemical gradient to enact conformational changes for drug translocation throughout the porter domain. Our cryo-EM structures represent static interpretations and do not account for dynamic residue movements. By analysing the proton relay residue interactions occurring along MD trajectories, we observed a small depletion in the engagement of D407-K938 in the binding state of MdtF^WT^ and MdtF^V610F^ compared to AcrB (**Fig. 5C-D**). This is compensated by a non-vanishingly engagement in the extrusion protomer, as seen in multiple independent simulations. Furthermore, MdtF features a relatively high occurrence of D407-R969 interactions in the access and binding states (**Fig. 5D**), which is absent in all AcrB protomers. It was proposed that R971 in this transporter could act as a conformational electrostatic switch coupled to the protonation state of D407 and D408^34^, while in our MD simulations we see a direct involvement of R969 (**Fig. 5D** and **Supplementary Fig. 22**).

Collectively, these results point to an engaged TM bundle in the extrusion state of MdtF, which may reduce proton exchange that could be the consequence of the altered conformational profile in MdtF. Notably, although our MD simulations reveal relatively large structural departures in the MdtF dynamics compared to the cryo-EM geometries, the simulations of AcrB provided findings that are in good agreement with experimental data^34^. This supports the accuracy of our computational results and the possibility of a truly distinct mechanism of coupling between proton translocation and substrate export in MdtF.

### MdtF drug transport is responsive to proton load

To better evaluate the effects of these reduced structural movements on MdtF-mediated efflux action, we performed accumulation assays with an increased range of substrates. We first determined whether the activity of MdtEF varies depending on the growth phase. When cells carrying an empty pUC19 vector or MdtEF^WT^ were collected in the mid-exponential or stationary phases, the accumulation of Hoechst 33342 (HT) was lower in cells producing MdtEF^WT^ than in cells with an empty vector. Thus, the pump is functional in both growing and stationary cells and can reduce the intracellular accumulation of HT.

Previous studies, however, showed that the outer membrane of *E. coli* and other bacteria is modified and becomes less permeable in the stationary non-growing cells^40^. Hence, such differences in the permeability barrier of the outer membrane could mask the differences in the efflux efficiency of MdtEF. Therefore, Δ9-Pore cells were induced with L-arabinose to produce a large non-specific Pore. Such hyperporination of the outer membrane eliminates its permeability barrier and enables a more specific assessment of efflux^41^. Surprisingly, we found that the exponentially growing and the stationary cells producing MdtEF differ in the accumulation of HT with the stationary cells accumulating significantly lower amount of HT than the exponential cells (**Fig. 6A** and **Supplementary Fig. 23A**). In contrast, cells lacking the pump accumulated similar levels of HT in growing and stationary cells. The levels of expression of MdtEF were similar in the compared cells (**Supplementary Fig. 23A**). However, the amounts of the Pore in the membrane fractions were notably lower in the stationary phase cells, suggesting that both the differences in the permeability of the outer membrane and differences in the activity of MdtF could contribute to the observed differences in HT accumulation in the stationary cells producing MdtF (**Supplementary Fig. 23B**).

**Fig. 6.**
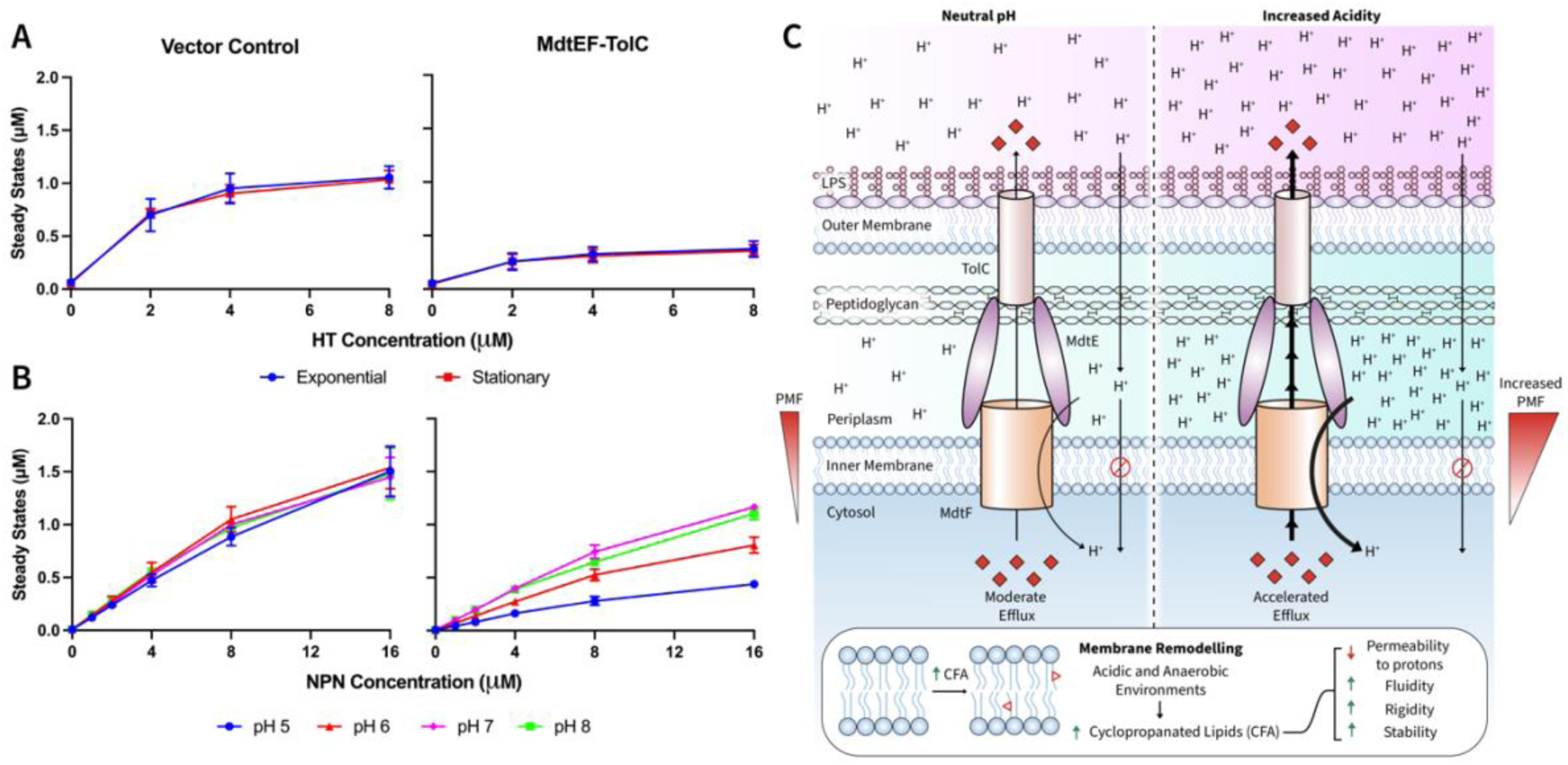
MdtEF-TolC functions within a remodelled membranous environment. **a,** Steady state accumulation levels of HT were measured in *E. coli* Δ9-pore cells, harbouring an empty vector or pUC19-MdtEF^WT^ plasmid, as a function of external HT concentration. Due to difference in pore expression at exponential and stationary phases, outer membrane pore expression was not induced (**Supplementary Fig. 23A**). Individual data points represent mean values from three independent measurements and error bars are indicative of the standard deviation (n = 3). **b,** Steady state accumulation levels of the fluorescent, non-ionisable probe NPN in *E. coli* Δ9-pore cells, harbouring an empty vector or pUC19-MdtEF^WT^ plasmid, as a function of external NPN concentration. Individual data points represent mean values from three independent measurements and error bars are indicative of the standard deviation (n = 3). **c,** Highly protonated environments within acidic and anaerobic conditions where MdtF is expressed results in an increased PMF, whereby its reduced rate of efflux may be essential to minimise proton entry and prevent bacterial acidification. Conditions of MdtF expression also indicate an altered membrane environment arising from increased cyclopropanation of inner membrane lipids. Here, decreased permeability to protons, alongside increased membrane fluidity, rigidity, and stability results in a remodelled membrane where MdtF exhibits an increased efflux efficiency which may be rationalised by its structural deviations from other RND homologs. CFA: Cyclopropanated fatty acid, HT: Hoechst 33342, NPN: *N*-Phenyl-1-naphthylamine, PMF: proton motive force.

We next analysed how the external pH affects the activity of MdtEF. Since HT is charged and its fluorescence is pH-sensitive (pK_a_: N_1_ 7.87, N_1_ 5.81, and N_3_ 5,02), in these experiments we used a non-ionisable membrane probe *N*-phenyl-naphthylamide (NPN). The exponential hyperporinated cells with and without MdtEF expression were resuspended in buffers with pH values decreasing from 8.0 to 5.0. We previously found that in this range of pH AcrAB retains similar NPN efflux activity^42^. In contrast, the activity of MdtEF was pH-dependent and was strongly enhanced at acidic pH (**Fig. 6B**). Interestingly, dynamic water exchange analysis demonstrated a conservation of the proton relay-based mechanism in MdtF (**Supplementary Fig. 24**). However, the V610F mutation restores a water accessibility pattern that closely resembles that of AcrB. Considering the environments which relate to the expression pattern of MdtF, these results signify the adaptation of its activity specifically to highly protonated environments.

We conclude that MdtF overexpression may facilitate bacterial survival because its proton translocation pathway is optimised to a different membrane composition. In accordance with this, the more rigid inner membrane within stationary phase of growth or under acidic stress arising from cyclopropanated lipid enrichment is more conducive for the activity of MdtEF (**Fig. 6C**). This indicates a potential adaption of MdtF to function within the hostile environments it occupies, transitioning to an overall survival mechanism with decreased membrane permeability to protons and a reduced, but acid-accelerated, rate of overall efflux.

## Discussion

*Escherichia spp.* bacteria encounter hostile environments during their enteric journey, such as oxygen deprivation and highly acidic conditions, within the mammalian gut. Here, bacterial membrane remodelling engenders a less permeable and more rigid environment; this milieu coincides with MdtEF-TolC overexpression patterns which ultimately effectuates bacterial survival within these stress conditions. Through combining cryo-EM structural information with molecular modelling predictions and functional assays, these findings collectively establish the structural basis for MdtF-mediated substrate efflux. A trapped transmembrane configuration between ‘access’ and ‘binding’ states in our MdtF model suggests a unique functional rotation mechanism of substrate translocation. This molecular architecture creates a moonlighting pump with moderate efflux under bacterial exhaustion conditions, where a PMF is putatively harder to maintain, but which becomes more efficient at acidic pH, when the membrane is hyperpolarised. Interestingly, previous studies have shown that *mdtEF* contributes to detoxification of acid induced metabolic by-products, but that it is not necessarily activated by the occurrence of acid stress. Even though it shares an operon with the central acid resistance regulator *gadE*. Instead, the induction of the *gadE-mdtEF* operon is thought to be rigged under many different circumstances that could presage an encounter with acid stresses^43^. Here, we demonstrate this phenomenon happening at the molecular level of MdtF pump action. This behaviour has not yet been identified in other RND pumps and rationalises the observed upregulation patterns of *mdtEF* in nutrient and oxygen starved bacterial populations, i.e., in stationary phase and biofilms^18,19^.

Here, we also resolve the molecular underpinnings for the altered MDR phenotype created by the V610F single-point mutant in MdtF, created through bacterial antibiotic pressure to levofloxacin^37^. The positioning of V610F in the drug binding pocket of MdtF creates a differing ligand recognition competent structure with the development of a ‘hydrophobic nook’. Interestingly, although MdtF^V610F^ retains similar TM protomer states (**Supplementary Fig. 25**) and slower rate of efflux, as found for MdtF^WT^, it has adapted its DBP volume and porter domain movements to be more like that of AcrB. Where modification to the porter domain alters its substrate recognition profile without significantly changing the mode of operation for powering translocation. This supports the growing evidence that broad recognition profile alterations can be achieved through small changes in the porter domains of RND efflux pumps, without greatly affecting their underlying functional capabilities^22^. This is a remarkable observation because of the high degree of allosteric conformational coupling between the TM and porter domains required for export function in RND transporters^34,44^.

More generally, our work provides a foundation for the enhanced design of effective efflux pump inhibitors (EPIs) – so-called resistance breakers – to restore and maintain antibiotic efficacy^45^. Especially as MdtF is the known RND pump to elicit a defined physiological role in anaerobic conditions, these results may serve as a paradigm for anaerobic efflux control by RND pumps across other bacterial species, such as the ESKAPE (*Enterococcus faecium*, *Staphylococcus aureus*, *Klebsiella pneumoniae*, *Acinetobacter baumannii*, *Pseudomonas aeruginosa*, and *Enterobacter spp.*) classes.

## Materials and Methods

### Reagents and Chemicals

Unless stated otherwise, all common reagents and chemicals were obtained from Sigma-Aldrich.

### Plasmids and Mutagenesis

*Construction of pET15b MdtF plasmids.* An overexpression plasmid containing the wildtype *mdtF* sequence with a C-terminal 6xHistidine tag (MdtF-6xHis) was constructed from a pET15b-MdtF-sGFP-6xHis plasmid^46^. In brief, the *sGFP* sequence was deleted by site-directed mutagenesis, according to the Q5® Site-Directed Mutagenesis Kit (New England Biolabs) using forward and reverse primers that flank the *sGFP* sequence.

Construction of pUC19-MdtEF plasmids. These plasmids, harbouring *mdtEF* and its native promoter, were constructed for bacterial susceptibility and Nile Red efflux assays. In brief, the pUC19 plasmid (Thermofisher) was linearised with PstI and BamHI restriction enzymes (New England Biolabs). The *mdtEF* gene, including the 180 base pairs upstream flanking sequence, was then amplified from *E. coli* genomic DNA (Zyagen) by PCR. The primers used incorporated a C-terminal 6 x Histidine tag for detection by anti-His immunoblotting. The amplicon was subsequently cloned into the linearised pUC19 vector using NEBuilder® HiFi DNA Assembly kit (New England Biolabs). For bacterial susceptibility and Nile Red efflux assays, the 6 x Histidine tag was deleted by site-directed mutagenesis, according to the Q5® Site-Directed Mutagenesis Kit (New England Biolabs) using forward and reverse primers that flank the 6 x Histidine sequence.

*Plasmid Mutagenesis.* For V610F mutagenesis, a single-point mutation was introduced in the pET15b-MdtF-6xHis plasmid according to the Q5® Site-Directed Mutagenesis Kit (New England Biolabs). Single-point mutations were also introduced into the pUC19-MdtEF plasmids through site-directed mutagenesis, according to the Q5® Site-Directed Mutagenesis Kit (New England Biolabs).

The final plasmid sequence within the aforementioned plasmids were confirmed by Sanger sequencing (Eurofins Genomics). Primers used are delineated in **Supplementary Table 2**.

### Overexpression and Purification of MdtF

*MdtF Overexpression in E. coli cells.* The pET15b-MdtF-6xHis plasmid, containing the sequence for MdtF^WT^ or MdtF^V610F^, was transformed into C43(DE3) Δ*acrAB E. coli* cells. 7 mL of an overnight LB culture was added to 1 L of pre-warmed LB culture containing 100 μg/mL ampicillin and grown at 37 °C until an OD_600_ of 0.6-0.8 was reached. The culture was incubated at 4 °C for 30 min and then induced with 1 mM IPTG and grown for 16-18 h at 18 °C. The cells were subsequently harvested by centrifugation at 4,200 x *g* and 4 °C for 30 min and washed with ice-cold phosphate buffer saline (PBS).

Cell pellets were immediately resuspended in Buffer A (50 mM sodium phosphate, 300 mM sodium chloride, pH 7.4), supplemented with 1 μL of Benzonase, 5 mM beta-mercaptoethanol (β-ME), 1 mM phenylmethylsulphonyl fluoride (PMSF), and a protease inhibitor tablet (Roche). The cell suspension was then passed twice through a microfluidizer processor at 25,000 psi and 4 °C. Insoluble material was removed by centrifugation at 20,000 x *g* and 4 °C for 30 min. Membranes were pelleted from the supernatant by centrifugation at 200,000 x *g* and 4 °C for 1 h. Membrane pellets were resuspended in 30 mL of ice-cold buffer B (50 mM sodium phosphate, 150 mM sodium chloride, 10 % (w/v) glycerol, pH 7.4), supplemented with 1 mM PMSF and a protease inhibitor tablet (Roche), and homogenised using a Potter-Elvehjem Teflon pestle and glass tube.

*Solubilisation and Purification in SMALPs.* MdtF was extracted from homogenised membranes by the addition of 2.5 % (w/v) SMA 2000 co-polymer powder and incubation for 2 h at room temperature with gentle agitation. Insoluble material was then removed by centrifugation at 100,000 x *g* for 1 h at 4 °C. 1 mL Ni-NTA resin (Generon or Thermo Scientific), equilibrated in Buffer B with 20 mM imidazole, was added directly to the supernatant and incubated overnight at 4 °C with gentle agitation. The sample was then transferred to a gravity flow column (Bio-Rad) and washed with 20 column volumes of Buffer B with 20 mM imidazole and 10 column volumes of Buffer B with 50 mM imidazole. MdtF was eluted with 5 column volumes of Buffer B with 500 mM imidazole. The eluted protein was filtered through a 0.22 μm filter membrane (Fisher Scientific) and loaded onto a Superdex 200 Increase 10/300 GL SEC column (GE Healthcare) equilibrated in Buffer B. A flow rate of 0.4 mL/min was used during SEC purification. Peak fractions eluted from the SEC column containing pure MdtF were pooled, spin concentrated using a 100 K MWCO Vivaspin^®^ 6 spin concentrator (Sartorius), flash-frozen in liquid nitrogen and stored at -80 °C. Samples were separated by SDS-PAGE in a 12 % NuPage^TM^ Precast Gel to assess MdtF purification. Gels were run at 180 V and room temperature for 1 h. Protein concentration was calculated using a NanoPhotometer^TM^ N60 UV/Vis spectrophotometer (Implen) with an extinction coefficient of ε_280_ = 86,305 M^−1^cm^−1^.

### DLS Measurements

10 μL of sample was diluted in 990 μL of deionised water in a 4.5 mL disposable cuvette. Dynamic light scattering (DLS) measurements were performed on a Litesizer^TM^ 500 (Anton Paar) at 25 °C for 20 processed runs. The size distribution in radius is displayed on a logarithmic scale.

### Negative Stain EM Analysis

MdtF-SMALPs were initially filtered through a 0.22 μm Spin-X centrifuge tube filter (Costar) and buffer exchanged into a Tris-based Protein Buffer (50 mM Tris, 150 mM sodium chloride, pH 7.4) using Micro Bio-Spin 6 columns (Bio-Rad). Samples were then spin concentrated using a 100 K MWCO Vivaspin^®^ 500 spin concentrator (Sartorius).

#### Negative Stain EM Grid Preparation and Imaging

Negative stain grids were prepared by applying 3 μL of the respective protein samples onto a previously glow-discharged carbon-coated copper (300 mesh) grid. The grids were then stained with 3 % uranyl acetate for 1 min. The grids were air dried for 2 min and imaged using a JEM-1400 FLASH (JEOL) TEM equipped with a tungsten filament operating at 120 kV. Digital images were acquired at nominal magnification of 60,000x with a bottom-mounted FLASH 2 k x 2 k CMOS camera.

### Cryo-EM Sample Preparation and Data Acquisition

MdtF samples were filtered through a 0.22 μm Spin-X centrifuge tube filter (Costar) and buffer exchanged into a Tris-based Protein Buffer (50 mM Tris, 150 mM sodium chloride, pH 7.4) using Micro Bio-Spin 6 columns (Bio-Rad). For the R6G-bound state, MdtF^V610F^ samples were treated similarly except for the addition of 75 μM R6G followed by a 4 h incubation on ice. To prepare grids for vitrification, 4 μL of apo-MdtF^WT^ at 0.25 mg mL^−1^, apo-MdtF^V610F^ at 0.5 mg mL^−1^, and R6G-MdtF^V610F^ at 0.5 mg mL^−1^ were applied to previously glow-discharged UltrAufoil 1.2/1.3 gold or Quantifoil R1.2/1.3 holey carbon EM grids (Quantifoil Micro Tools). Cryo-EM samples were subsequently blotted using a Mark IV Vitrobot (Thermo Fisher Scientific) for 4 s with a blot force of 7 at 4 °C and 100 % humidity prior to immediate plunge-freezing in liquid nitrogen-cooled ethane. Cryo-EM specimens were screened using a Glacios^TM^ Cryo-TEM (Thermo Fisher Scientific) to examine particle density and ice quality.

For apo-MdtF^WT^, 5,675 movies were collected on a Glacios^TM^ Cryo-TEM with a Falcon III detector, operating at 200 kV accelerating voltage and a nominal magnification of 190,000x, corresponding to a physical pixel size of 0.946 Å pixel^−1^. The apo-MdtF^V610F^ and R6G-MdtF^V610F^ datasets were collected using a similar approach, except 8,000 movies were recorded for both on a Falcon 4 camera with a physical pixel size of 0.947 Å pixel^−1^ and a nominal magnification of x 150,000.

### Image Processing

The image processing steps for each dataset are depicted in *Supplementary* Figs. 3,5*, and 7*. In brief, movie stacks were pre-processed with Relion’s implementation of MotionCor2^47,48^ using a 5×5 grid (25 tiles), for motion correction and dose-weighting according to the implemented pre-calibrated filtering scheme. The micrograph-based CTF was determined with CTFFIND4^49^ on non-dose weighted, drift-corrected micrographs. A subset of micrographs with a CTF fit accurate up to 8 Å or better were selected. Particles were subsequently auto-picked in Relion-4.0^48^ using Laplacian-of-Gaussian with a diameter between 100 and 150 nm. Particles were extracted with a box size of 260×260 pixels, after binning by a factor of two. Low quality particles and false-positive picks in the automatically picked dataset were identified by a reference-free 2D classification, separating particles into 200 classes. Of these 2D classes, representative classes were subsequently used for a reference-based particle picking in Relion-4.0^48^. Subsequent classifications and refinements were performed in Relion-4.0. For 3D classification, a map of AcrB (EMD-24653)^50^ was low-pass filtered to 60 Å and used as an initial reference. 3D classification was performed with no symmetry imposed (C1). From three 3D classes, the particle images comprising the 3D class with better features corresponding to MdtF were subsequently un-binned to the original pixel size. To prevent the loss of any asymmetric structural features of MdtF, all steps were undertaken with no applied symmetry. Following Bayesian polishing, CTF refinement, and beam tilt estimation with Relion-4.0^48^, refinement of this subset of particles with a soft mask and no applied symmetry (C1), produced maps at nominal resolutions of 3.56, 3.28, and 3.20 Å (gold-standard Fourier shell correlation (FSC) = 0.143) for the apo-MdtF^WT^, apo-MdtF^V610F^, and R6G-MdtF^V610F^, respectively.

### Model Building and Structure Refinement

A homology model was obtained for MdtF^WT^ using SwissModel^51^, which was used as a starting point for modelling with Coot^52^. This model was subsequently used as a starting point for generating the atomic models for apo-MdtF^V610F^ and R6G-MdtF^V610F^. Models for lipids within the transmembrane region were modelled using Coot^52^ as previously described^28^. Briefly, PE or dodecane molecules were modelled into the structures with varying chain lengths which corresponded to the electron density they occupied. Atomic models were refined in Phenix^53^ using real-space refinement and validated with Molprobity^54^. ChimeraX^55^ was used for visualisation of Cryo-EM maps and preparation of figures.

### Fluorescence Polarisation Assay

MdtF ligand binding was determined using fluorescence polarisation (FP) assays as performed by Su *et al*^38^. MdtF protein titration experiments were conducted in ligand binding solution (50 mM sodium phosphate, 150 mM sodium chloride, 10 % (w/v) glycerol, 1 μM R6G, pH 7.4). FP measurements were recorded following incubation for 10 min for each corresponding protein concentration to ensure binding had reached equilibrium. All measurements were performed at 25 °C and the excitation wavelengths were set to 525 nm and fluorescence polarisation signals were measured at an emission wavelength of 550 nm. Each data point was an average of 15 FP measurements and ligand binding data were fitted to a hyperbola function: 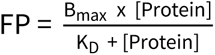 using SigmaPlot (Version 14.5) as conducted by Su *et al*^38^.

### Nile Red Efflux Assay

*E. coli* Δ9-Pore cells^56^ were transformed with either pUC19-MdtEF^WT^, pUC19-MdtEF^V610F^, pUC19-MdtEF^D408A^, or p151A-AcrAB plasmids. An overnight LB culture was added to 50 mL of pre-warmed LB culture containing 100 μg/mL ampicillin at a starting OD_600_ of 0.05. Cells were sub-cultured with 0.1 % L-arabinose and grown at 37 °C and 200 rpm until an OD_600_ of 1.0 was reached. Bacteria were then centrifuged and resuspended in 10 mL of PBS, containing 1 mM MgCl_2_ (PPB). Cells were pelleted again and resuspended in 10 mL of PPB. Samples were separated into 2 mL aliquots with an OD_600_ of ∼1.0 within 10 mL Pyrex® disposable glass conical centrifuge tubes to avoid Nile Red adherence to the walls. 10 μM of Carbonyl cyanide m-chlorophenylhydrazone (CCCP, Cambridge Bioscience) was added and were incubated for 20 min at 37 °C and 200 rpm. Following this incubation, 5 μM of Nile Red was added to cells and were incubated for an additional 1 h 30 min at 37 °C and 200 rpm. Cells were pelleted and washed twice with PPB buffer and further diluted 10-fold. Nile Red fluorescence was measured at 650 nm over 250 s, at 1 s intervals (λ_ex_ = 544 nm, slit width of 5 nm; λ_em_ = 650 nm, slit width of 10 nm) using a Fluoromax^®^-4 with FluorEssence^TM^ (Horiba). 50 mM glucose was added after 50 s to re-energise the proton-powered pumps. The resulting fluorescence curves were normalised to a starting value of 1.0. All curves are an average of at least three independent runs undertaken with *E. coli* Δ9-Pore cells which have been separately transformed with either pUC19-MdtEF^WT^, pUC19-MdtEF^V610F^, pUC19-MdtEF^D408A^, or p151A-AcrAB plasmids.

### MdtEF Expression Test

An overnight culture of *E. coli* Δ9-Pore cells harbouring pUC19-MdtEF, or empty pUC19 plasmids were sub-cultured with 0.1 % L-arabinose until they reached either exponential (OD_600_ = 0.8) or stationary (OD_600_ = 2.0) phase. Here, 1 mL of each culture was pelleted by centrifugation at 16,000 *x g* at 4 °C for 5 min and washed twice with PBS. The pellets were resuspended in 100 mL of ice-cold RIPA lysis buffer (1 mM EDTA, 1 % Triton X, 0.1 % sodium deoxycholate, 0.1 % SDS, 140 mM NaCl, 10 mM Tris-HCl, pH 8.0), supplemented with 100 mM PMSF and a protease inhibitor tablet, and incubated at 4 °C with shaking for 10 min. The cells were subsequently sonicated for 15 min and insoluble contents were removed by centrifugation at 16,000 *x g* at 4 °C for 5 min. The supernatant was aliquoted and mixed with SDS loading buffer prior to loading 20 mL on a 4-12 % NuPAGE^TM^ Bis-Tris Protein Gel. For detection of protein using western blotting, the protein was transferred to a nitrocellulose membrane and an anti-His antibody was used to detect the His-tag containing proteins.

### Minimal Inhibitory Concentration Assay

Susceptibilities of the *E. coli* Δ9-Pore cells harbouring various plasmid-borne MdtF variants against ciprofloxacin, crystal violet, deoxycholate, doxorubicin (Fluorochem Limited), erythromycin, linezolid, novobiocin, PAβN, rhodamine 6G, rifampicin, SDS, and tetracyline (Abcam) were determined by two-fold broth microdilution as previously described^57^. Briefly, overnight cultures were sub-cultured in LB broth (tryptone, 10 g/l; yeast extract, 5 g/l; NaCl, 5 g/l), and cells were grown at 37 °C in a shaker at 220 rpm until OD_600_ reach to 0.2–0.3. For the proper expression of the pore, L-arabinose (final concentration of 0.1 %) was added to each culture, and cells were further grown until OD_600_ reached 1.0. The minimum inhibitory concentration of each bacterial strain against different antibiotics was measured in 96-well plates. Exponentially growing cells were added to each well and incubated for 18 h. European Committee on Antimicrobial Susceptibility Testing (EUCAST) guidelines were followed conforming to ISO 20776-1:2006^58,59^. Substrates were prepared and used according to the manufacturer’s instructions. *E. coli* Δ9-Pore cells harbouring an empty pUC19 plasmid was used as the negative control.

### SDS-PAGE

Proteins were diluted in 5 x Laemmli sample buffer (312.5 mM Tris, 50 % glycerol, 100 mg/mL SDS, 80 mg/mL dithiothreitol (DTT), 0.1 % bromophenol blue, pH 6.8). Samples were run on either a 4-10, 10, or 12 % pre-cast NuPAGE^TM^ Bis-Tris or Novex Tris-Glycine Gel (Thermo Fisher Scientific). Bis-Tris gels were run with a 20 x NuPAGE^TM^ MOPS SDS running buffer (50 mM MOPS, 50 mM Tris Base, 0.1 % SDS, 1 mM EDTA, pH 7.7). Tris-Glycine gels were run with a 10 x Tris-Glycine running buffer (25 mM Tris Base, 192 mM glycine, pH 8.3). The Novex Sharp Pre-Stained Protein Standard (Thermo Fisher Scientific) was used as a protein ladder for molecular weight estimation. Samples were run at room temperature for 1 h at 180 V and protein bands were subsequently visualised using Coomassie® Brilliant Blue G-250 stain (VWR).

### Western blotting

For anti-His Western blotting, SDS-PAGE gels were transferred to a nitrocellulose membrane (0.2 μm pore size) membrane. Membranes were subsequently equilibrated in transfer buffer (25 mM Tris, 192 mM glycine, 0.77 % (w/v) SDS, 10 % (v/v) methanol) and proteins were then transferred from the cell to the membrane using a Cytvia Amersham^TM^ TE 77 PWR semi-dry transfer unit (fisher scientific). Transfer was run at room temperature for 1 h at 45 A per gel. Following this, membranes were subsequently blocked in 5 % (w/v) milk powder in PBS-Tween (PBS-t, 0.1 % (v/v) Tween) for 1 h at room temperature or 4 °C overnight with gentle shaking. Membranes were further incubated with anti-polyhistidine-HRP antibody (1:10,000) dilutions for 1 h at room temperature or 4 °C overnight with gentle shaking. Membranes were subsequently washed with PBS-t four times at 15 min intervals. Membranes were then developed with 1 mL Amersham^TM^ ECL Select^TM^ solution for 1 min and imaged using an A1600 Imager (GE Healthcare).

### Native SMA-PAGE

Native SMA-PAGE was performed as previously described^60^. In brief, samples were run on either a pre-cast Novex^TM^ Value^TM^ Tris-Glycine protein gel, 4-20 % (Thermo Fisher Scientific). Protein samples were diluted in 2 x Novex^TM^ Tris-Glycine Native sample buffer (Thermo Fisher Scientific) or 4 x NativePAGE sample buffer (Thermo Fisher Scientific). The NativeMark^TM^ Unstained Protien Standard (Thermo Fisher Scientific) was used as a protein ladder for molecular weight estimation. 30 μL of protein was loaded per well and samples were run for 90 min at 150 V and 4 °C. Protein bands were subsequently visualised using Coomassie® Brilliant Blue G-250 stain (VWR).

### Lipid Extraction

For lipidomics and GC-MS experiments, lipids were extracted according to a modified version of Bligh and Dyer^61^. In brief, lipid-containing samples (0.5 mL) were added to 1.7 mL chloroform : methanol : 1 M Tris at pH 8 (10:23:1 (vol/vol/vol)) and mixed extensively. To achieve phase separation, 1 mL of a 1:1 mixture of chloroform and 0.1 M Tris at pH 8 was added. The lipid-containing organic phase was then collected and evaporated under a stream of nitrogen to provide a total lipid extract film.

### GC-MS

#### Fatty Acid Derivatisation

Fatty acyl methyl esters (FAMEs) were prepared by derivatisation of the lipid-containing samples for identification of CFAs using GC-MS. In brief, 0.5 mg of dry lipid extract was dissolved in 100 μL toluene, 750 μL methanol, and 150 μL 8 % HCl solution. Following an hour incubation at 100 °C, 0.5 mL hexane and 0.5 mL water was added. The mixture was then vortexed and centrifuged at 6,000 *x g* for 5 min. The FAME-containing organic phase was separated, and the aqueous phase was re-extracted with 250 μL of hexane. The organic phases were combined and used for fatty acid analysis by GC-MS.

*GC-MS*. The FAME mixture was separated on a Shimadzu QP2020 NX GC-MS instrument using an SH-RXI-5MS column (30 m x 0.25 mm x 0.25 μm, Shimadzu). The temperature gradient was as follows: 150 °C (4 min); 4 °C/min to 250 °C (11 min). The carrier gas was helium with a linear flow rate of 25.5 cm/s. The injector temperature was 250 °C and the injection volume was 1 μL with a 10:1 split. Detection was performed using Selected Ion Monitoring (SIM) and a commercial mixture of bacterial acid methyl esters was used for identification of fatty acids based on their retention time (RT).

*GC-MS Analysis*. The raw data were processed using GCMSsolution software (Shimadzu). Signal peaks were identified using ions m/z = 74 collected in the SIM mode and their RT and sample peak areas were calculated. GC-MS values are presented as relative proportions within each sample.

### Lipidomics

Lipids extracts were prepared with an internal standard mix as previously described^30^. All samples were analysed via liquid chromatography-mass spectrometry (LC-MS) using a Vanquish Flex LC (Thermo Fisher, Hemel Hempstead, UK) with a Acquity BEH C18 column (Waters, Wilmslow, UK) attached, coupled to a Q Exactive Plus mass spectrometer (Thermo Fisher, Hemel Hempstead, UK). Mobile phase A was composed of acetonitrile : water (60:40) with 10 mM ammonium acetate. Mobile phase B was composed of isopropanol : acetonitrile (90:10) with 10 mM ammonium acetate. An 18-minute LC gradient was run at 0.3 μL/min, starting at 40 % mobile phase B, increasing to 99 % B at 10 min, and then decreasing back to 40 % B at 13.5 min to the end of run. The source was operated at 3.2 kV, capillary temperature 375 °C, and desolvation temperature and gas flow 400 °C and 60 L/h respectively. All mass spectra were acquired in negative ion mode in the range of 150-2000 m/z. Data processing and analysis was performed using Expressionist software (v15, Genedata, Basel, Switzerland), for chromatographic and spectral peak detection and isotope clustering. Identification of lipids was performed via accurate mass against the LIPIDMAPs database.

### Functional Assays and Immunoblotting Normalisation

The HT and NPN uptake assays were conducted using *E. coli* Δ9-Pore cells complemented with either the pUC19 vector or its derivative carrying the gene encoding MdtEF^WT^. The HT uptake experiment was performed with both exponential-phase cells, collected 4 h after subculture, and stationary-phase cells, collected 20 hours post-subculture. Overnight cultures of *E. coli* Δ9-Pore strains (pUC19 and MdtEF^WT^) were diluted 1:100 in fresh LB and incubated at 37 °C with shaking until reaching mid-log phase (OD_600_ ∼0.3–0.4). For conditions requiring pore induction, 0.1 % L-arabinose was added, and the cultures were allowed to grow for either 4 h (exponential phase) or 20 h (stationary phase). In the absence of L-arabinose, cultures were grown for the respective time periods without induction. After incubation, cultures were adjusted to OD_600_ ∼1.0, centrifuged, and resuspended in HMG buffer (50 mM HEPES-KOH, 0.4 % (w/v) glucose, 1 mM MgSO₄, pH 7.0) at the appropriate pH. The cells were then readjusted to OD_600_ ∼1.0 before proceeding with the uptake assay. For studies of the pH effect, induced exponential *E. coli* Δ9-Pore cells were collected by centrifugation, washed, and resuspended in the buffer adjusted to different pH values to final OD_600_ ∼1.0.

For the uptake assay, a 96-well black clear-bottom plate was prepared with increasing concentrations of HT as previously described^62^. Briefly, bacterial suspensions at OD_600_ ∼1.0 were added to each well (100 μl per well), and fluorescence measurements were taken immediately upon cell addition using a spectrophotometer. The fluorescence settings were optimised for HT (excitation: 355 nm, emission: 450 nm). Steady-state concentrations were calculated using MATLAB, and raw fluorescence data were analysed and plotted in Microsoft Excel and GraphPad Prism for visualisation and comparison.

For protein expression analysis, aliquots of bacterial cell suspensions prepared for the uptake experiments above were harvested by centrifugation. Cells were lysed via sonication, and lysates were subjected to differential centrifugation, first at 4,000 rpm to remove debris, followed by ultracentrifugation at 40,000 rpm to obtain membrane fractions. The total protein concentration was measured using the Bradford assay. Equal amounts of protein were resolved via SDS-PAGE and transferred onto a PVDF membrane for western blot analysis. The membrane was blocked with 1 % BSA and probed with an anti-His primary antibody, followed by an anti-mouse alkaline phosphatase-conjugated secondary antibody.

### Differential Scanning Fluorimetry (DSF)

NanoDSF experiments were performed as follows. 10 μL of 2 mg/mL MdtF-SMALPs ± 75 μM R6G was loaded into standard capillaries and onto the capillary tray of a Prometheus Panta instrument (Nanotemper Technologies). The experiment was run at a temperature gradient from 10-100 °C at 1 °C/min, an excitation power of 7 % and DLS power of 100 %. The data for each channel was subsequently analysed using the PR.Panta Analysis Software (Nanotemper Technologies) to analyse thermal stability and particle size.

### Molecular Docking and Simulations

#### Molecular Docking

We investigated the binding of R6G, PAβN, linezolid, ciprofloxacin, and nitrosyl oxide at physiological pH. These studies were conducted for both the MdtF^WT^ and MdtF^V610F^ transporter (utilising two ensemble structures for each variant), focusing on the T protomer using the cryo-EM structures of apo and holo states of MdtF^WT^ and MdtF^V610F^ (WT bound structure was generated using homology modelling by Prime using mutant V610F bound) for the distal pocket, whereas only apo structure were used for the channels docking. The protein structures of MdtF (obtained in this work) were prepared using the *Protein Preparation Wizard* in Maestro (Schrödinger Release 2024-1) with the default settings. This preparation involved adjusting for missing hydrogen atoms, incomplete side chains and loops, assigning appropriate protonation states, and flipping residues to reflect the physiological condition^63^. Following this, we performed a restrained minimisation that permitted free movement of hydrogen atoms (threshold of < 0.30 Å) while allowing sufficient relaxation of heavy atoms to alleviate strained bonds and angles. The missing loops (residues 496-515) were modelled and refined using Prime using the reference UniProt sequence P37637 utilising OPLS4 force field and implicit solvation model VSGB^64–66^. In particular, at the proton relay site in transmembrane domain, protonation states were assigned following Eicher *et al*^34^, i.e, E346 and D924 were protonated only in L and T protomers, while D407, D408, and D566 were protonated only in the O protomer^34,67,68^. For the ligands modelling, possible ionisation states were generated using Epik (*Ligprep*) at pH 7.4^69^. Ligand conformation sampling for each pH condition was executed using Macromodel^70^ (Mixed Mode). The optimised conformers of the five ligands were docked against the grid generated centring around the native ligand (Rhodamine 6G) for the distal pocket, and centred around specific residues for CH1 (T559, S834, G836, A838, K840, E864, L866, S868, Q870, P872), CH2 (E564, T643, Q647, I651, V660, T674, S676, G713, R715, N717, E720, L826, E828) and CH3 (A33, Q37, P100, G296,N298)^71^ residues using standard precision mode (SP)^72,73^. Top 20 poses were further processed for the second docking run using extra precision (XP) mode^73^. Best poses corresponding to the Glide score were ranked and chosen to analyse the interaction details. Prior to docking, the redocking (as a control) was performed to confirm the performance and accuracy of the protocol for the R6G in the DP. For our entire ligand and receptor modelling, we adhered to OPLS4 force field^74^. The images were plotted using pymol^75^.

#### Molecular Dynamics (MD)

MD simulations of apo proteins were performed for three systems: MdtF (MdtF^WT^ and MdtF^V610F^), and the well-established AcrB structure in the asymmetrical LTO state (PDB ID: 4DX5)^76^. The structure of AcrB was prepared following the same protocol described for MdtF in the docking section. Three chains of consistent lengths used and corresponding residues for AcrB/MdtF were protonated in the LTO states. In MdtF (AcrB) residues E346 (E346) and D922 (D924) were protonated only in the L and T protomers, while residues D407 (D407), D408 (D408), and D566 (D568) were protonated only in the O protomer. All other residues were protonated at physiological pH^67,68^. Amber24 was used for the system setup and simulation^77,78^. The structures were converted to GAFF2 atom types using pdb4amber (Ambertools23), during which hydrogens were removed and re-added, and protonation states for the proton relay site were assigned^77^. Membranes were inserted using PACKMOL-Memgen^79^ whereby the protein is embedded in a mixed bilayer patch composed of 1-palmitoyl-2-oleoyl-sn-glycero3-phosphoethanolamine (POPE) and 1-palmitoyl-2-oleoyl-sn-glycero-3-phosphoglycerol (POPG) in a 2:1 ratio, for a total of ∼730 lipid molecules symmetrically distributed across the two bilayer leaflets. The AMBER force field ff19SB^80^ was used to represent the protein, the lipid21^81^ parameters were used for the phospholipids, the OPC model was employed for water^82^, and ions parameters were taken from Joung *et al*^83^. A potassium ion concentration of 0.15 M was added, and the lipid packing was optimised with a 15 Å distance from the protein and a 17.5 Å water layer. Randomisation ensured proper lipid placement, and minimisation was performed post-generation. The system was built iteratively over 100 cycles with a convergence tolerance of 2.4 Å for optimal structural stability. *parmed* was used to repartition the mass of hydrogen atoms.

Each system was first subjected to a multistep structural relaxation via a combination of steepest descent and conjugate gradient method using *pmemd.cuda* program implemented in AMBER24 as described previously^84–89^. The system was then heated in two stages: first, from 0 to 100 K over 1 ns under constant-volume conditions applying harmonic restraints (k = 1 kcal·mol^−1^·Å^−2^) to the heavy atoms of both the protein and the lipids, subsequently, temperature was increased from 100 to 310 K under constant pressure (1 atm) for 1 ns, with harmonic restraints (k = 2 kcal·mol^−1^·Å^−2^) applied to the Cα atoms of the protein and *z* coordinates of the phosphorous atoms (P31) in lipids. This approach allowed for membrane rearrangement during the heating process. The system was equilibrated in a series of six equilibration steps at 1 fs time steps for 250 ps each (total 1.5 ns) with restraint on the protein coordinates under isotropic pressure scaling using the Berendsen barostat, whereas a Langevin thermostat (with a collision frequency of 1 ps^−1^) was used to maintain a constant temperature. Finally, production simulations were performed at 310 K with a 4 fs time step employing Langevin thermostat and anisotropic pressure scaling (under an isothermal-isobaric ensemble), with outputs recorded every 25,000 steps (100 ps). The Particle mesh Ewald algorithm was used to evaluate long-range electrostatic forces with a non-bonded cut-off of 9 Å. For each system (MdtF^WT^, MdtF^V610F^, and AcrB), three independent replicates were performed, each for 1 μs, resulting in a total simulation time of 3 μs per system. Trajectories post-processing was performed with *cpptraj*. The contacts at the PC1-PC2 domain by Cα atoms were calculated using “*nativecontact*” command (skipping native contacts) with a cut-off of 10 Å^90^. Extreme structures of maximum or minimum contacts (correspond to closing and opening of cleft) each associated with their respective minimum and maximum distances were chosen. The hbonds were calculated using “hbond” command with a distance cutoff 3.0 Å; angle cutoff 135°. The cut-off distance for salt bridge interaction was 3.2 Å between O and N atoms, calculated using *cpptraj*.

Hydration of the transmembrane (TM) region within 10 Å of the proton relay site (PRS) was analysed using the grid tool in AMBER24, calculating the average water density (cut-off > 0.25) along the cumulative trajectory. Densities were normalised to bulk water (1 g/mL), and significant hydration sites (corresponding gridpoints) were identified. Dynamic exchange was assessed by tracking water molecules that entered within 5 Å of the PRS in at least 1 % of trajectory frames per monomer. Their single water trajectories were analysed within a 30 Å of PRS.

### Differential Scanning Calorimetry

Conformational stability was assessed by Differential Scanning Calorimetry (DSC). MdtF^WT^ and MdtF^V610F^ were prepared at 7.5 μM, ± 75 μM R6G (10 x molar excess) in a 96-well plate, centrifuged at 4,000 *x g* for 5 min to remove air and loaded onto an automated MicroCal VP DSC (Malvern Panalytical, Great Malvern, UK). A temperature gradient from 10-100 °C, at a scan rate of 1 °C/min was performed, using a pre-scan thermostat of 15 min, a filtering period of 5 s and in passive feedback mode. Scans were automatically buffer subtracted, concentration corrected, and a manual baseline set and subtracted in the MicroCal PEAQ-DSC software v1.64.

### Circular dichroism (CD) spectroscopy

All CD spectra were measured in 20 mM sodium phosphate, 150 mM sodium fluoride, pH 7.4 at a final concentration of 1 μM. Far-UV CD spectra were recorded on an Chirsacan CD spectrophotometer (Applied Photophysics, Leatherhead, UK), from 250-190 nm, with a step size of 0.5 nm, a bandwidth of 1 nm and a cuvette pathlength of 10 mm. A continuous temperature ramp from 20-90 °C, at 2 °C intervals at a rate of 0.23 °C / min was applied. The transition mid-points (Tm) of the spectra were analysed using a multi-wavelength fitting algorithm in Global 3 Analysis Software (Applied Photophysics, Leatherhead, UK). Scans were averaged, buffer subtracted and converted to mean residue ellipticity [θ] according to;

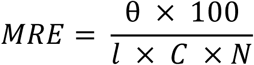

Where θ is the mean residue ellipticity (deg cm^2^ dmol^−1^), mdeg is the raw CD, MRW is the mean residue weight (g/mol), *l* is the cuvette pathlength (cm), *C* is protein concentration (g/L), *N* is the number of residues. Spectra were plotted in GraphPad Prism.

## Acknowledgements

M.A. and A.V.V. gratefully acknowledge the “One Health Basic and Translational Research Actions addressing Unmet Needs on Emerging Infectious Diseases (INF-ACT)” foundation by the Italian Ministry of University and Research, PNRR, mission 4, component 2, investment 1.3, project number PE00000007 (University of Cagliari) and the NIH/NIAID grant no. R01AI136799. The studies at King’s College London and the University of Southampton were supported by a UKRI Future Leaders Fellowship (MR/S015426/1 and MR/X009580/1) to E.R. and a BBSRC iCASE studentship with UCB Pharma (BB/T008709/1) to R.L. The studies at University of Southampton were also supported by a BBSRC industry studentship with Vitacress to L.S. (BB/T008768/1). The studies at the University of Oklahoma, USA were supported by the NIH/NIAID grant RO1AI132836 to H.I.Z.

## Data availability

Density maps and structure coordinates have been deposited in the Electron Microscopy Data Bank (EMDB) and the Protein Data Bank (PDB) with the following accession codes: apo-MdtF^WT^ (EMD-53281 and PDB 9QPR), apo-MdtF^V610F^ (EMD-53282 and PDB 9QPS), R6G-MdtF^V610F^ (EMD-53283 and PDB 9QPT). Source data are provided with this paper. MD simulation trajectories and the docking poses at the DBP, CH1, CH2 and CH3 are available at zenodo: 10.5281/zenodo.15038634.

## Additional information Contributions

R.L., H.I.Z., A.V.V., Z.A., and E.R. designed the project; R.L. and C.A. performed cryo-EM experiments; R.L. and J.S. performed cryo-EM analysis; R.L. and E.R. performed structural investigation and analyses; R.L. and O.D. performed biophysical experiments and analysis; R.L. cloned, purified, and characterized all protein constructs; M.R.U. and H.I.Z. performed bacterial accumulation assays and analysis; R.L. performed bacterial efflux assays and analysis; R.L. and L.S. performed bacterial susceptibility assays and analysis; R.L. performed lipid extraction and GC-MS; S.L. performed lipidomic experiments and analysis; M.A. and A.V.V. carried out molecular docking and MD simulations and post-MD analyses; C.W.K, N.P., C.P, D.M., H.I.Z., A.V.V., Z.A., and E.R. supervised and/or financially supported the project; R.L., M.A., H.I.Z., A.V.V., and E.R. wrote the manuscript with input from the other authors.

