## Supplementary Information for "Molecular basis for multidrug efflux by an anaerobic RND transporter"

### Supplementary Discussion

#### Molecular docking at channels 1, 2 and 3 in MdtF compared to AcrB

Three substrate channels already identified in AcrB can also be seen within MdtF (**Supplementary Fig. 26**). Channel 1 (CH1) provides substrate access from the periplasm and preferably exports various antibiotic including tetracycline[1]. The export of high molecular mass (HMM) compounds is preferentially facilitated by channel 2 (CH2) which enables periplasmic drug access through a cleft opening between PC1 and PC2 subdomains[2]. Substrates which enter via these two channels pass consecutively through the proximal binding pocket (PBP) and then DBP, whereby its selection is mediated by the switch loop[3]. Several key amino acids surrounding the CH2 in MdtF (F662, R715, T674, and M650) differ from those in AcrB. These residues in MdtF point inwards towards the entry channel and, therefore, may affect how the binding site interacts with substrates (**Supplementary Fig. 27**). Interestingly, the export of PACs, such as ethidium bromide and R6G, are mainly mediated by channel 3 (CH3) which provides access via the central cavity. Here, the compounds gain direct access to the DBP and bypass the PBP and switch loop[1–3]. CH3 also exhibits dissimilarities between AcrB and MdtF; in particular, residues surrounding the entrance to the gate of CH3, likely to be essential in substrate recognition and binding[3], are inherently different between AcrB (T37 and A100) and MdtF (Q37 and P100) (**Supplementary Fig. 26**). The presence of a structurally disruptive proline and a polar molecule attracting glutamine may facilitate a better comprehension of the suggested preference of MdtF for cationic, flat, aromatic substances (especially dyes) in comparison to AcrB[4]. In agreement with this hypothesis, molecular docking of a range of different drug-like molecules (PAβN, linezolid, ciprofloxacin and R6G), as well as its natural nitrosyl indole substrate, on MdtF revealed an overall larger affinity to CH3 than to CH2, particularly for PAβN and ciprofloxacin (**Supplementary Table 3**).

To determine the difference in functionality owing to the CH3 entry gate, we tested the resistance phenotype of MdtF harbouring these mutations Q37T and P100A through implementing MIC assays (**Supplementary Fig. 26** and **Supplementary Table 4**). Although this demonstrated that the mutations did not significantly affect the MdtF resistance phenotype, the ability of MdtF to export PACs was corroborated due to its conferred resistance to crystal violet, doxorubicin, and R6G. Interestingly, MdtF was also able to confer resistance to deoxycholate, a naturally occurring secondary bile acid found in human gut. Here, the acidic and anaerobic environment of the human gut correlates with the expression pattern of MdtF which may enable bacteria to tolerate the hostile conditions it may encounter in the human enteric pathway. In summary, this information alongside the conformational alterations adopted by MdtF (specifically, a smaller PC1-PC2 crevice) may suggest a distinct antiport mechanism through which it is primarily a CH3 exporter. This is because CH3 bypasses the PBP and would not require the same structural movements exhibited by AcrB. Due to the preference of CH3 for PACs, this could allude to the functional importance of MdtF within acidic conditions because of the specificity of cationic substrates for CH3. As the access protomer is in a 'primed' state for ligand binding, the structural alterations in transmembrane domain movements may force CH3 into an open conformation to facilitate export of PACs. Here, the expulsion of aromatic acids, cationic antibiotics, bile acids, etc. can recover the depleted PMF in extreme acidic conditions by being more readily transported into the cytoplasm, bringing protons with them, and enabling its function.

Interestingly, both single-point mutants, $MdtF^{Q37T}$ and $MdtF^{P100A}$, displayed a reduced resistance to the macrolide antibiotic erythromycin which was reversed in the presence of the double mutant $MdtF^{Q37T/P100A}$. This resistance phenotype effect arising from a combinatorial mutation indicates an epistatic interaction between these two residues. This genetic cooperativity could indicate a significant impact and functional interdependency of these residues on the evolutionary trajectory of RND transporters in erythromycin-specific transport.

**Supplementary Table 1 | Cryo-EM data collection, refinement and validation statistics**

| | Apo-MdtF$^{WT}$ | Apo-MdtF$^{V610F}$ | MdtF$^{V610F}$-R6G |
|---|---|---|---|
| | EMD-53281 | EMD-53282 | EMD-53283 |
| | PDB 9QPR | PDB 9QPS | PDB 9QPT |
| **Data collection and processing** | | | |
| Magnification | x 190,000 | x 150,000 | x 150,000 |
| Voltage (keV) | 200 | 200 | 200 |
| Electron exposure ($e^-/Å^2$) | 48.82 | 41.25 | 41.25 |
| Defocus range (µm) | -0.8 to -2.2 (-0.2 steps) | -0.8 to -2.2 (-0.2 steps) | -0.8 to -2.2 (-0.2 steps) |
| Pixel size (Å) | 0.946 | 0.947 | 0.947 |
| Symmetry imposed | C1 | C1 | C1 |
| Initial particle images (no.) | 2,501,990 | 5,069,376 | 4,498,131 |
| Final particle images (no.) | 593,735 | 1,446,003 | 1,967,418 |
| Map resolution (Å) | 3.56 | 3.28 | 3.2 |
| FSC threshold | 0.143 | 0.143 | 0.143 |
| Map resolution range (Å) | … | … | … |
| **Refinement** | | | |
| Initial model used | Homology model | 9QPR (this work) | 9QPS (this work) |
| Model resolution (Å) | 3.14 | 2.87 | 2.84 |
| FSC threshold | 0.143 | 0.143 | 0.143 |
| Model resolution range (Å) | 3.1-3.6 | 2.7-3.3 | 2.8-3.3 |
| Map sharpening *B* factor ($Å^2$) | -194 | -188 | -177 |
| Model composition | | | |
| Non-hydrogen atoms | 23,906 | 24,004 | 24,062 |
| Protein residues | 3,051 | 3,056 | 3,054 |
| Ligands | 21 (PTY)<br>7 (D12) | 22 (PTY)<br>11 (D12) | 23 (PTY)<br>12 (D12)<br>1 (RHQ) |
| *B* factors ($Å^2$) | | | |
| Protein | 9.46/87.37/38.22 | 0.00/101.12/37.30 | 0.00/110.35/44.01 |
| Ligand | 42.53/12.12/82.87 | 55.90/124.22/81.90 | 41.77/133.24/89.22 |
| R.m.s. deviations | | | |
| Bond lengths (Å) | 0.005 | 0.003 | 0.003 |
| Bond angles (degrees) | 0.668 | 0.556 | 0.540 |
| Validation | | | |
| MolProbity score | 2.00 | 1.76 | 1.70 |
| Clashscore | 11.25 | 8.47 | 8.62 |
| Poor rotamers (%) | 0.32 | 0.28 | 0.28 |
| Ramachandran plot | | | |
| Favoured (%) | 93.34 | 95.66 | 96.38 |
| Allowed (%) | 6.66 | 4.34 | 3.62 |
| Disallowed (%) | 0 | 0 | 0 |

| Name | Sequence (5' - 3') | Use |
|---|---|---|
| sGFP_Del_Fwd | CTC GAG CAC CAC CAC CAC | Deletion of TEV-sGFP region using Q5® mutagenesis kit from pET15b-MdtF-sGFP-6xHis plasmid; forward primer |
| sGFP_Del_Rev | GCT CTG GAA GTA CAG GTT TTC AC | Deletion of TEV-sGFP region using Q5® mutagenesis kit from pET15b-MdtF-sGFP-6xHis; reverse primer |
| MdtF_V610F_Fwd | GGT GTT TAC CTT TGG CGG CTT TG | Q5® mutagenesis of V610F in *mdtF* gene, reverse primer |
| MdtF_V610F_Rev | GAC TGG ACA TTA TCT TTC TCT TTA GTC | Q5® mutagenesis of V610F in *mdtF* gene, forward primer |
| pUC19_MdtEF_Fwd | GCC AAG CTT GCA TGC CTG CAG AAC TGT TGG CAG AAC GG | Cloning *mdtEF* genes with its natural promoter from K-12 *Escherichia coli* chromosomal DNA into a pUC19 plasmid (linearised with *Pst*I and *Bam*HI restriction enzymes); forward primer |
| pUC19_MdtEF_His_Rev | ATT CGA GCT CGG TAC CCG GGT CAG TGG TGG TGG TGG TGG TGC TCG AGG CTC TGG AAG TAC AGG TTT TCA CCG CTA GCC GCT TTT TTA AAG CGG GC | As delineated above, however, a 6xHistidine tag sequence was included in the reverse primer to provide a 6xHis tag at the C-terminus of MdtF; reverse primer |
| pUC19_MdtEF_Tag_Del_Fwd | TGA CCC GGG TAC CGA GCT | Deletion of TEV-6xHis region using Q5® mutagenesis kit from puC19-MdtEF-TEV-6xHis plasmid; forward primer |
| pUC19_MdtEF_Tag_Del_Rev | CGC TTT TTT AAA GCG GGC AAA GAG | Deletion of TEV-6xHis region using Q5® mutagenesis kit from puC19-MdtEF-TEV-6xHis plasmid; reverse primer |
| MdtF_Q37T_Fwd | GCA GTA TCC GAC GAT TGC GCC AC | Q5® mutagenesis of Q37T in *mdtF* gene, forward primer |
| MdtF_Q37T_Rev | GCA ACC GGT AAG TTC ATG | Q5® mutagenesis of Q37T in *mdtF* gene, reverse primer |
| MdtF_P100A_Fwd | TGG GAC ATC TGC GGA TAT CGC AC | Q5® mutagenesis of P100A in *mdtF* gene, forward primer |

| | | |
|---|---|---|
| MdtF_P100A_Rev | GTC TCG AAG GTC AGA GTG | Q5® mutagenesis of P100A in *mdtF* gene, reverse primer |
| MdtF_Y327A_Fwd | GGT TTA TCC TGC TGA CAC CAC GCC G | Q5® mutagenesis of Y327A in *mdtF* gene, forward primer |
| MdtF_Y327A_Rev | GTC TTC AGA CTT GCC GGG | Q5® mutagenesis of Y327A in *mdtF* gene, reverse primer |
| MdtF_D408A_Fwd | GTT GGT GGA TGC GGC CAT CGT TGT GG | Q5® mutagenesis of D408A in *mdtF* gene, forward primer |
| MdtF_D408A_Rev | AGG CCG ATG GCG AGC | Q5® mutagenesis of D408A in *mdtF* gene, reverse primer |
| Q37T_Seq_Rev | TGC TCG ACT TAT CGA CGC TAA TC | Sequencing of Q37T mutation; reverse primer |
| P100A_Seq_Fwd | AGG TGG TCT GGC GAT CAT GAA C | Sequencing of P100A mutation; forward primer |
| Y327A_Seq_Fwd | ATC GCC ATC AAA CTG GCT GC | Sequencing of Y327A mutation; forward primer |
| V610F_Seq_Fwd | ATC GCC ATC AAA CTG GCT GC | Sequencing of V610F mutation; forward primer |
| MdtF_Mid_Seq_Fwd | GAA CTG AAC CGC TTA TCA GC | Sequencing of MdtF middle region; forward primer |
| MdtF_Mid_Seq_Rev | AAC CAA CCT GGA AGT AGA CG | Sequencing of MdtF middle region; reverse primer |
| T7 _Seq_Fwd | TAA TAC GAC TCA CTA TAG GG | Sequencing downstream from T7 promoter for pET15b plasmids; forward primer |
| T7_term_Seq_Rev | CTA GTT ATT GCT CAG CGG T | Sequencing upstream from T7 terminator for pET15b plasmids; reverse primer |
| M13_Seq_Fwd | GTT TTC CCA GTC ACG AC | Sequencing N-terminus of inserts for pUC19 plasmids; forward primer |
| M40_Seq_Rev | CGG ATA ACA ATT TCA CAC AG | Sequencing N-terminus of inserts for pUC19 plasmids; reverse primer |

**Supplementary Table 2 | Primer Table**

| | R6G | | Linezolid | | PAβN | | Ciprofloxacin | | Nitrosyl Indole | |
|---|---|---|---|---|---|---|---|---|---|---|
| | $MdtF^{WT}$ | $MdtF^{V610F}$ | $MdtF^{WT}$ | $MdtF^{V610F}$ | $MdtF^{WT}$ | $MdtF^{V610F}$ | $MdtF^{WT}$ | $MdtF^{V610F}$ | $MdtF^{WT}$ | $MdtF^{V610F}$ |
| **$DBP_{binding}$** | -8.3 | -8.7 | -7.5 | -7.6 | -10.2 | -9.9 | -6.0 | -7.1 | -6.0 | -6.0 |
| **$CH2_{binding}$** | -3.6 | -4.2 | -5.1 | -6.1 | -4.2 | -4.8 | -4.6 | -5.4 | -4.3 | -5.2 |
| **$CH3_{binding}$** | -4.6 | -5.4 | -7.2 | -6.2 | -5.9 | -6.2 | -9.5 | -7.8 | -7.4 | -8.0 |

**Supplementary Table 3 |** Predicted binding affinities (kcal/mol) of various substrates to both $MdtF^{WT}$ and the $MdtF^{V610F}$ variant from the molecular docking calculations (see *Materials and Methods* for details). Binding affinities are reported for R6G, linezolid, PAβN, ciprofloxacin, and nitrosyl indole at three distinct binding sites: the distal binding pocket (DBP, based on the cryo-EM structure from this study), and the CH2 and CH3 channel entrances (selected based on well-characterised corresponding AcrB residues). CH2: Channel 2, CH3: Channel 3, DBP: Distal binding pocket, PAβN (phenylalanine-arginine β-naphthylamide), R6G: Rhodamine 6G.

| | Minimum Inhibitory Concentration (MIC, μg mL$^{-1}$) | | | | | | | | | | | |
|---|---|---|---|---|---|---|---|---|---|---|---|---|
| | Low Molecular Mass Drugs (LMMDs) | | | | | | High Molecular Mass Drugs (HMMDs) | | | Planar Aromatic Cations (PACs) | | |
| | Ciprofloxacin | Deoxycholate | Linezolid | PAβN | SDS | Tetracycline | Erythromycin | Novobiocin | Rifampicin | Rhodamine 6G | Crystal Violet | Doxorubicin |
| **MdtF$^{WT}$** | 0.0078 | 1250 | 2 | 125 | 205 | 0.0975 | 100 | 0.78 | 6.24 | 800 | 0.04 | >200 |
| **MdtF$^{Q37T}$** | 0.0019 | 1250 | 2 | 125 | 205 | 0.04875 | 50 | 3.12 | 1.56 | 800 | 0.04 | >200 |
| **MdtF$^{P100A}$** | 0.0019 | 1250 | 2 | 125 | 205 | 0.0975 | 25 | 3.12 | 1.56 | 800 | 0.04 | >200 |
| **MdtF$^{Q37T/P100A}$** | 0.0019 | 1250 | 2 | 125 | 205 | 0.0975 | 100 | 3.12 | 1.56 | 800 | 0.04 | >200 |
| **MdtF$^{Y327A}$** | 0.0039 | 1250 | 2 | 125 | 125 | 0.04875 | 25 | 1.56 | 1.56 | 800 | 0.04 | >200 |
| **Empty pUC19** | 0.0019 | <78 | 1 | 31.25 | 25.625 | 0.04875 | 0.78 | 0.39 | 6.24 | 3.125 | <0.02 | 1.56 |

Supplementary Table 4 | MIC Calculations of CH3 Mutants

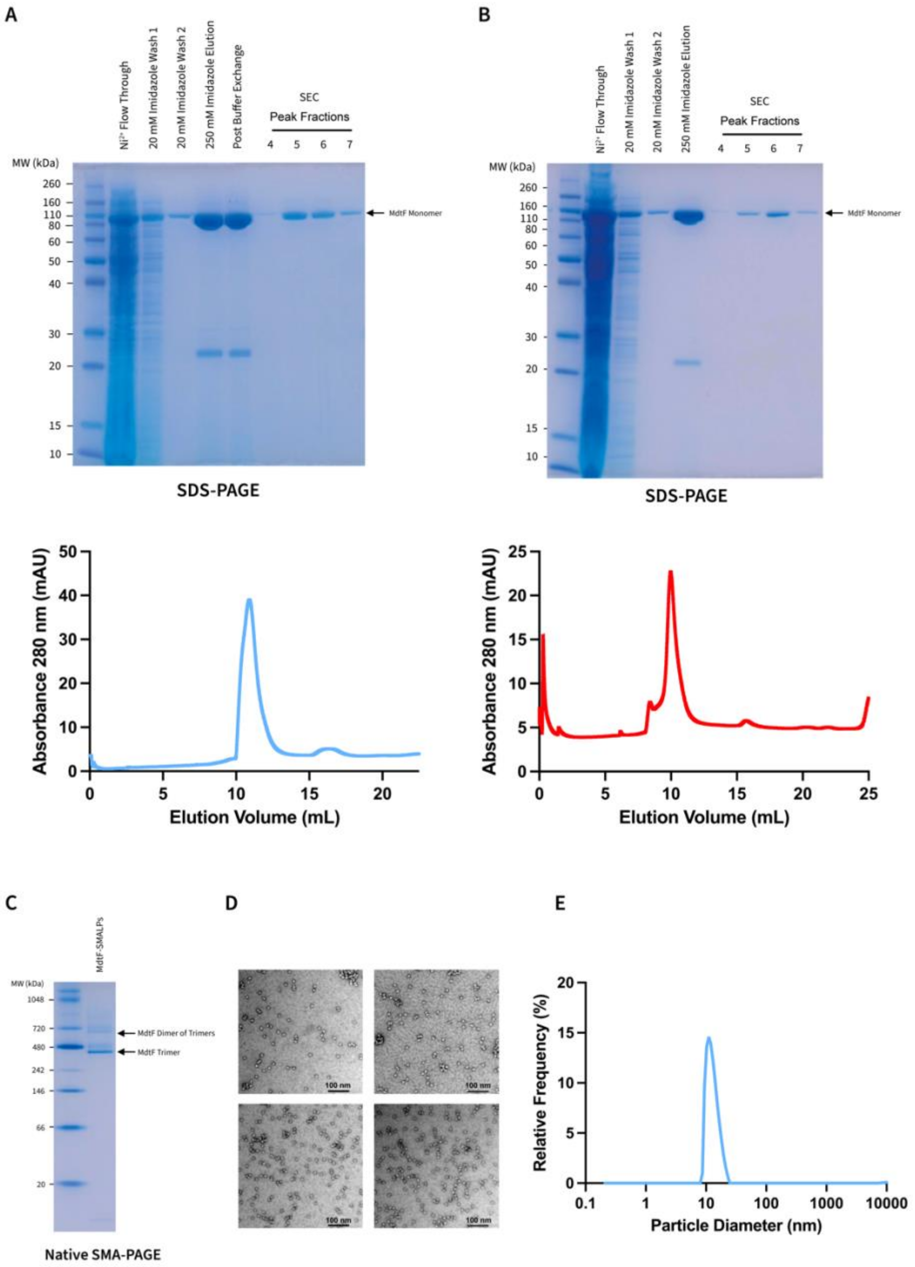


**Supplementary Fig. 1 | Expression, Purification, and Characterisation of MdtF in SMALPs. a and b,** Representative SDS-PAGE gels and corresponding SEC traces of $MdtF^{WT}$ **(a)** and $MdtF^{V610F}$ **(b)**. The MdtF monomer at ~111 kDa is indicated on the SDS-PAGE gel images. **c,** Native SMA-PAGE gel demonstrating the purification of the MdtF homo-trimer within SMALPs, indicated on the gel image. A dimer of the MdtF homo-trimer is also observed due to electrostatic interactions between the SMA nanodisc. **d,** Negative stain EM analysis of MdtF-SMALPs, demonstrating particle monodispersity and homogeneity. Here, the particle size is observed to be ~13 nm, corresponding to the size of AcrB. **e,** Representative DLS trace of MdtF-SMALPs, showing particle homogeneity at approximately 13 nm in accordance with the size observed by negative stain EM analysis.

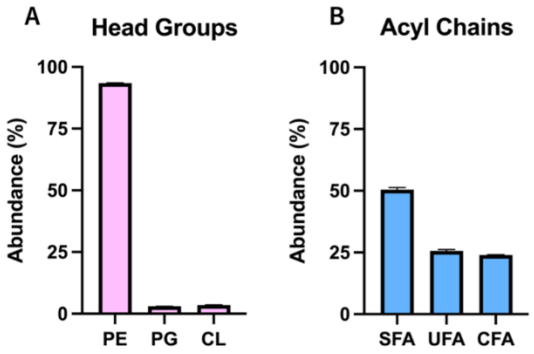


**Supplementary Fig. 2 | Lipidomic Analysis of MdtF-SMALPs Lipid Belt. a,** Lipidomic analysis of lipid head group composition within the purified MdtF-SMALP nanodics. **b,** GC-MS analysis of fatty acyl chain composition within the purified MdtF-SMALP nanodisc. Bars represent mean values from three independent measurements and error bars are indicative of the standard deviation (n = 3). CFA: Cyclopropanated fatty acid, CL: cardiolipin, GC-MS: Gas-chromatography mass spectrometry, PE: Phosphatidylethanolamine, PG: Phosphatidylglycerol, SFA: Saturated fatty acid, UFA: Unsaturated fatty acid.

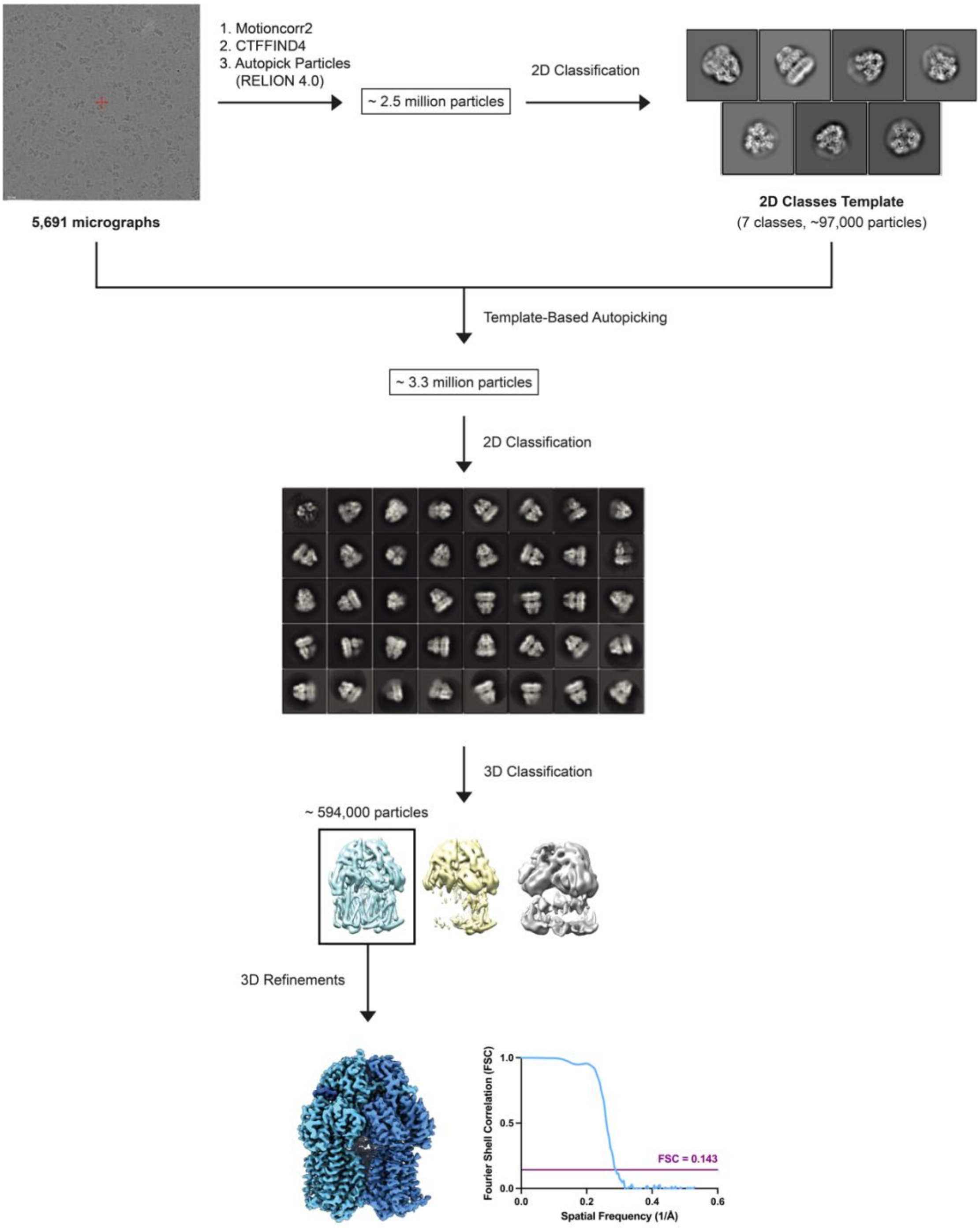


**Supplementary Fig. 3 | MdtF$^{WT}$ Cryo-EM Data Processing.** An outline of the cryo-EM data processing workflows. Representative micrographs with the physical pixel size (50 nm) indicated. The representative 2D classification images and 3D classification models are also displayed. FSC curves of the final 3D reconstruction calculated in Relion-4.0 is also demonstrated. FSC: Fourier shell correlation.

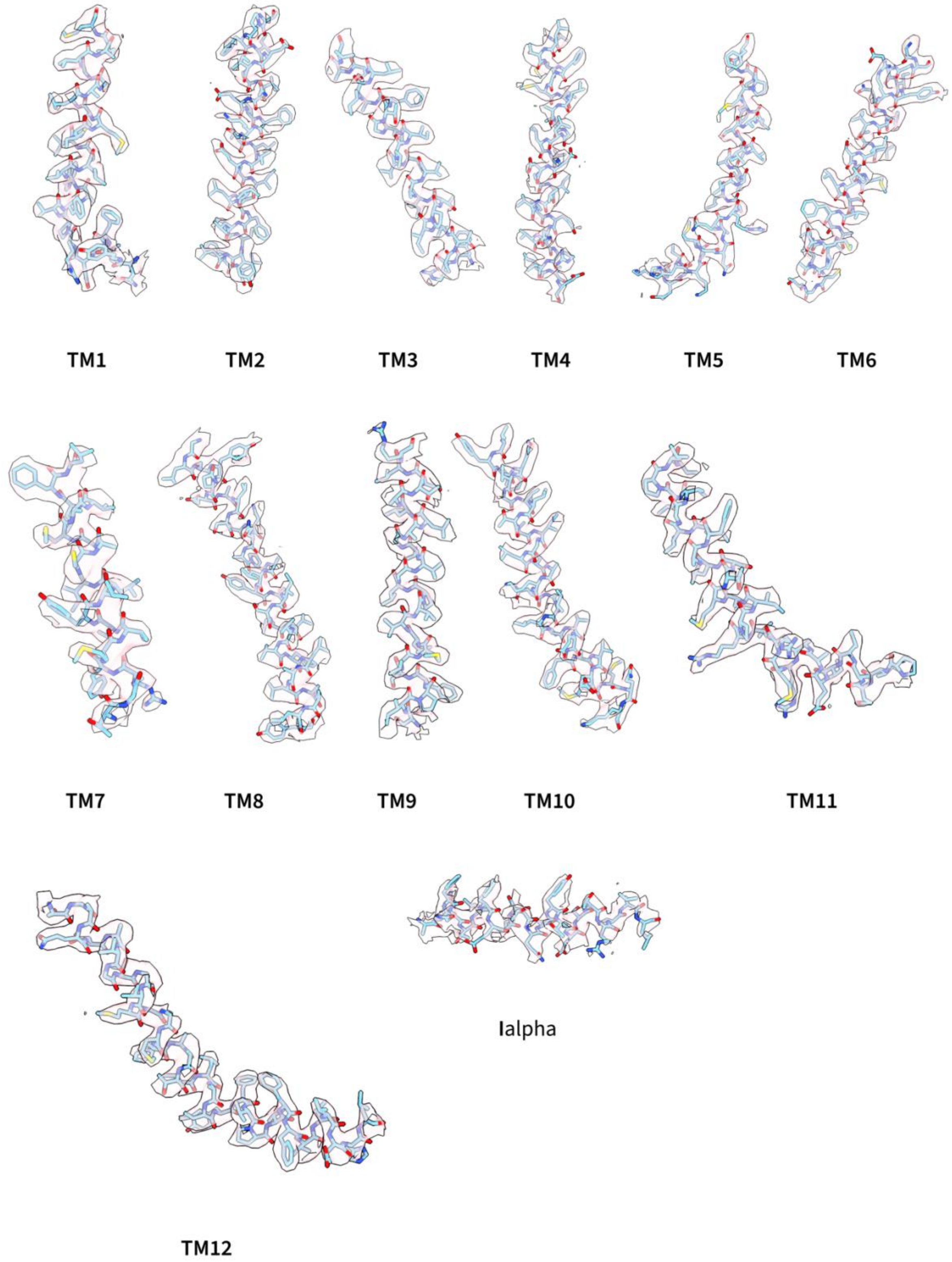


**Supplementary Fig. 4 | Quality of MdtF$^{WT}$ Cryo-EM Map.** Representative cryo-EM densities for the TM helices of MdtF$^{WT}$. The MdtF$^{WT}$ structure is depicted in atom representation (light blue) and the density maps are represented in surface form (light pink). TM: Transmembrane helix.

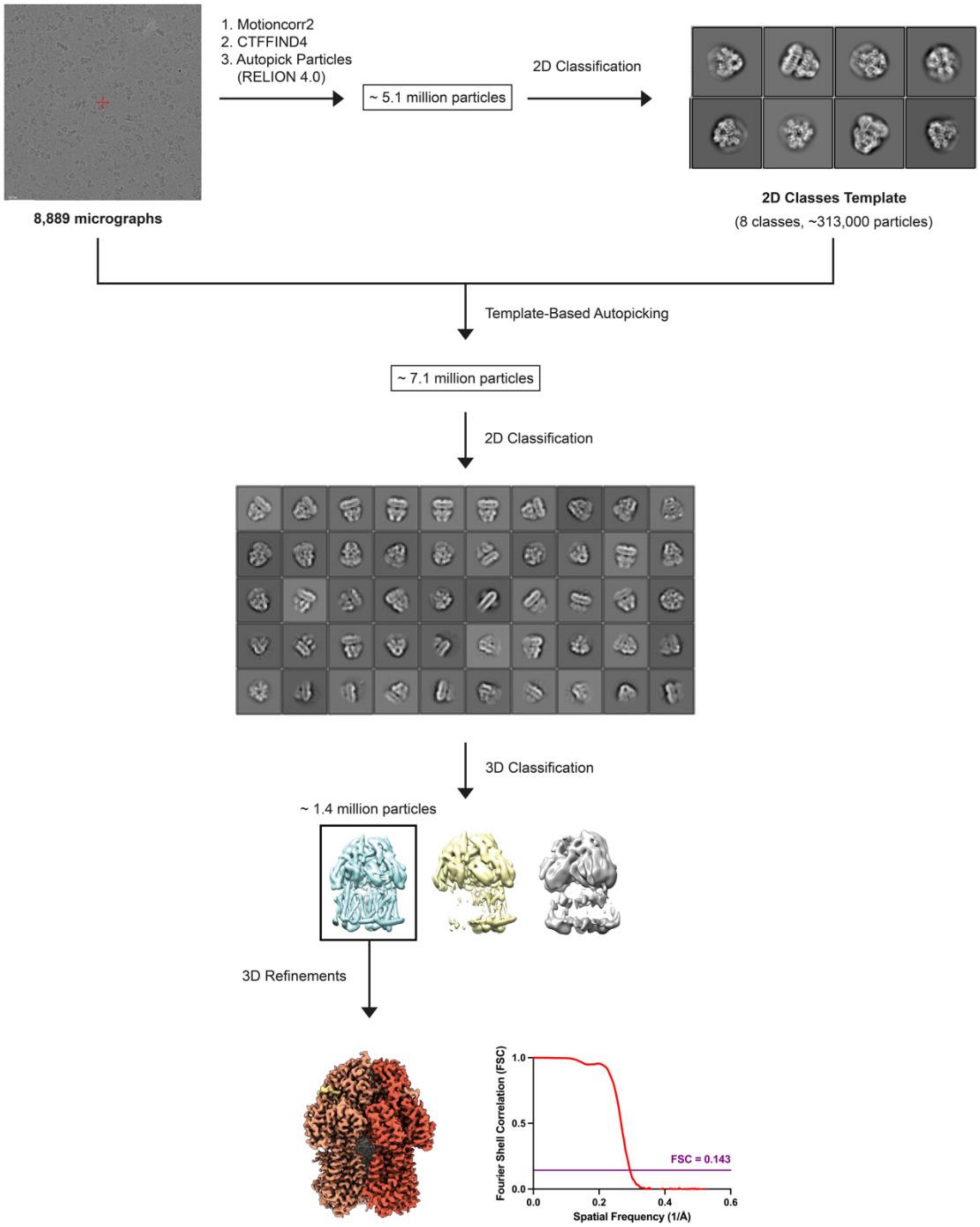


**Supplementary Fig. 5 | MdtF$^{V610F}$ Cryo-EM Data Processing.** An outline of the cryo-EM data processing workflows. Representative micrographs with the physical pixel size (50 nm) indicated. The representative 2D classification images and 3D classification models are also displayed. FSC curves of the final 3D reconstruction calculated in Relion-4.0 is also demonstrated. FSC: Fourier shell correlation.

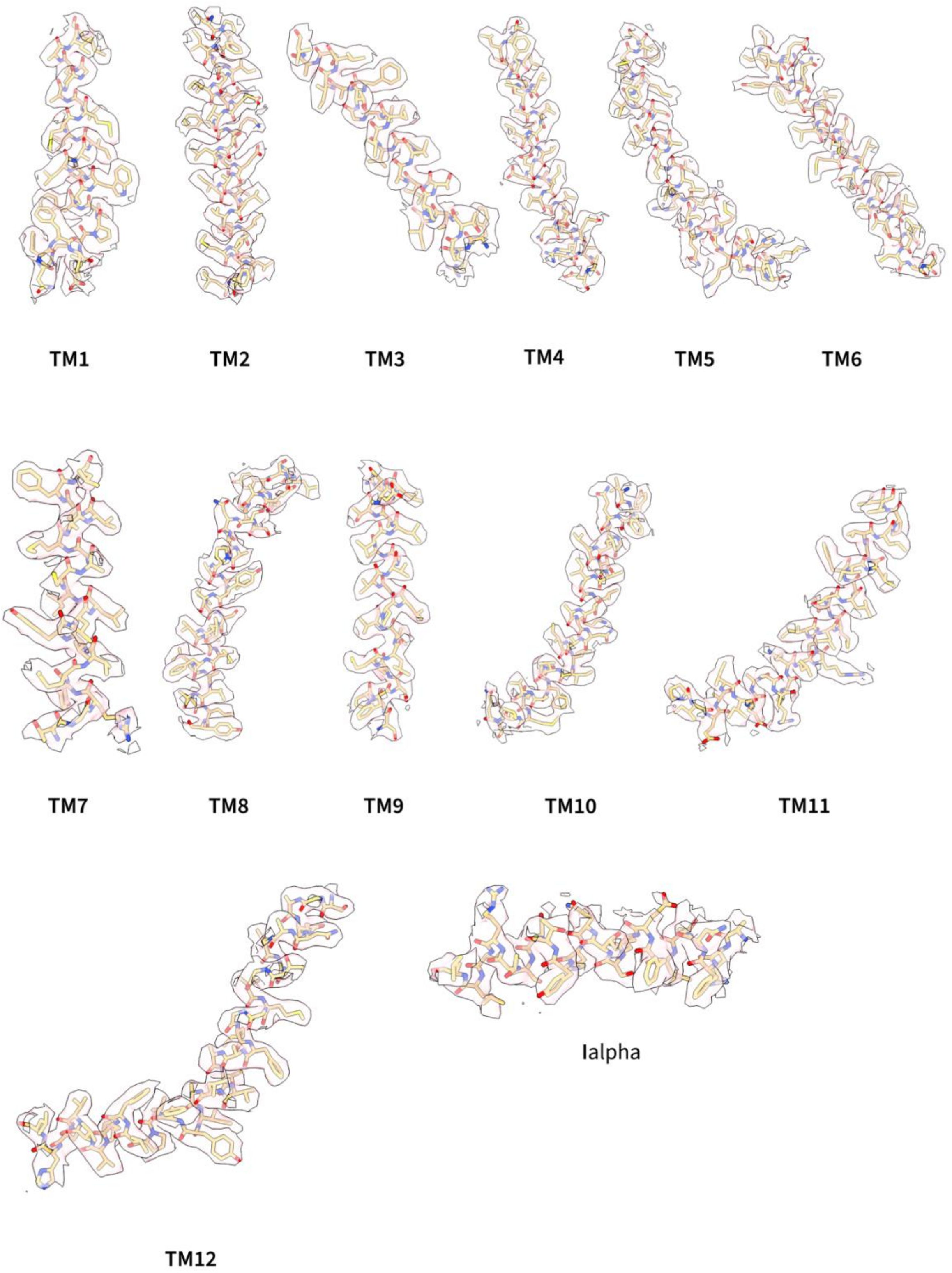


**Supplementary Fig. 6 | Quality of MdtF$^{V620F}$ Cryo-EM Map.** Representative cryo-EM densities for the TM helices of MdtF$^{V610F}$. The MdtF$^{V610F}$ structure is depicted in atom representation (light orange) and the density maps are represented in surface form (light pink). TM: Transmembrane helix.

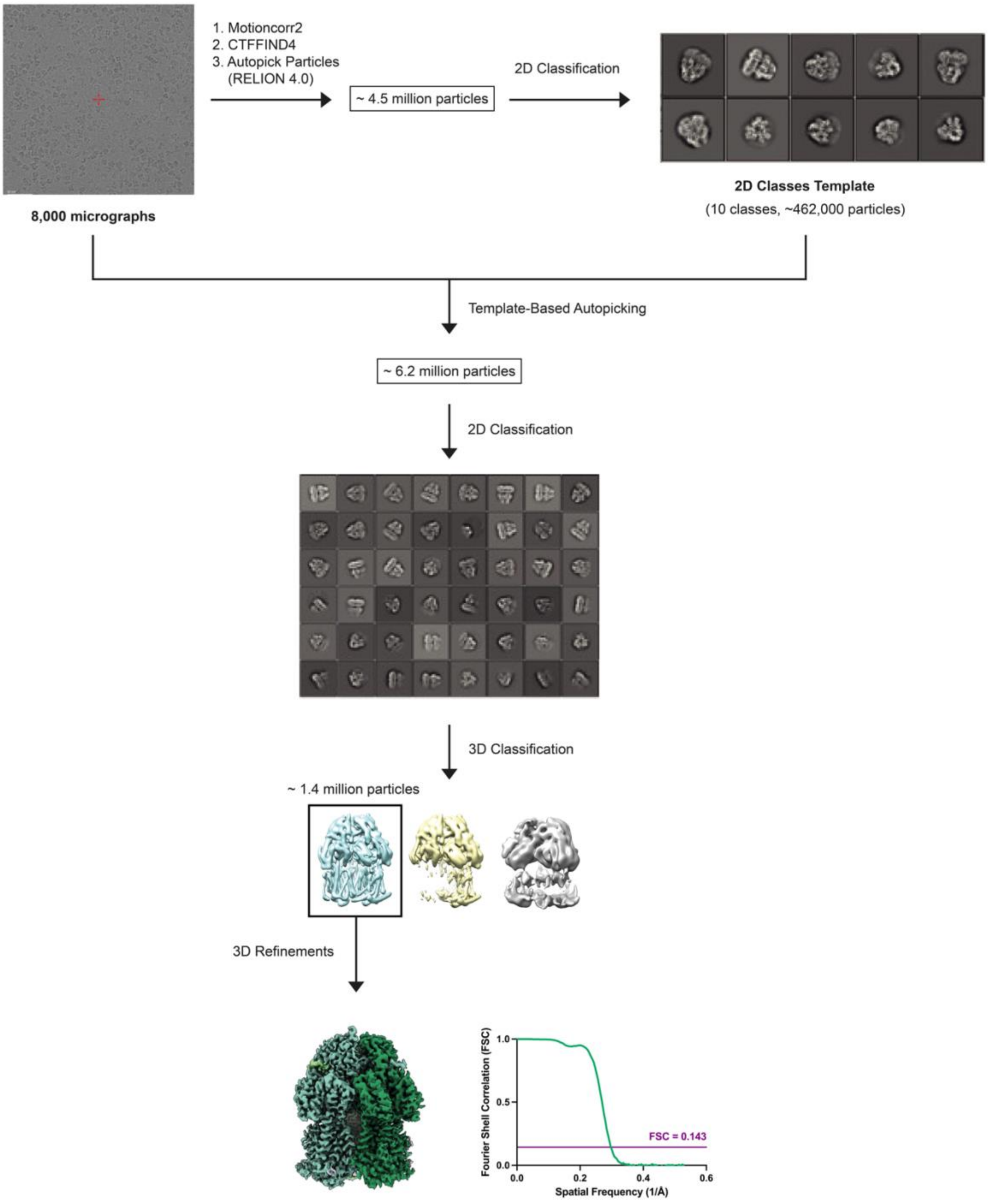


**Supplementary Fig. 7 | MdtF$^{V610F}$-R6G Cryo-EM Data Processing.** An outline of the cryo-EM data processing workflows. Representative micrographs with the physical pixel size (50 nm) indicated. The representative 2D classification images and 3D classification models are also displayed. FSC curves of the final 3D reconstruction calculated in Relion-4.0 is also demonstrated. FSC: Fourier shell correlation.

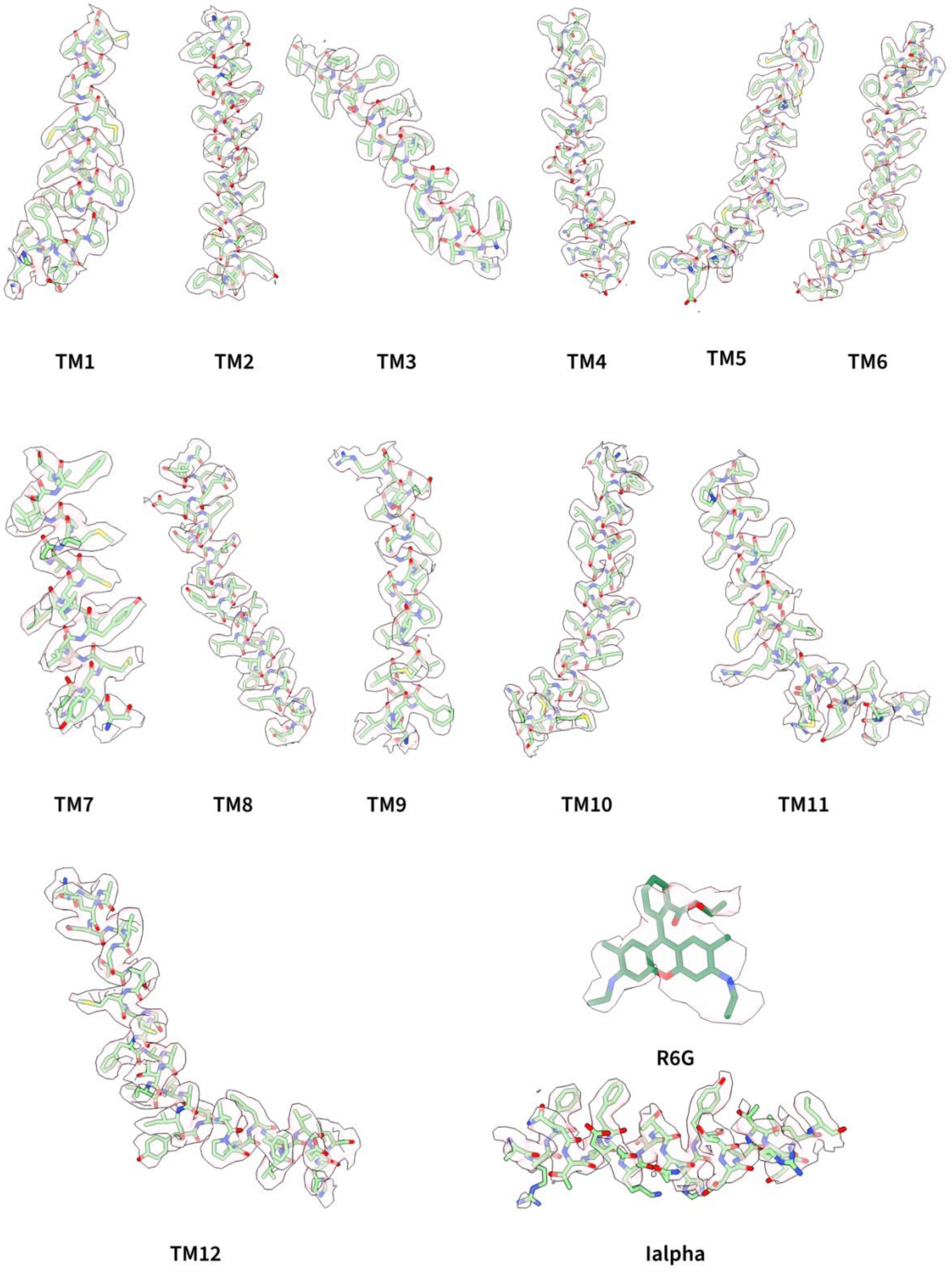


**Supplementary Fig. 8 | Quality of MdtF$^{V610F}$-R6G Cryo-EM Map.** Representative cryo-EM densities for the TM helices and R6G ligand of MdtF$^{V610F}$-R6G. The MdtF$^{V610F}$-R6G structure and R6G ligand are depicted in atom representation (light and dark green, respectively) and the density maps are represented in surface form (light pink). TM: Transmembrane helix.

```
MdtF            MANYFIDRPVFAWVLAIIMMLAGGLAIMNLPVAQYPQIAPPTITVSATYPGADAQTVEDS 60
AcrB            MPNFFIDRPIFAWVIAIIIMLAGGLAILKLPVAQYPTIAPPAVTISASYPGADAKTVQDT 60
                * *:*****:****:***:********::******* ****::*:**:******:**:*:

MdtF            VTQVIEQNMNGLDGLMYMSSTSDAAGNASITLTFETGTSPDIAQVQVQNKLQLAMPSLPE 120
AcrB            VTQVIEQNMNGIDNLMYMSSNSDSTGTVQITLTFESGTDADIAQVQVQNKLQLAMPLLPQ 120
                ***********:*.******.**::*...******:**. **************** **:

MdtF            AVQQQGISVDKSSSNILMVAAFISDNGSLNQYDIADYVASNIKDPLSRTAGVGSVQLFGS 180
AcrB            EVQQQGVSVEKSSSSFLMVVGVINTDGTMTQEDISDYVAANMKDAISRTSGVGDVQLFGS 180
                 *****:**:****.:***...*. :*::.* **:****:*:** :***:***.******

MdtF            EYAMRIWLDPQKLNKYNLVPSDVISQIKVQNNQISGGQLGGMPQAADQQLNASIIVQTRL 240
AcrB            QYAMRIWMNPNELNKFQLTPVDVITAIKAQNAQVAAGQLGGTPPVKGQQLNASIIAQTRL 240
                :******::*::***::*.* ***: **.** *::.***** * . .********.****

MdtF            QTPEEFGKILLKVQQDGSQVLLRDVARVELGAEDYSTVARYNGKPAAGIAIKLAAGANAL 300
AcrB            TSTEEFGKILLKVNQDGSRVLLRDVAKIELGGENYDIIAEFNGQPASGLGIKLATGANAL 300
                 : **********:****:*******::***.*:*. :*.:**:**:*:.****:*****

MdtF            DTSRAVKEELNRLSAYFPASLKTVYPYDTTPFIEISIQEVFKTLVEAIILVFLVMYLFLQ 360
AcrB            DTAAAIRAELAKMEPFFPSGLKIVYPYDTTPFVKISIHEVVKTLVEAIILVFLVMYLFLQ 360
                **: *:: ** ::. :**:.** *********::***:**.*******************

MdtF            NFRATIIPTIAVPVVILGTFAILSAVGFTINTLTMFGMVLAIGLLVDDAIVVVENVERVI 420
AcrB            NFRATLIPTIAVPVVLLGTFAVLAAFGFSINTLTMFGMVLAIGLLVDDAIVVVENVERVM 420
                *****:*********:*****:*:*.**:*******************************:

MdtF            AEDKLPPKEATHKSMGQIQRALVGIAVVLSAVFMPMAFMSGATGEIYRQFSITLISSMLL 480
AcrB            AEEGLPPKEATRKSMGQIQGALVGIAMVLSAVFVPMAFFGGSTGAIYRQFSITIVSAMAL 480
                **: *******:******* ******:******:****:.*:** ********::*:* *

MdtF            SVFVAMSLTPALCATILKAAPEGGH--KPNALFARFNTLFEKSTQHYTDSTRSLLRCTGR 538
AcrB            SVLVALILTPALCATMLKPIAKGDHGEGKKGFFGWFNRMFEKSTHHYTDSVGGILRSTGR 540
                **:**: ********:**   :*.*    :.:*. ** :*****:*****. .:**.***

MdtF            YMVVYLLICAGMAVLFLRTPTSFLPEEDQGVFMTTAQLPSGATMVNTTKVLQQVTDYYLT 598
AcrB            YLVLYLIIVVGMAYLFVRLPSSFLPDEDQGVFMTMVQLPAGATQERTQKVLNEVTHYYLT 600
                *:*:**:* .*** **:* *:****:******** .***:***  .* ***::**.****

MdtF            KEKDNVQSVFTVGGFGFSGQGQNNGLAFISLKPWSERVGEENSVTAIIQRAMIALSSINK 658
AcrB            KEKNNVESVFAVNGFGFAGRGQNTGIAFVSLKDWADRPGEENKVEAITMRATRAFSQIKD 660
                ***:**:***:*.****:*:***.*:**:**:*** *::* ****.* **  **  *:*.*:.

MdtF            AVVFPFNLPAVAELGTASGFDMELLDNGNLGHEKLTQARNELLSLAAQSPNQVTGVRPNG 718
AcrB            AMVFAFNLPAIVELGTATGFDFELIDQAGLGHEKLTQARNQLLAEAAKHPDMLTSVRPNG 720
                *:** *****:.*****:***:**:*:..***********:**: **: *: :*.*****

MdtF            LEDTPMFKVNVNAAKAEAMGVALSDINQTISTAFGSSYVNDFLNQGRVKKVYVQAGTPFR 778
AcrB            LEDTPQFKIDIDQEKAQALGVSINDINTTLGAAWGGSYVNDFIDRGRVKKVYVMSEAKYR 780
                ***** **::::  **:*:**::.*** *:.:*:*.******::********* : : :*

MdtF            MLPDNINQWYVRNASGTMAPLSAYSSTEWTYGSPRLERYNGIPSMEILGEAAAGKSTGDA 838
AcrB            MLPDDIGDWYVRAADGQMVPFSAFSSSRWEYGSPRLERYNGLPSMEILGQAAPGKSTGEA 840
                ****:*.:**** *.* *.*:**:**:.* ***********:*******:** *****:*

MdtF            MKFMADLVAKLPAGVGYSWTGLSYQEALSSNQAPALYAISLVVVFLALAALYESWSIPFS 898
AcrB            MELMEQLASKLPTGVGYDWTGMSYQERLSGNQAPSLYAISLIVVFLCLAALYESWSIPFS 900
                *::* :*.:***:****.***:**** **.****:******:****.*************

MdtF            VMLVVPLGVVGALLATDLRGLSNDVYFQVGLLTTIGLSAKNAILIVEFAVEMMQKEGKTP 958
AcrB            VMLVVPLGVIGALLAATFRGLTNDVYFQVGLLTTIGLSAKNAILIVEFAKDLMDKEGKGL 960
                *********:*****: :***:****************************  ::*:****

MdtF            IEAIIEAARMRLRPILMTSLAFILGVLPLVISHGAGSGAQNAVGTGVMGGMFAATVLAIY 1018
AcrB            IEATLDAVRMRLRPILMTSLAFILGVMPLVISTGAGSGAQNAVGTGVMGGMVTATVLAIF 1020
                ***  ::*.******************:***** *****************.:******:

MdtF            FVPVFFVVVEHLFARFKKA----------          1037
AcrB            FVPVFFVVVRRRFSRKNEDIEHSHTVDHH          1049
                *********.: *:* ::
```

**Supplementary Fig. 9 | Multiple Sequence Alignment of MdtF and AcrB.** The sequence alignment was performed using Clustal Omega[5]. The residue numbering is shown for both MdtF and AcrB. Hydrophobic, negatively charged, and positively charged residues are coloured red, blue, and magenta, respectively. Polar, aromatic, cysteine, glycine, and proline residues are indicated in green. Dashes indicate a gap in the sequence to highlight residue discrepancies between the two proteins. Sequences were obtained from UniProt, Accession codes: P31224 (AcrB) and P37637 (MdtF).

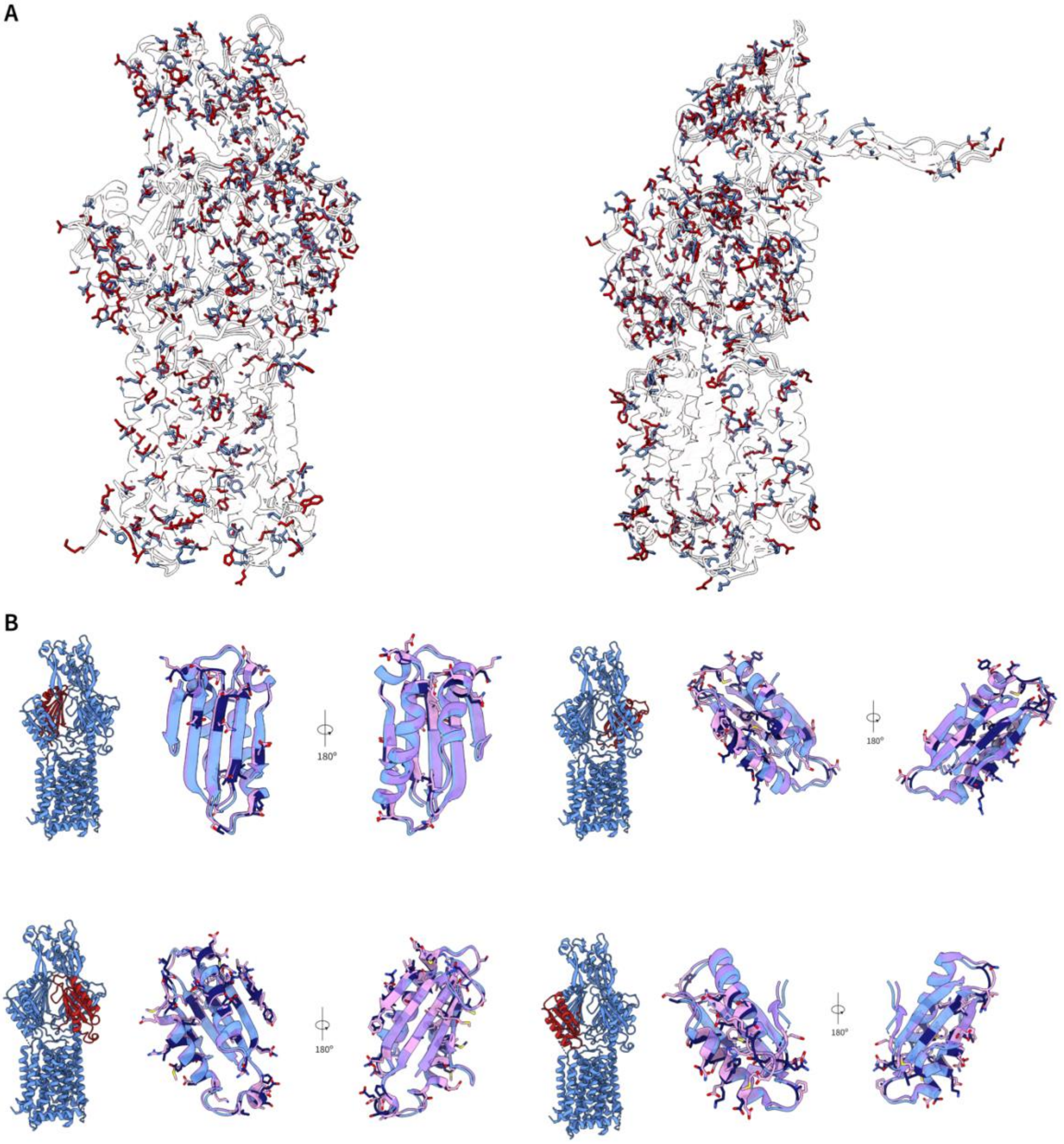


**Supplementary Fig. 10** | **Overlay of different residues observed between MdtF$^{WT}$ and AcrB$^{WT}$. a,** All residues which are not conserved between MdtF$^{WT}$ (blue) and AcrB$^{WT}$ (red) are represented in atomic representation on their respective overlayed structures. This demonstrated the high distribution of different residues across the entire structure. **b,** The different residues observed between each domain of MdtF$^{WT}$ (blue) and AcrB$^{WT}$ (pink). The different residues within MdtF$^{WT}$ which deviate from AcrB$^{WT}$ are located in the interfaces between the domains rather than within the channels and pockets that these domains create.

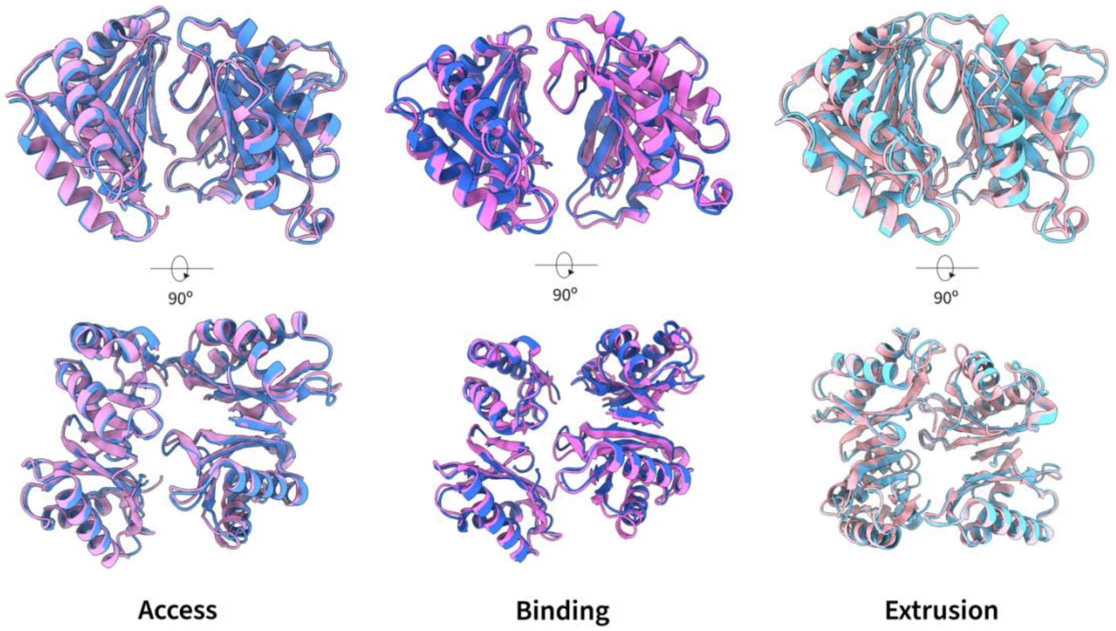


**Supplementary Fig. 11 | Porter domain conformation comparison between MdtF$^{WT}$ and AcrB.** MdtF$^{WT}$ (blue) and AcrB (PDB: 2HRT[6], pink) alignment of porter domain in each of the access, binding, and extrusion states occupied during a drug binding event. A similar conformational change is observed between states in both structures.

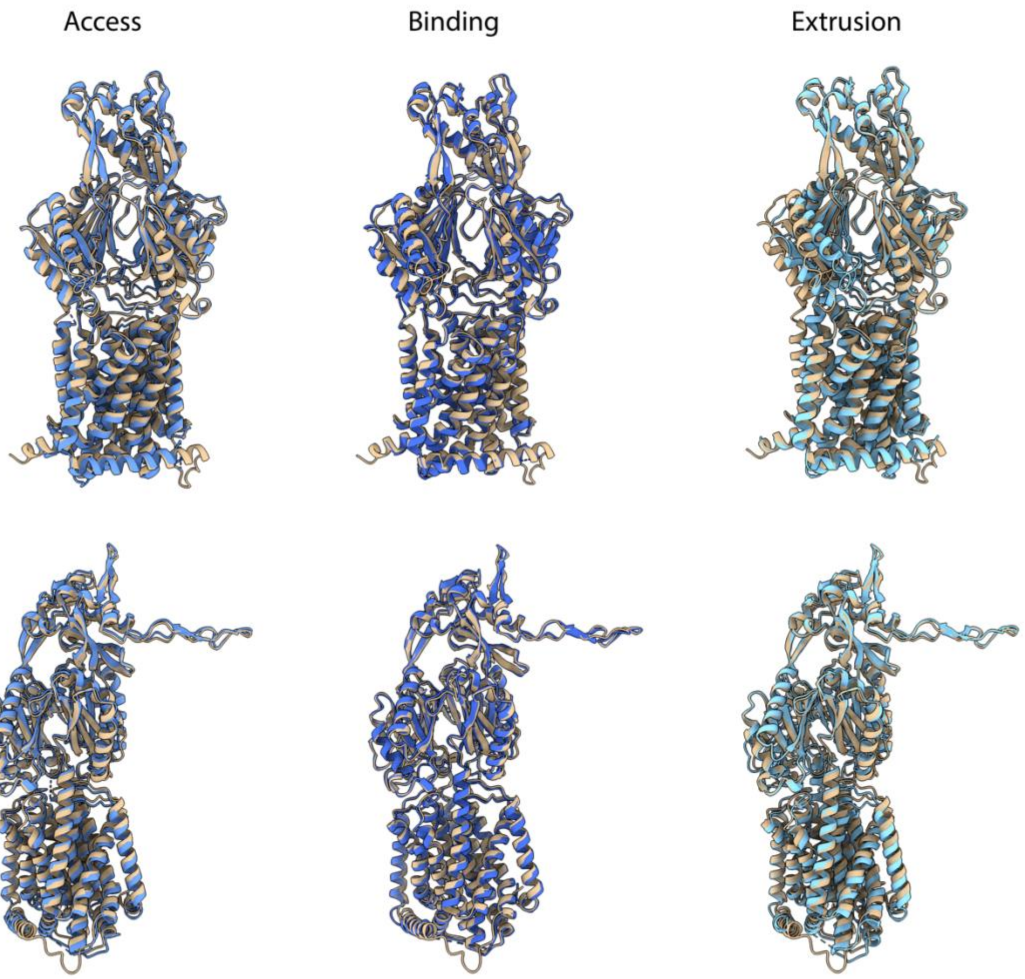


**Supplementary Fig. 12** | **Assessment of AlphaFold Prediction of MdtF Structures.** An analysis of the predicted structure for MdtF by AlphaFold 3[7] was compared to our obtained experimental structure of $MdtF^{WT}$. The model was not successful in predicting the asymmetric structures adopted by $MdtF^{WT}$ observed experimentally which are, ultimately, biologically important.

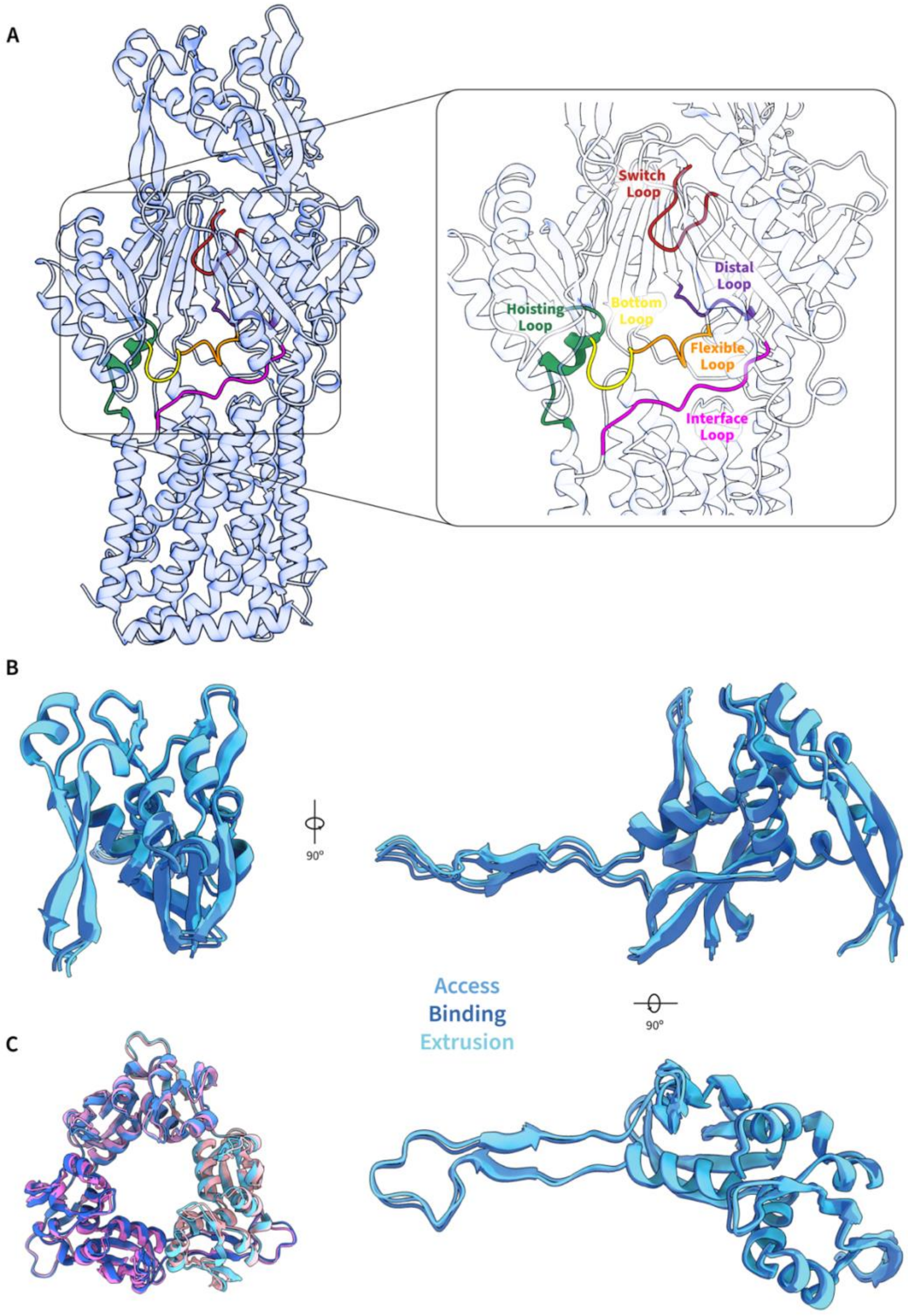


**Supplementary Fig. 13 | Analysis of loops and funnel domains of MdtF$^{WT}$. a,** Conserved RND-related loops within a monomer of MdtF$^{WT}$. **b,** The funnel domain structure within each of the monomeric states of MdtF$^{WT}$ were aligned and demonstrate an overall structural conservation between each protomer. **c,** In comparison to AcrB (pink), MdtF$^{WT}$ (blue) adopts a similar funnel domain structure.

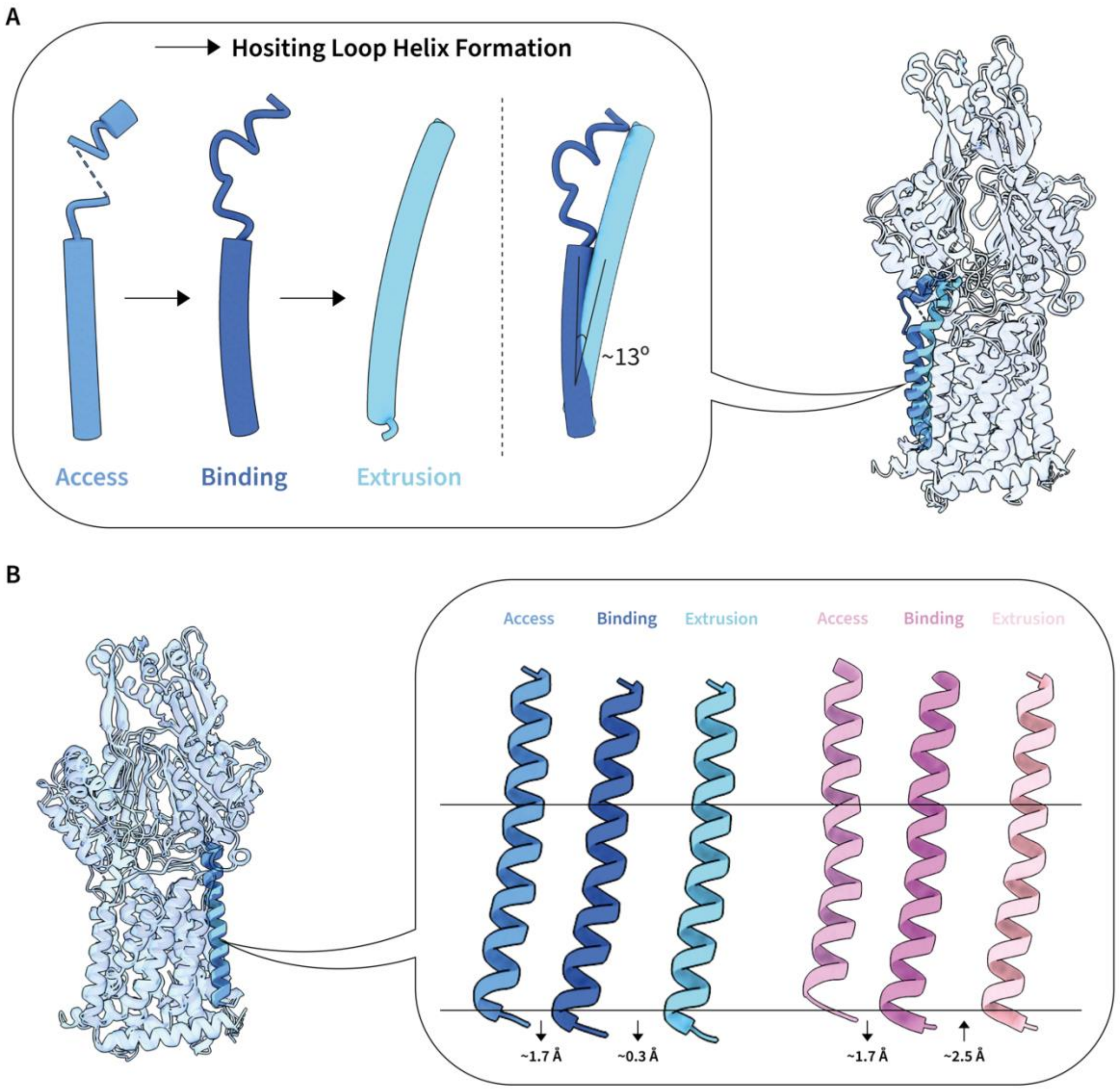


**Supplementary Fig. 14 | Structural transitions observed in TM2 and TM8. a,** The formation of a helical structure in the hoisting loop of TM2 from access to binding to extrusion is conserved within our $MdtF^{WT}$ structure. **b,** The vertical transitions between access, binding, and extrusion states in AcrB and $MdtF^{WT}$ as measured from its distance from the inner membrane. $MdtF^{WT}$ undergoes a reduced vertical transition between the binding and extrusion states.

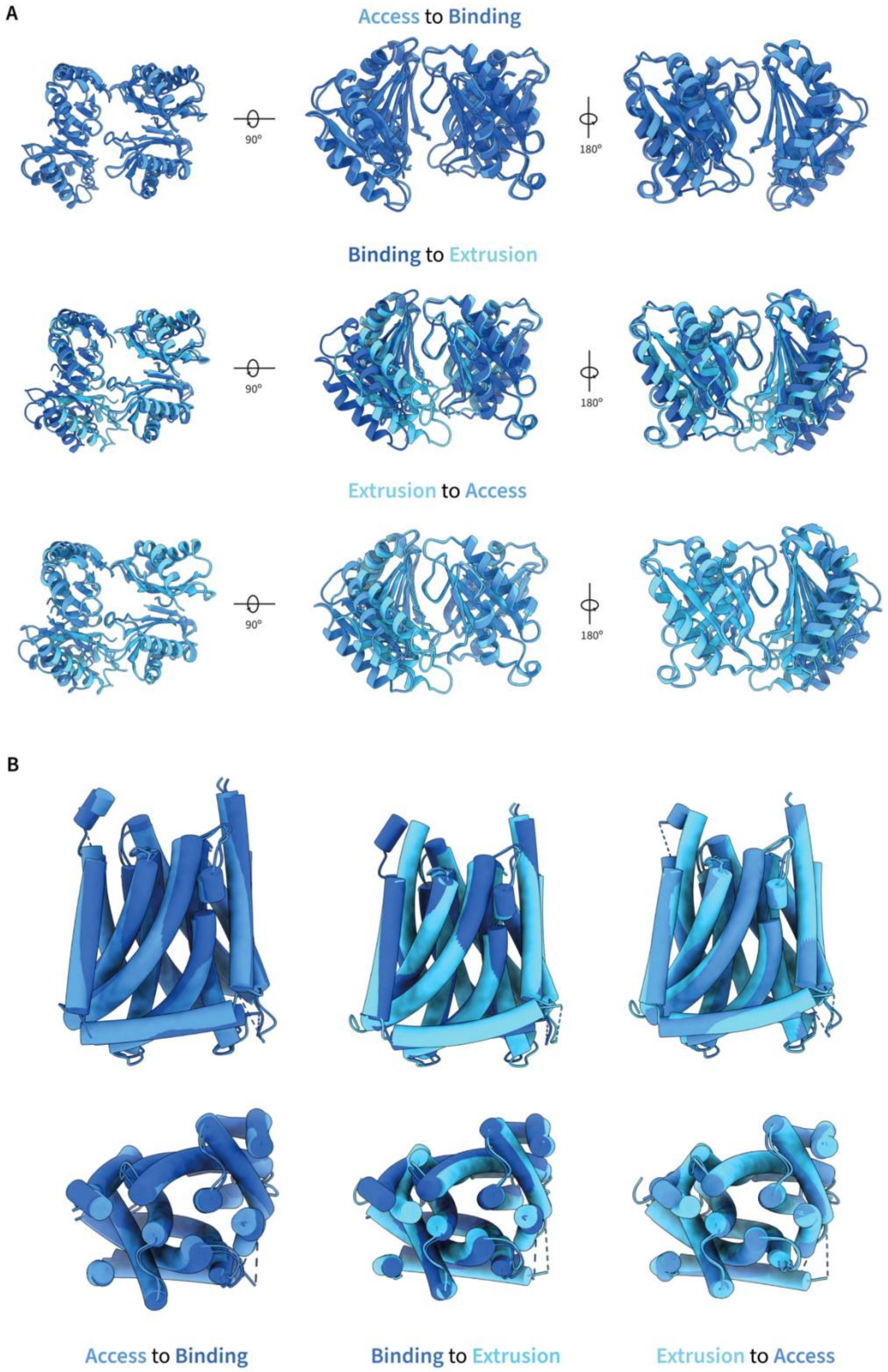


**Supplementary Fig. 15 | Analysis of conformational changes between states of MdtF$^{WT}$.** **a,** Conformational transitions observed in the porter domain of MdtF$^{WT}$ between access, binding, and extrusion states. **b,** Conformational transitions observed in the transmembrane domain in MdtF$^{WT}$ between access, binding, and extrusion states.

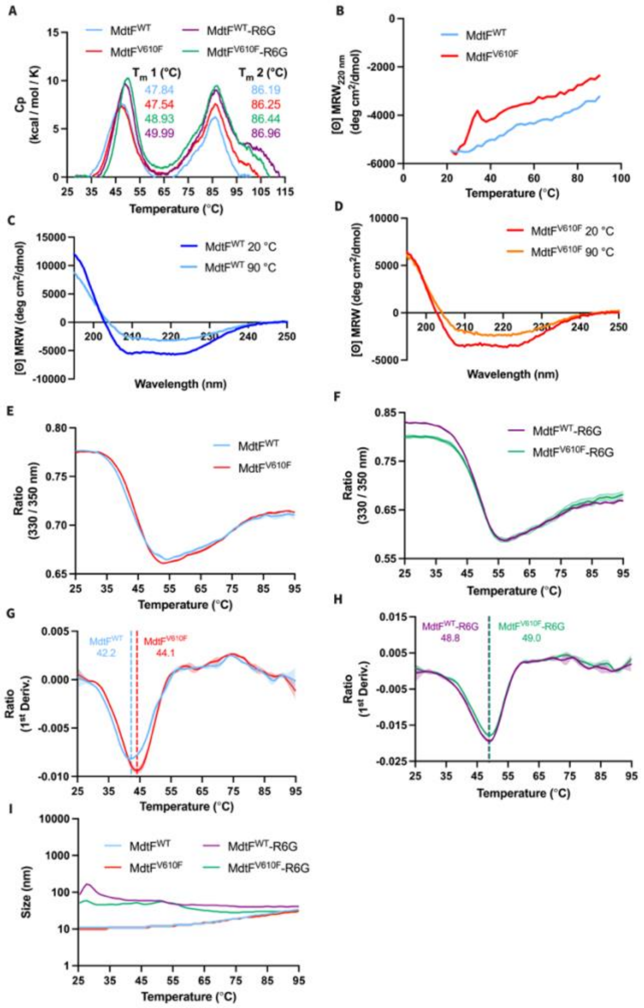


**Supplementary Fig. 16 | MdtF thermal stability. a,** DSC demonstrated two thermal transitions. The two melting temperatures for each protein (± R6G) are indicated on the graphs. The first peak likely corresponds to the periplasmic domain of the protein and undergoes a stabilising upon R6G binding; however, the second peak likely corresponds to the highly stable transmembrane helices which are maintained within the native nanodiscs and remain unaffected by R6G binding. **b,** To further explore this, we utilised CD experiments to demonstrate the secondary structure melting. Here, CD displayed the periplasmic protein domain transition in accordance with DSC, however, the spectra show the protein does not fully unfold up to 90 °C (**c and d**). Again, this suggests that the second transition corresponds to the stable, buried TM $\alpha$-helical bundle. **e-h,** Finally, the stabilising effect of R6G binding to both $MdtF^{WT}$ and $MdtF^{V610F}$ structure of ~5 °C was corroborated through monitoring the specific probing of the tertiary environment using DSF. **i,** Here, we also revealed that the self-assembled lipid nanodiscs remain intact during thermal MdtF unfolding which consolidates that this response is not due to possible artefacts arising from particle or oligomer disassembly. Collectively, this supports the thermal stability of MdtF within the SMALP nanodisc in addition to the stabilising effect arising from R6G binding. CD: Circular dichroism, Cp: Specific heat, DSC: Differential scanning calorimetry, DSF: differential scanning fluorimetry, MRW: Mean residue ellipticity, R6G: Rhodamine 6G, SMALP: Styrene maleic acid lipid particle, $T_m$: Melting temperature, TM: transmembrane helix.

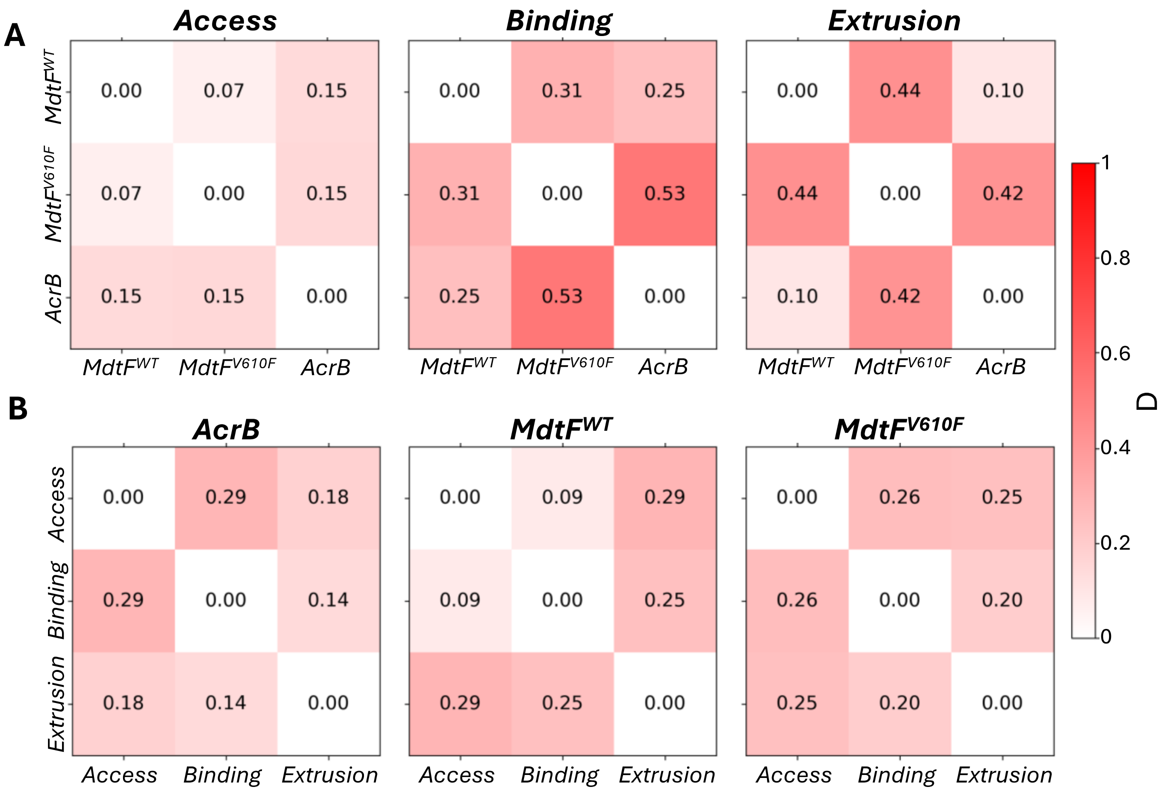


**Supplementary Fig. 17 | PC2-PC2 pseudo-contacts distribution analysis between monomers and proteins.** The matrices show the Kolmogorov-Smirnov (*Dnorm*) statistic (calculated using ks_2samp from SciPy), which quantifies the maximum difference between the empirical cumulative distribution functions (ECDFs) of PC2-PC2 pseudo-contacts. **(A)** comparison within LTO states of different proteins ($MdtF^{WT}$, $MdtF^{V610F}$, and AcrB), and **(B)** Comparison within each protein across different functional states (Access, binding, extrusion). The KS statistic is calculated as: $D = \max|F_1(x) - F_2(x)|$, where: $F_1(x)$ and $F_2(x)$ are the ECDFs of two monomer or proteins dataset. $D$ represents the KS statistic. The colour intensity represents the magnitude of differences in pseudo-contact distributions, with white ($D = 0$) indicating perfect overlap between the two distributions (similar) and red (high $D = 1$) indicating no overlap between the two distributions (dissimilar).

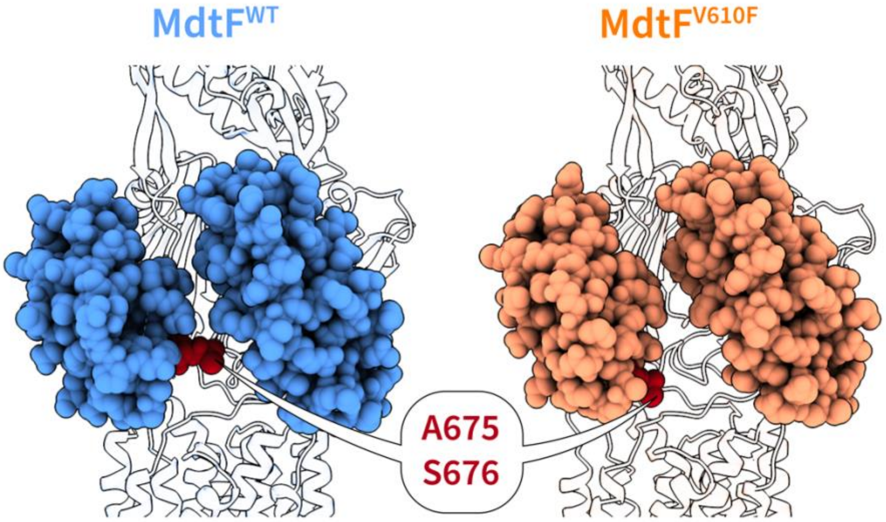


**Supplementary Fig. 18** | **PC1-PC2 subdomain cleft of MdtF$^{WT}$ and MdtF$^{V610F}$.** The PC1 and PC2 subdomains of MdtF$^{WT}$ and MdtF$^{V610F}$ displayed in atom representation (spherical form) in blue and orange, respectively. These 2 subdomains form the binding cleft at the front of the monomer where there is an extension of residues A675 and S676 in MdtF$^{WT}$, displayed in red, which is not conserved in MdtF$^{V610F}$.

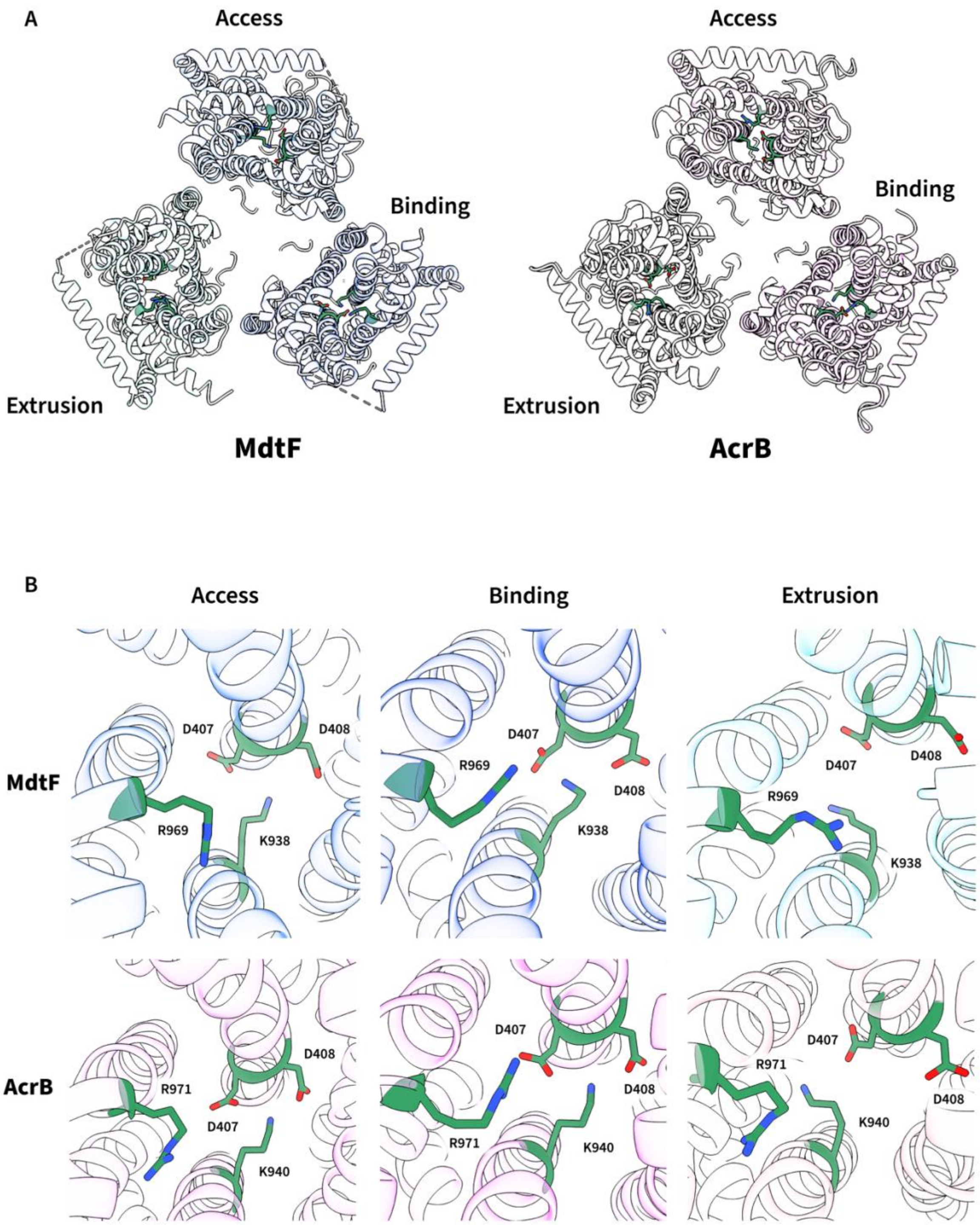


**Supplementary Fig. 19 | Conservation of proton relay network residues between AcrB$^{WT}$ and MdtF$^{WT}$. a,** The location of the proton relay network residues in each protomer as viewed from the cytosolic side, demonstrating its conservation. **b,** The proton relay residues D407, D408, K938, and R969 (K940 and R971, respectively, in AcrB) are conserved positionally in addition to their movements between each monomeric state.

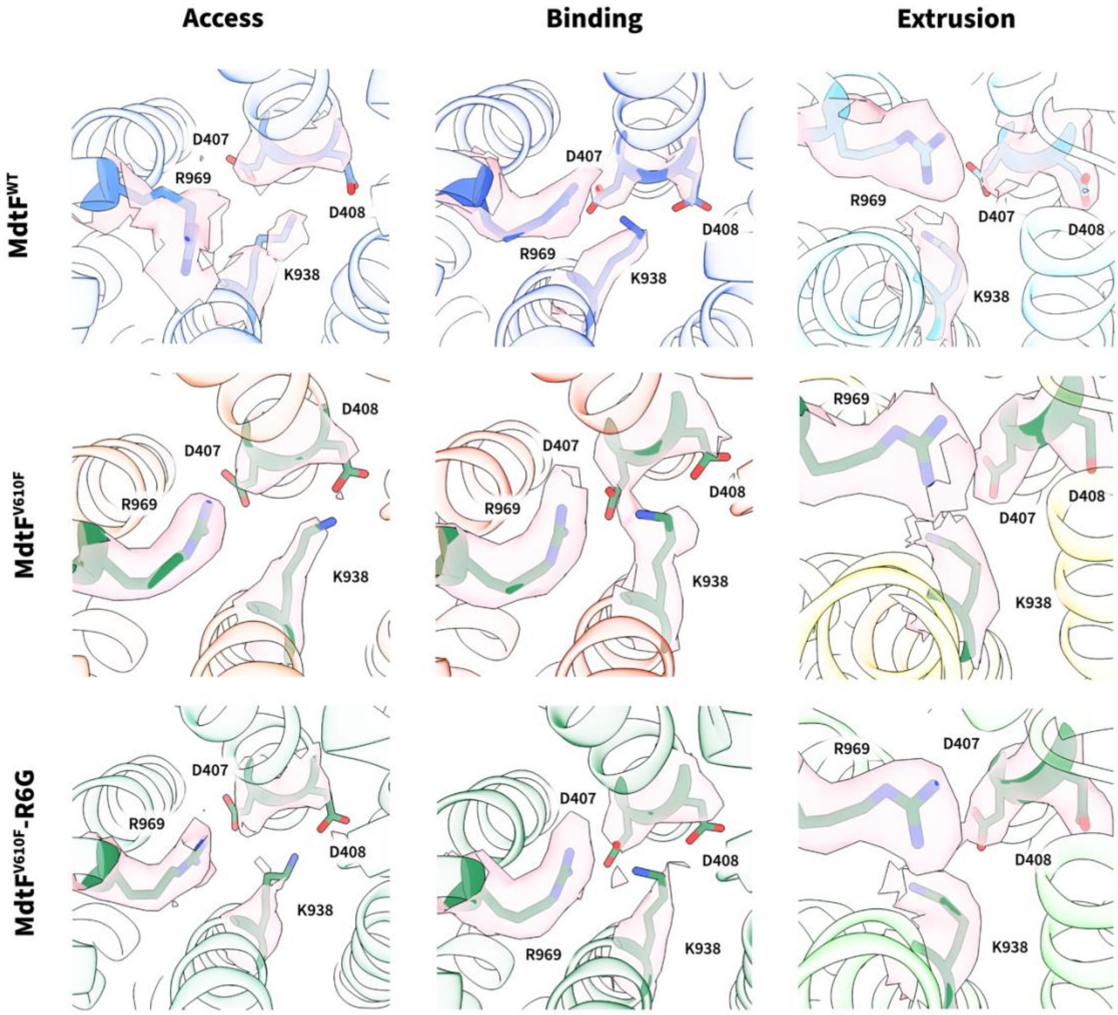


**Supplementary Fig. 20 | Conservation of proton relay network esidues across MdtF structures.** The proton relay residues D407, D408, K938, and R969 across MdtF$^{WT}$, MdtF$^{V610F}$, and MdtF$^{V610F}$-R6G and their respective electron density is shown. The residues and their respective conformational transitions between monomeric states are conserved, including the K938 transition. Cryo-EM density maps are represented in surface form (light pink). R6G: Rhodamine 6G.

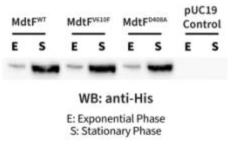


**Supplementary Fig. 21 | MdtF Nile Red efflux assay mutant expression test.** Expression test of MdtF mutants for the Nile Red efflux assay. Cell lysates of *E. coli* Δ9-Pore cells expressing MdtF mutants at exponential and stationary phase were separated by SDS-PAGE. Gels were transferred to a nitrocellulose membrane and subsequently immunoblotted with anti-His antibodies. WB: Western blot.

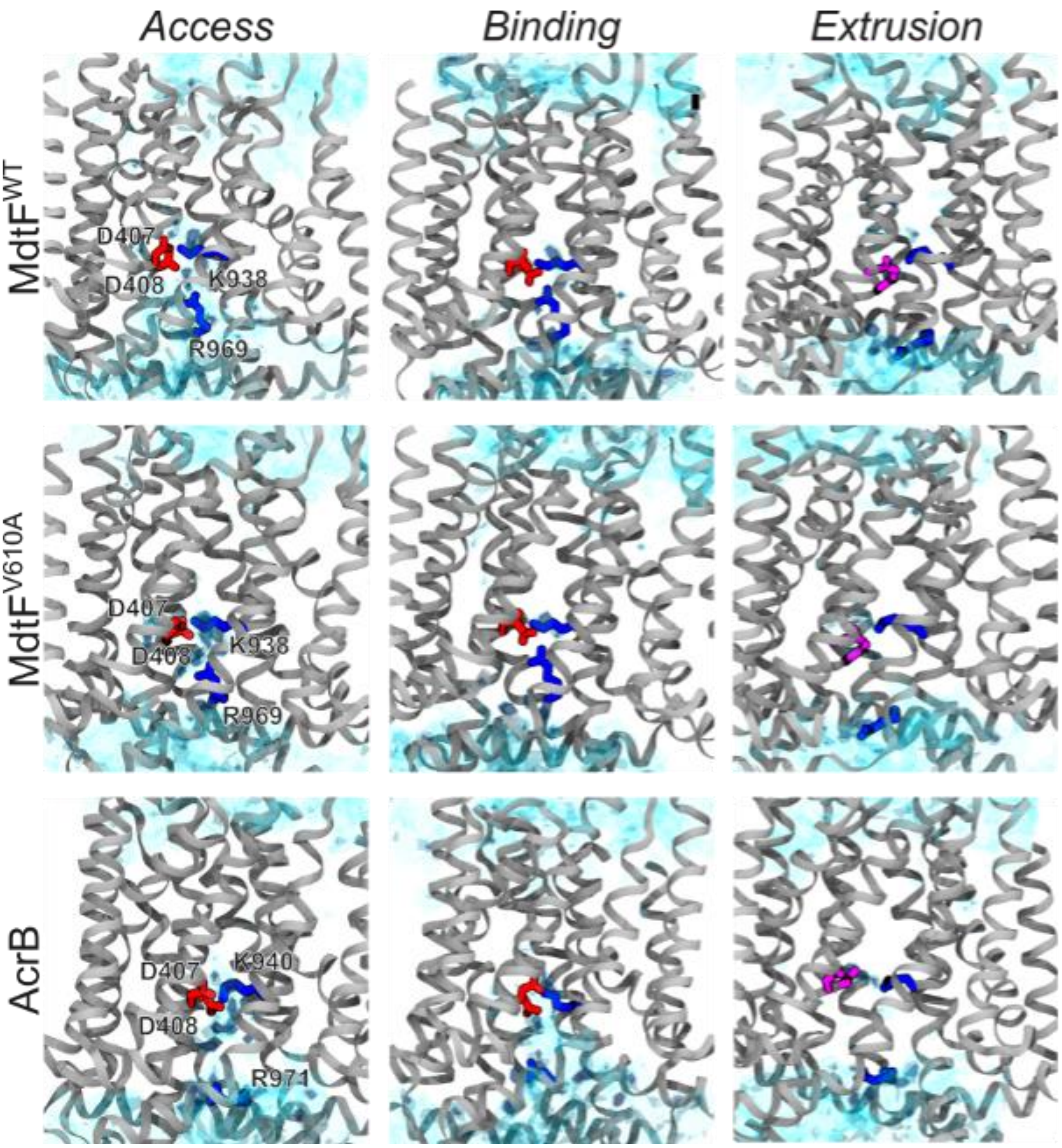


**Supplementary Fig. 22 | Hydration at the proton relay site.** Figure shows the preferred cumulative location of hydration sites (cyan surfaces) calculate along the MD simulations of the three systems discussed here. Surface transparency reflects water density, with higher density appearing more opaque and lower density more transparent. Residues D407 and D408 are shown in red liquorice for the L and T states and in pink for the extrusion state, to reflect their different preferred protonation state. K938 (K940) and R969 (R971) are depicted in blue liquorice for MdtF (AcrB). MD: Molecular dynamics.

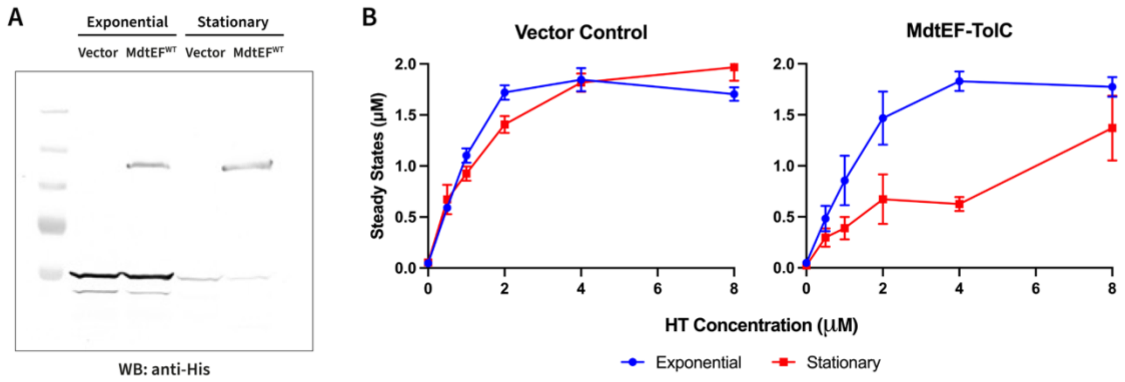


**Supplementary Fig. 23 | Outer membrane pore expression differs between exponential and stationary phases. a,** *E. coli* Δ9-pore cells, harbouring an empty vector or pUC19-MdtEF$^{WT}$ plasmid, were growth to exponential (4h of growth) and stationary (20 h of growth). phases of growth, pelleted, washed, and resuspended in HMG buffer (pH 7.0). Cell lysates were separated by SDS-PAGE. Gels were transferred to a PVDF membrane and subsequently immunoblotted with anti-His antibodies, followed by an anti-mouse alkaline phosphatase-conjugated secondary antibody. Although MdtEF$^{WT}$ expression was observed at both exponential and stationary phases, there was a lower expression of the outer membrane pore observed at stationary phases which is likely a consequence of arabinose metabolism within the *E. coli* Δ9-pore cells. **b,** Steady state accumulation levels of HT were measured in *E. coli* Δ9-pore cells, harbouring an empty vector or pUC19-MdtEF$^{WT}$ plasmid, as a function of external HT concentration. Cellular expression of an outer membrane pore was induced with 0.1 % L-arabinose. Cells were grown to exponential (4h of growth) and stationary (20 h of growth) phases of growth. Individual data points represent mean values from three independent measurements and error bars are indicative of the standard deviation (n = 3). Due to the differential expression of the outer membrane pore in the stationary phase, there is not convincing evidence that MdtF has a different efficiency in stationary phase non-growing cells. HT: Hoechst 33342, PVDF: Polyvinylidene fluoride, WB: Western blot.

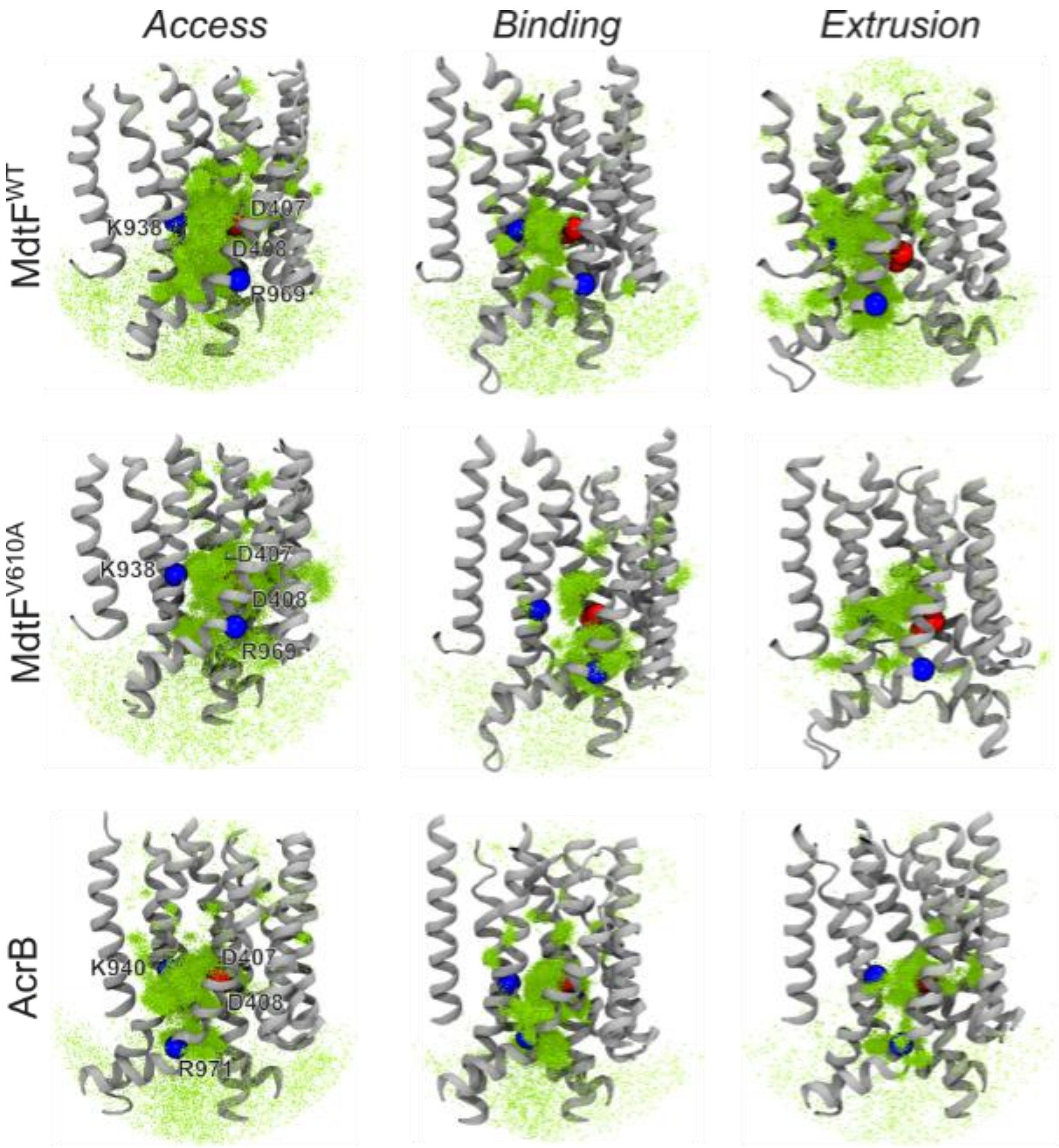


**Supplementary Fig. 24 | Water accessibility at the proton relay site.** Dynamic water exchange showing the locations from where water molecules are exchanged to the proton relay site (within 5 Å) in the transmembrane region. It shows the accessibility of water in AcrB and $MdtF^{WT}$ and $MdtF^{V610F}$.

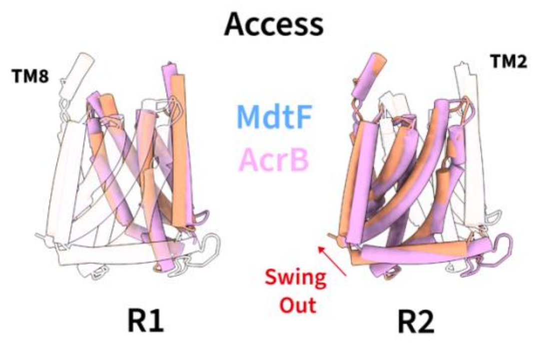


**Supplementary Fig. 25 | MdtF$^{V610F}$ exhibits MdtF$^{WT}$-like 'swung out' R2 state.** Alignment of AcrB (pink) and MdtF$^{V610F}$ (orange) transmembrane domains which reveals structural differences between its helical arrangement as it cycles through the protomeric states as observed within MdtF$^{WT}$ (**Fig. 2C**).

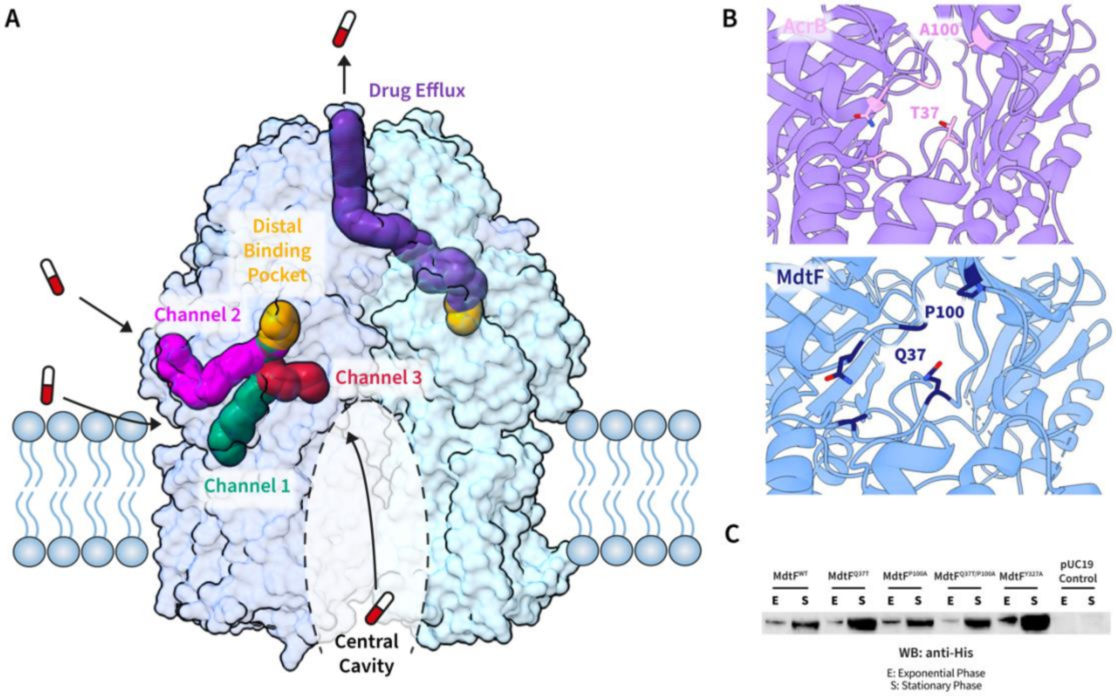


**Supplementary Fig. 26 | MdtF$^{WT}$ channels of export and channel 3 entrance. a,** The RND-related channels of MdtF$^{WT}$ as calculated by MOLE[8]. **b,** The residues gating the channel entrance exhibit variations between MdtF$^{WT}$ (Q37 and P100, blue) and AcrB (T37 and A100, pink). **c,** MdtF CH3 mutant expression test. Here, cell lysates of *E. coli* Δ9-Pore cells expressing MdtF mutants at exponential and stationary phase were separated by SDS-PAGE. Gels were transferred to a nitrocellulose membrane and subsequently immunoblotted with anti-His antibodies. CH3: Channel 3, WB: Western blot.

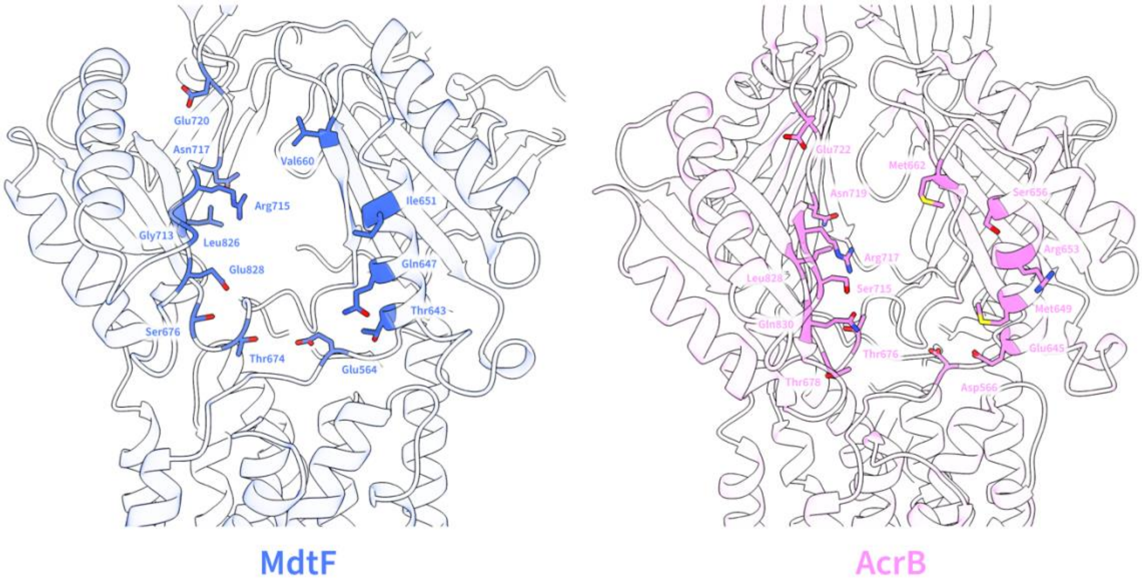


**Supplementary Fig. 27 | Residues observed at CH2 entrance of MdtF$^{WT}$ and AcrB$^{WT}$.** CH2 residues of MdtF$^{WT}$ (blue) and AcrB (pink) are displayed in stick representation. Here, key residues in MdtF (F662, R715, T674, and M650) differ from those in AcrB which may impact binding site interaction due to their inward orientation to the channel entrance. CH2: Channel 2.

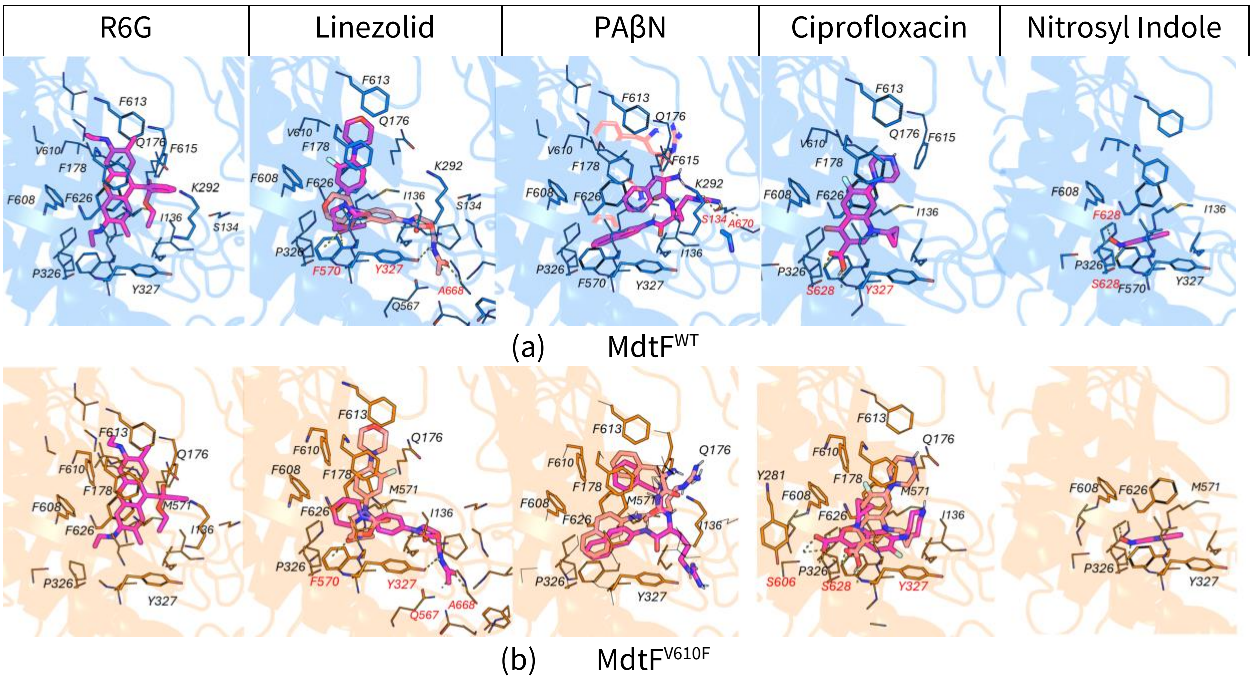


**Supplementary Fig. 28 | Docking poses in the DBP of MdtF.** Key residues (side chains) are shown in line representation, while ligands are depicted in stick representation. When an alternative pose with comparable binding affinity is identified, the second-best ranked pose is also displayed. The best-ranked ligand pose is coloured dark pink, while the alternative pose is shown in light pink. Hydrogen bonding interactions are represented by dotted lines and labelled in red. The protein is rendered as a transparent cartoon, with MdtF$^{WT}$ and MdtF$^{V160F}$ structures shown in blue and gold, respectively. Panels **(a)** and **(b)** are the complexes for MdtF$^{WT}$ and MdtF$^{V610F}$, respectively. DBP: Distal binding pocket, PAβN (phenylalanine-arginine β-naphthylamide), R6G: Rhodamine 6G.

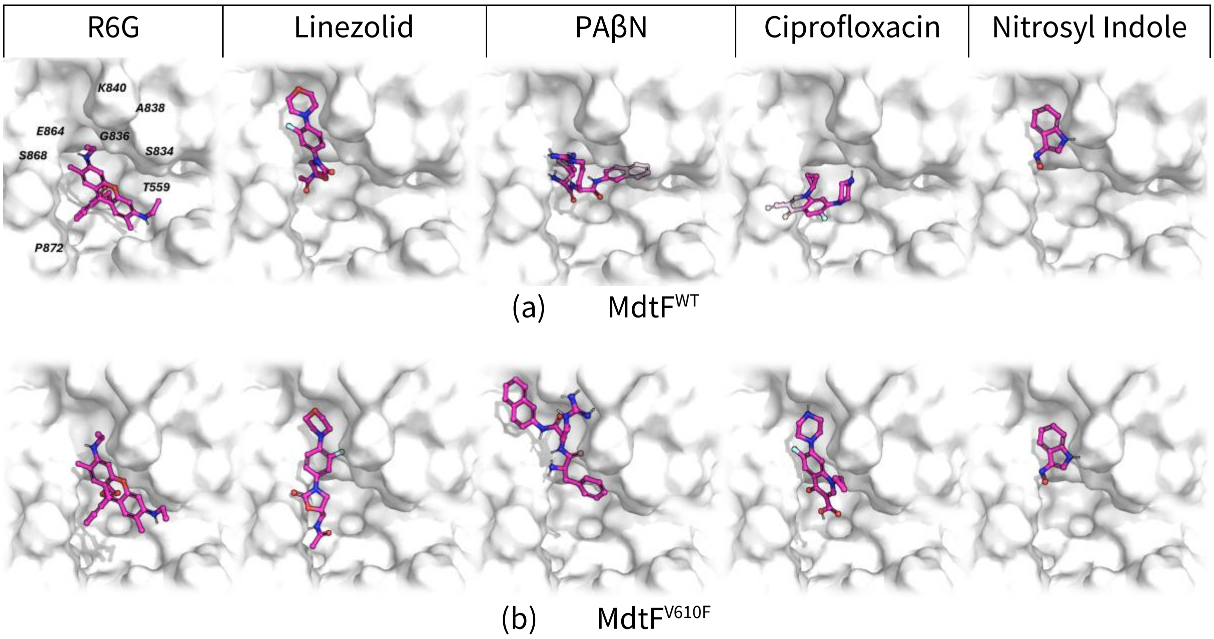


**Supplementary Fig. 29 | Docking poses at the CH1 entrance of MdtF$^{WT}$ and MdtF$^{V610F}$.** Protein is represented as white transparent surface for clarity. CH1: Channel 1, PAβN (phenylalanine-arginine β-naphthylamide), R6G: Rhodamine 6G.

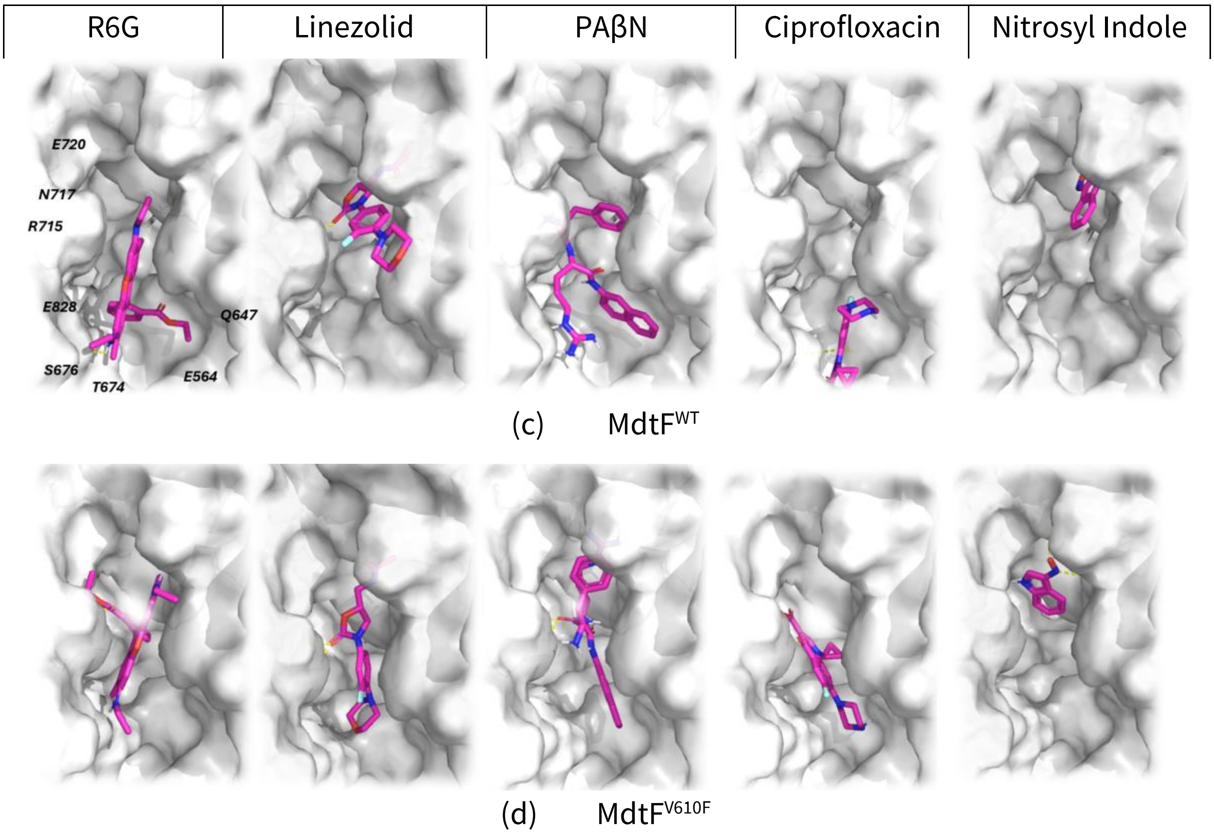


**Supplementary Fig. 30 | Docking poses at the CH2 entrance of MdtF$^{WT}$ and MdtF$^{V610F}$.** Protein is represented as white transparent surface for clarity. CH2: Channel 2, PAβN (phenylalanine-arginine β-naphthylamide), R6G: Rhodamine 6G.

| R6G | Linezolid | PAβN | Ciprofloxacin | Nitrosyl Indole |
|---|---|---|---|---|

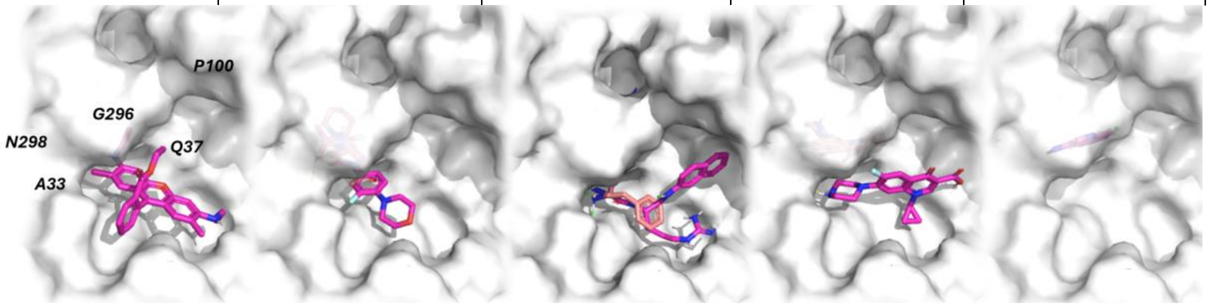


(a) $MdtF^{WT}$

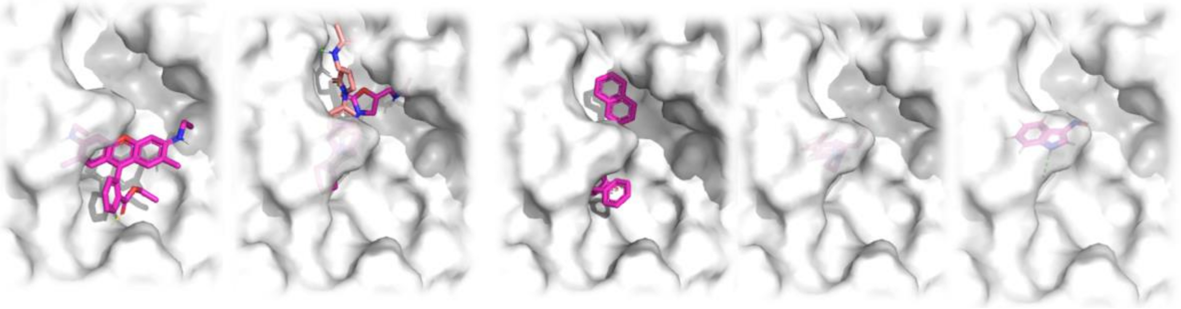

(b) $MdtF^{V610F}$

**Supplementary Fig. 31 | Docking poses at the CH3 entrance of $MdtF^{WT}$ and $MdtF^{V610F}$.** Protein is represented as white transparent surface for clarity. When an alternative pose with similar binding affinity is found, it is shown in light pink alongside the top-ranked pose in dark pink. CH3: Channel 3, PAβN (phenylalanine-arginine β-naphthylamide), R6G: Rhodamine 6G.
